# Cellularly-Retained Fluorogenic Probes for Sensitive Cell-Resolved Bioactivity Imaging

**DOI:** 10.1101/2025.04.17.649302

**Authors:** Philipp Mauker, Lucas Dessen-Weissenhorn, Carmen Zecha, Nynke A. Vepřek, Julia I. Brandmeier, Daniela Beckmann, Annabel Kitowski, Tobias Kernmayr, Julia Thorn-Seshold, Martin Kerschensteiner, Oliver Thorn-Seshold

**Affiliations:** Faculty of Chemistry and Food Chemistry, Dresden University of Technology, Bergstrasse 66, Dresden 01069 (Germany); Department of Pharmacy, Ludwig-Maximilians University of Munich, Butenandtstrasse 7, Munich 81377(Germany); Institute of Clinical Neuroimmunology, LMU University Hospital, Ludwig-Maximilians University of Munich, Marchioninistrasse 15, Munich 81377 (Germany); Biomedical Center (BMC), Faculty of Medicine Ludwig-Maximilians University of Munich, Grosshaderner Strasse 9, Martinsried 82152 (Germany); Faculty of Medicine, University Hospital Dresden Dresden University of Technology, Fiedlerstrasse 42, Dresden 01307 (Germany); Munich Cluster for Systems Neurology (SyNergy), Feodor-Lynen-Strasse 17, Munich 81377 (Germany)

**Keywords:** Xeniidae, Barcoding, Disturbance, Caribbean, pulsing corals

## Abstract

Here, we develop a general design for high-quality fluorogenic activity probes to quantify biological processes in live cells, by creating a scaffold that efficiently generates cell-retained bright fluorescent soluble products upon reaction with biochemical targets. Live cell probes must be designed to be membrane-permeable; but that often means that their fluorophore products are similarly permeable, resulting in rapid signal loss from the activating cell: which limits their cell-by-cell resolution as well as their sensitivity for quantifying low-turnover processes. Current strategies to retain fluorescent products within cells usually disrupt native biology: e.g. by non-specific alkylation or solid precipitation. Here, scanning charge- and polarity-based approaches to trigger cell retention, we developed a bright fluorogenic rhodol scaffold **<u>Tra</u>ppable <u>G</u>reen** (**TraG**) that balances all key requirements for signal integration (rapid probe entry, but effective product retention, across a variety of cell lines) and is easily adaptable to quantify many target types (shown here with probes for GSH, TrxR, and H_2_O_2_). The simple and rugged **TraG** scaffold can now permit straightforward elaboration to a range of cell-retained enzyme activity probes, that enable more accurate cell-resolved imaging as well as higher-sensitivity integration of low-turnover processes, without the drawbacks of alkylation or precipitation-based strategies.

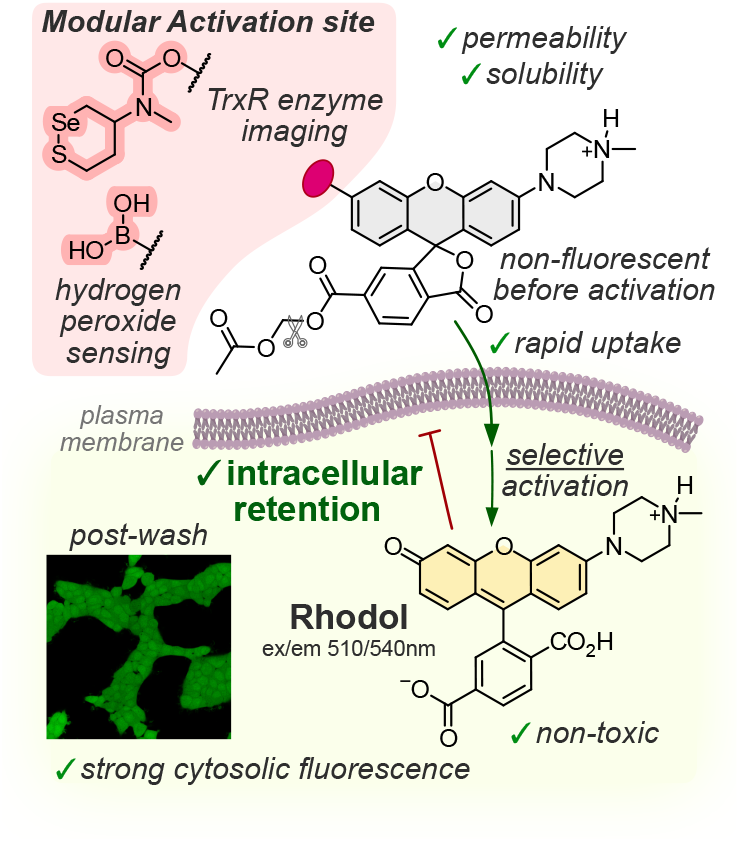

## Introduction

Imaging and quantifying biological activity are key challenges in basic and applied research. Focusing on enzyme activity rather than mRNA or protein levels takes post-translational regulation mechanisms into account (PTMs, compartmentalisation, chaperoning), and allows researchers to correctly interpret biochemistry in action. Fluorogenic probes are, ideally, non-fluorescent probes that only generate a fluorescent product after activation by their specific biochemical target or enzyme. They have become crucial tools for sensitive and non-invasive bioactivity imaging, especially in lysates: with probes for peptidases, esterases, phosphatases, glycosidases, and oxidoreductases,^[1–7]^ or reactive analytes such as hydrogen peroxide and hydrogen sulfide,^[8,9]^ in widespread use.

Sensitive, cell-resolved detection is crucial for longitudinally visualising bioactivity during assay time courses, and for understanding the heterogeneity of cell populations. However, fluorogenic probes often encounter a major problem in live cells and tissues: that the signal of their fluorescent products becomes diffuse or is lost over time. Apolar, membrane-permeable fluorophores such as coumarins can rapidly exit the cell across the plasma membrane by passive diffusion;^[10]^ while negatively charged fluorophores such as fluoresceins are instead excreted from cells by active transport.^[11,12]^ This *post-activation signal loss* sabotages cell-resolved activity imaging, and lowers the sensitivity and reliability of signal quantification (higher and time-dependent background signal); and the *rate* of signal loss sets a lower limit on the enzyme activity or analyte concentration that can even be detected at all. For *in vivo* work requiring low probe concentrations, or for imaging low-turnover processes, or for situations demanding high sensitivity and quantitative reliability, building up and retaining the product signal inside the activating cell in the long term is a crucial challenge for probe design.

The importance of retaining fluorescence signal within the cell has driven three probe designs for intracellular signal trapping (full discussion at **Fig S3**). (1) *Charge/polarity-based product impermeabilization* usually suppresses its passive membrane transit with ionic motifs e.g. carboxylates^[13]^, phosphonates,^[14]^ sulfonates,^[15]^ or tetraalkyl-ammoniums^[16,17]^. To deliver probes into the cell in the first place, intracellularly-cleaved lipophilic masking groups^[13–15,18]^ (e.g. masking carboxylates as acetoxymethyl esters^[9,13,19,20]^ or amines as carbamates^[21]^), endocytosis (e.g. by cell-penetrating peptides),^[22]^ or transporter-mediated uptake,^[16]^ are commonly used. None of these approaches is *generally applicable*, however, mainly since the activation triggers are not modular; product fluorophore brightness is low; cellular uptake is slow; reactive side products are released; or unwanted compartmentalisation occurs. (2) *Water-insoluble solid-state fluorophores* can be released as reaction products, that precipitate as fluorescent cellularly-trapped crystals (e.g. probe ELF-97 that releases the fluorophore HPQ).^[23–25]^ Yet, crystal deposits cause strong inflammatory responses and cytotoxicity, perturbing biology or preventing longitudinal imaging; and these probes are unsensitive at low turnover since the precipitation threshold must be crossed before any signal is seen; also, these probes rarely have good solubility (mirroring their product insolubility). (3) Releasing *products that alkylate cell-impermeable biomolecules* was pioneered as a cell retention strategy by Urano. SPiDER probes featuring non-reactive benzylfluorides which are enzymatically triggered to generate electrophilic quinone methides that rapidly react with proteins and GSH, enabling long-term signal retention.^[6,26–29]^ However, they can also alkylate their target enzyme,^[29]^ induce electrophile stress responses, or accumulate toxic effects,^[27]^ especially in high-turnover cells.

We identify eight requirements for an ideal probe to allow high-sensitivity enzyme imaging in live cells (**Fig 1**): (a) good aqueous solubility to ensure reproducible handling and high bioavailability, and to avoid probe aggregation or sequestration; (b) high robustness, i.e. no non-specific background product release; (c) reliably effective cell entry in different cell lines (e.g. by passive diffusion); (d) linear fluorescence signal (e.g. the target activates fluorescence by just one reaction site per probe molecule); (e) high signal turn-on ratio (i.e. probe is very dark under typical imaging conditions); (f) bright product fluorescence; (g) effective cellular retention of the product, allowing long-term signal integration; (h) lack of toxicity for the fluorophore and any probe byproducts, so biology is unperturbed in the long term (unlike precipitating or alkylating probes). Ideally, the probe design would also be *modular*, i.e. easily chemically adaptable to image various target types. Considering that none of the prior strategies meets all these requirements (**Fig S3**), we set out to develop a probe design that does. We chose *O*- unmasking of a phenolic fluorophore for activation: a reaction that is applicable for many types of molecular imaging. We now outline the development and demonstration of a generalised *O*- masked rhodol probe design for high-sensitivity enzyme imaging with a cell-retained product, that meets all these requirements.

**Figure 1.**
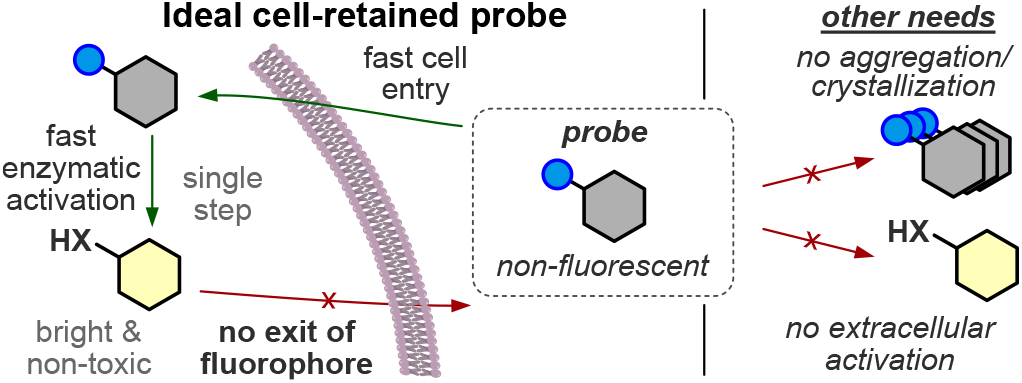
Requirements for an ideal, sensitive, cell-retained fluoro-genic probe.

## Results and Discussion

### Approach 1: Lipidated Charged Fluoresceins

Facing an eight-factor optimisation problem, we first focused on using synthetically accessible *O-*masked fluorogenic probe designs to test what physicochemical features would ensure good cell entry and cell retention and left the other requirements for later. Fluorescein spirolactone *O*-alkylated *O’*-esters are a convenient fluorogenic test system with only one reaction site;^[30]^ and we had previously noted that some monosulfonated fluorescein diesters were surprisingly capable of cell entry despite their charge.^[31]^ We now took these known systems and measured their cell retention after washing, with standard conditions: which was promising for the diester (**i**_**2**_**-FS, Fig S4**) though poor for the monoester (**iPS-F**). Following the notion that medium-length lipids enhance the cell uptake and retention of natural products,^[32]^ we next synthesised a set of more lipid-like *O*-alkylated sulfonated fluorescein monoester probes (**iC4-FS** — **iC10-FS**, for C4—C10 alkyl; all structures: see **Fig S1**). All lipidated designs gave good product retention after washing (**Fig S5bd**), but uptake was poor (**Fig S5c**; their cellular signal distribution also varied from uniform **C4-FS** to membrane-only **C10-FS**; **Fig S5e**). Deleting the sulfonate to raise permeability was a failure (**iC4-F**: no signal seen in cells (**Fig S6e**) or in cell-free esterase assays (**Fig S24**)), illustrating the importance of solubility for bioavailability. Using a reversibly ionisable carboxylate in **iC4-F<u>C</u>** (pK_a_ ≈ 4.3) instead of the sulfonate gave 20-fold higher cellular signal intensity than **iC4-FS**, but kept its good post-wash retention and uniform signal distribution (**Fig S6**). However, we had noted significant cellular distress (rounding and blebbing) with all FS- and FC-ester probes so far. By switching the trigger group from an *O-*isobutyrate ester to a more hydrolytically robust, reductively cleavable *O*-carbamate (**GL-C4-FC**; **Figs S23, S25**), we discovered that the pairing of strong signal with cell blebbing was caused by the *combination* of the lipidated FS/FC probes with the salt buffer solution that had been needed to avoid isobutyrate ester hydrolysis. In more complete buffer (DMEM), cell morphology stayed healthy, and no signal was seen (**Figs S6f, S7**). We imagine that the membrane stress of the amphipathic FS/FC probes, plus the lack of nutrients in salt buffer, was causing enough membrane disruption for the charged probes to enter cells^[33]^ (discussion at **Fig S6**): indicating that amphipathic probes are unsuitable for non-invasive cell imaging.

### Approach 2: Charge-Balanced Rhodol Probes

To reach good cell entry and good retention, we expected that we would need additional charges compared to the FS/FC designs. We also prioritised good probe solubility, used the hydrolytically stable *O-*carbamates for systematic studies in complete cell culture media, and switched to rhodol scaffolds, since rhodols can be *O-*acylated to give nonfluorescent (fully spirolactone) probes, but can be much brighter than *O*’-alkyl fluoresceins (limit: ∼7×10^3^ L mol^−1^ cm^−1^, **Table S1**), and their fluorescence is more photostable and is constant over the pH range 4 - 10.^[35]^ We then synthesised a set of six reductively-activated^[34]^ “GL-Rho” probes: with (1) either an apolar piperidine (**Rho**) or a basic piper**a**zine (**Rho-A**) as *N-*substituent; and (2) optionally, a 6-carboxylate (**C**) or its masked, membrane-permeable AM ester (**C**^**m**^) (**Fig 2a**). The rhodol fluorophores were accessed from the fluorescein ditriflates by one-sided Buchwald-Hartwig coupling, then triflate hydrolysis with LiOH or TBAF. A one-pot sequence then transformed the phenols into pentafluorophenyl carbonates before using them to acylate the GSH-labile **GL** disulfide motif. Where relevant, the 6-carboxylates were then deprotected, and optionally masked, to give the probes.

**Figure 2.**
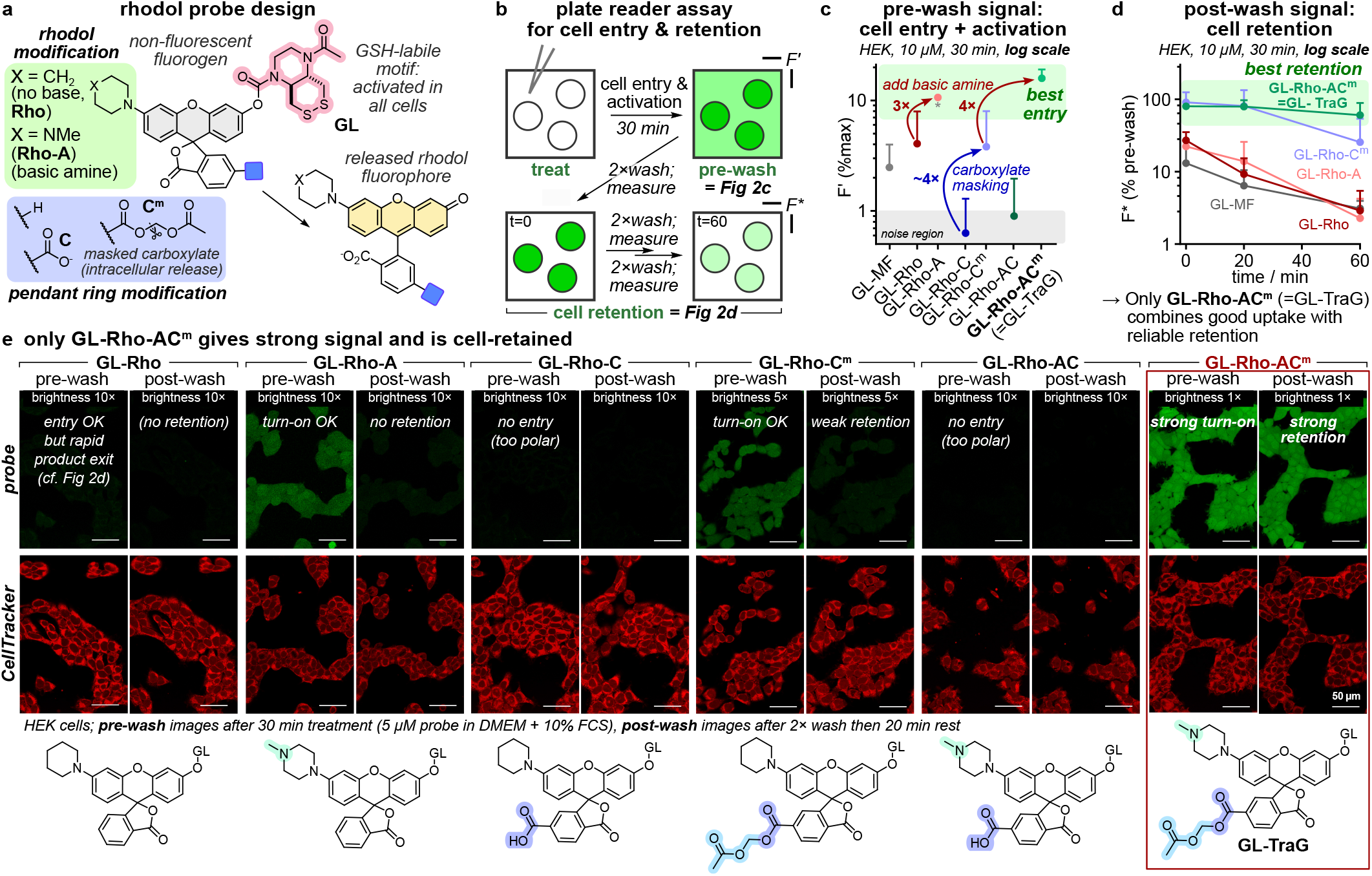
The rhodol probe design Rho-AC^m^ (= Trappable Green: TraG) delivers strong intracellular signal activation and retention. **(a)** Probe panel overview. **(b)** Plate reader assay for cell entry, activation, and retention. **(c)** Entry and activation in HEK293T cells (10 µM probe, DMEM with 10% FCS, 30 min treatment; F’ is fluorescence as % of full activation of whole well; error bars: SD; biological replicates normalised to **GL-Rho-A** (marked with asterisk); n=3; error bars: SD; the methyl fluorescein probe **GL-MF**^[34]^ serves as a non-rhodol scaffold entry and cell retention benchmark, see **Fig S1**). **(d)** Cell retention (conditions as in **c**; F* is fluorescence as % of pre-wash value; error bars = SD). **(e)** Cell turn-on and retention assayed by confocal microscopy (conditions as in **c** except 5 µM probe; scale bars: 50 µm; full data in **Fig S9a**).

The optical properties of the rhodol products varied somewhat, with excitation maxima at 490 - 530 nm, Stokes shifts of ∼25 nm (emission maxima 515 - 560 nm), and extinction coefficients of 30−60×10^3^ L mol^−1^ cm^−1^. Fluorescence quantum yields varied from 4 − 64%, with the piperazinyl probes **H-Rho-A** and **H-Rho-AC** being brightest (∼30×10^3^ L mol^−1^ cm^−1^; **Fig S20, Table S1**). Importantly for uses in high-sensitivity imaging, all probes were non-fluorescent, with outstanding probe/product signal turn-on ratios of up to ∼550 (**Fig S20**); the piperazinyl probes were particularly efficiently activated by their target GSH (**Fig S25**); and all probes were hydrolytically stable for hours in FCS-supplemented DMEM, for long term cell experimentation (**Fig S23**).

We then tested cell entry, activation, and signal retention, in HEK cells (**Fig 2b**). Cell entry and activation was moderate for piperidinyl **GL-Rho**, but 3× higher for the basic piperazinyl (**GL-Rho-A**; **Fig 2c**). Adding a carboxylate blocked cellular uptake (**GL-Rho-C** / **GL-Rho-AC**); masking it as an acetoxymethyl ester (**GL-Rho-C**^**m**^ / **GL-Rho-AC**^**m**^) returned to the same signal as **GL-Rho** / **GL-Rho-A**. Importantly, probe treatment does not impair cell morphology, and uptake occurs homogeneously in healthy cells with uniform cellular distribution of the product signal (**Fig S9a**,**10**). After uptake, we then subjected cells to three cycles of “wash, measure, and wait”, to monitor intracellular signal retention. The rhodol product from **GL-Rho** / **GL-Rho-A** leaked out rapidly from cells; but an added carboxylate (**GL-Rho-C**^**m**^ / **GL-Rho-AC**^**m**^) greatly enhanced cell retention, particularly for **GL-Rho-AC**^**m**^ where cells stayed bright despite three medium exchanges over 1 h (**Fig 2d**). We used confocal microscopy to complement these plate reader assays. **GL-Rho-A, GL-Rho-C**^**m**^, and **GL-Rho-AC**^**m**^ indeed show cell entry and activation, while **GL-Rho-C** and **GL-Rho-AC** do not (**Fig 2e**). Only **GL-Rho-AC**^**m**^ had strong post-wash signal retention (**Figs 2e, S9ab**) and gave good performance in HeLa, MEF, and A549 cell lines (**Fig S11**, strong retention in HeLa cells, weaker in A549 cells).

Thus, the **Rho-AC**^**m**^ scaffold’s combination of a basic amine with a masked, intracellularly-revealed carboxylate became our preferred design for cell-retained fluorogenic rhodol probes. Its combination of high fluorogenicity with good cell retention allows sensitive cell-resolved imaging either without washing, or with washing (even after a significant delay). Its aqueous solubility avoids aggregation effects; its full spirocyclisation plus its biochemical robustness allow zero-background imaging; and it rapidly enters different cell lines where it is efficiently activated to give a uniform signal. Crucially, neither the probe nor the fluorophore causes apparent cellular harm, supporting that the data acquired during longitudinal imaging or enzyme activity integration can be reliably interpreted. Since the **Rho-AC**^**m**^ scaffold shows best performance, we renamed it <u>Tra</u>ppable <u>G</u>reen (**TraG**) and used this modular scaffold for activity probes.

### Extension to Hydrogen Peroxide Sensing

Hydrogen peroxide (H_2_O_2_) is a major physiological messenger with baseline levels that fuel cell signalling and metabolic function,^[36]^ but which can also be created as an unwanted metabolic byproduct at harmful levels that have been correlated to neurodegeneration, cancer, or autoimmune disorders.^[37–39]^ Sensitive and linearly-responsive tools are needed to resolve and study its multiple roles. The most common small molecule probes for sensing H_2_O_2_ exposure use arylboronic acids that H_2_O_2_ converts to phenols.^[8]^ Signal integration is crucial for sensitively detecting low H_2_O_2_ concentrations, so cell retained probes (e.g. SPiDER: intracellular quinone-methide trapping) have been utilised despite their moderate cell-toxicity.^[19,27,40]^

We hoped that a **TraG**-based design could deliver a more biocompatible cell-retained H_2_O_2_ sensor (**Fig 3a**). As the common pinacol boronate diester was somewhat hydrolytically unstable during purification, we applied the probe as a free boronic acid (membrane permeable, pK_a_ ≈ 8−9^[41]^). This probe **HP-TraG** gave linear signal generation with H_2_O_2_ (**Fig 3b**), with up to 48-fold turn-on (**Fig S21**). Loading it into HEK cells (15 min), then washing and extracellularly administering 25−100 µM H_2_O_2_, gave H_2_O_2_-dependent intracellular fluorescence signals with high turn-on index (up to 7-fold increase, **Fig 3cd**), that were cell-retained for >2 h after washing off the extracellular medium (**Fig 3ef**). Finally, we used **HP-TraG** for imaging endogenous H_2_O_2_ in Hoxb8-derived macrophages^[42]^ after activation with phorbol 12-myristate 13-acetate (PMA).^[43]^ **HP-TraG** signal increases by 60% upon PMA treatment, i.e. sensitively detecting both the low endogenous baseline and the slightly increased H_2_O_2_ concentrations upon activation (1-4 µM in macrophages^[44]^), again with strong post-wash signal retention (**Figs 3g, S14**). Thus, the **TraG** cell-retained fluorogenic probe design adds a useful new hydrogen peroxide sensor to the toolbox of chemical biology, that gives strong performance (rapid, H_2_O_2_-dependent intracellular signal) while overcoming the drawbacks of cell-reactive quinone methides as trapping agents.

**Figure 3.**
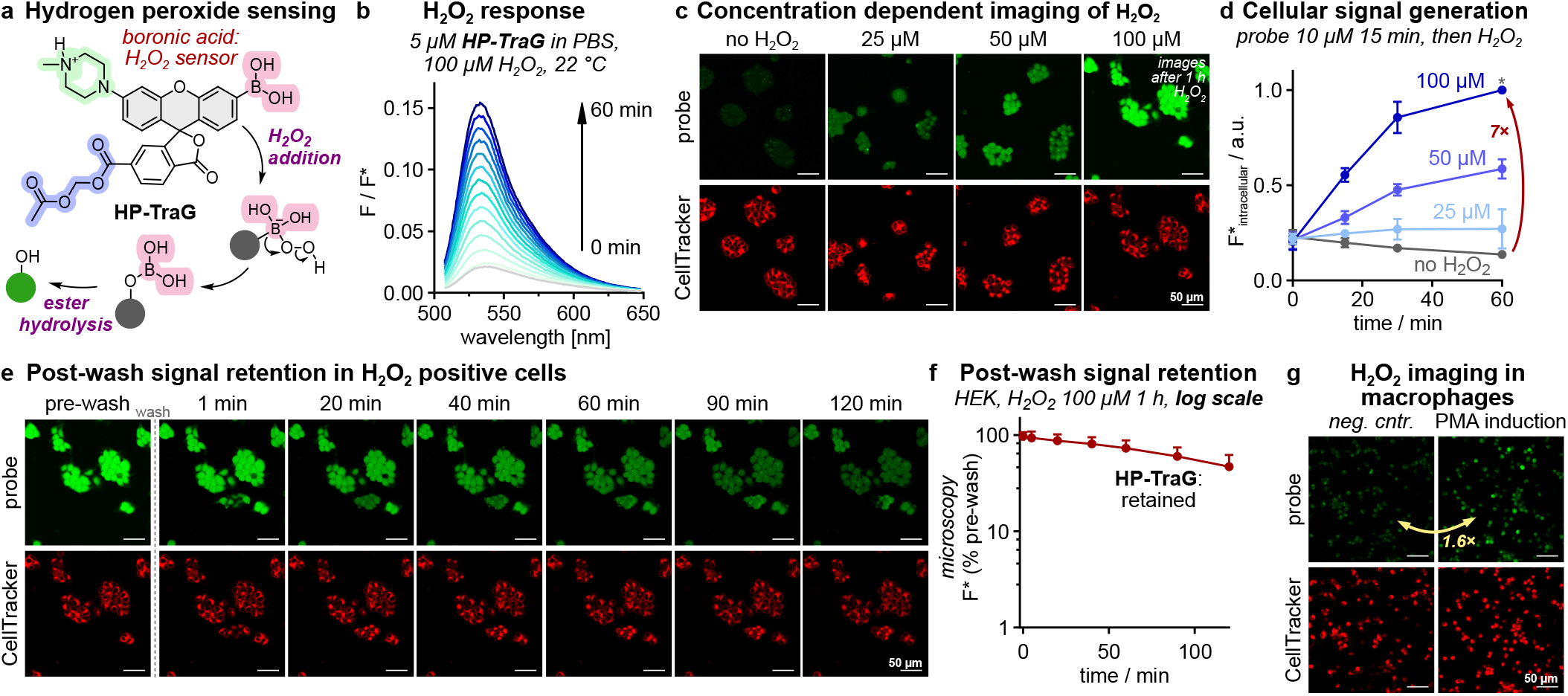
H_2_O_2_ sensing with HP-TraG. **(a)** Structure and mechanism of **HP-TraG. (b)** Cell-free H_2_O_2_ response (5 µM in PBS, 100 µM H_2_O_2_ over 60 min). **(c-d)** [H_2_O_2_]-dependent activation in HEK cells (15 min **HP-TraG** loading (10 µM), wash, then 60 min H_2_O_2_ treatment; full images in **Fig S12**; panel **d**: intracellular signal quantified from microscopy; *biological replicates benchmarked to 100 µM value at 1 h; *n*=3; error bars: SD). **(e-f)** Post-wash intracellular signal retention (HEK cells treated as in **c-d**, with 100 µM H_2_O_2_ for 1 h then washed and imaged; full images in **Fig S13**; *n*=3; error bars: SD). **(g)** Hoxb8-derived macrophages loaded with **HP-TraG** (10 µM for 15 min) then treated with phorbol 12-myristate 13-acetate (PMA, 1.6 µM for 1 h) then imaged (ratio quantified from images; full images in **Fig S14**). (All scale bars: 50 µm).

### Extension to TrxR Enzyme Imaging

Mammalian thioredoxin reductase (TrxR) is a key enzyme that uses NADPH to reduce thioredoxins, which drive hundreds of redox reactions involved in metabolism, protein folding, and signaling.^[45,46]^ TrxR is also one of just 25 selenoproteins in the human proteome; its selenium is needed so that its activity is resilient against biochemical damage.^[47]^ TrxR is a difficult target for molecular imaging due to its low expression level (ca. ≤20 nM) vs. high levels of chemically similar thiol off-targets (>10 µM).^[48]^ Only activity imaging can map TrxR function, since its activity is decoupled from mRNA levels (Se incorporation is regulated post-transcriptionally), and antibodies do not distinguish non-functional or non-Se forms. The first and only TrxR-selective probe for live cell activity imaging, RX1, was published in 2022,^[49]^ and is used for redox biology studies and high-throughput screens.^[50]^ RX1’s target specificity stems from its cyclic selenenylsulfide, a substrate that is selectively reduced by TrxR then cyclises to release its phenolic cargo HPQ (**Fig 4a**). The precipitating and thereby cellularly retained solid-state fluorophore HPQ was chosen for signal accumulation to overcome low TrxR levels. However, high probe dosage and long incubation times were needed to surpass the precipitation threshold (K_S_), and the crystalline HPQ precipitates that generate signal also stimulate inflammatory responses and are toxic to cells. A soluble cell-retained fluorophore product could solve both drawbacks, giving a more biocompatible probe (lower dosage that is less cellularly damaging but still quantifiable signal) that is also faster-quantifiable (since K_S_ need no longer be overcome). Thus, we patched RX1’s selenenylsulfide onto our rhodol scaffold, hoping the resulting probe **TR-TraG** would keep TrxR selectivity while accessing the advantages of cell-retained soluble fluorophores.

**Figure 4.**
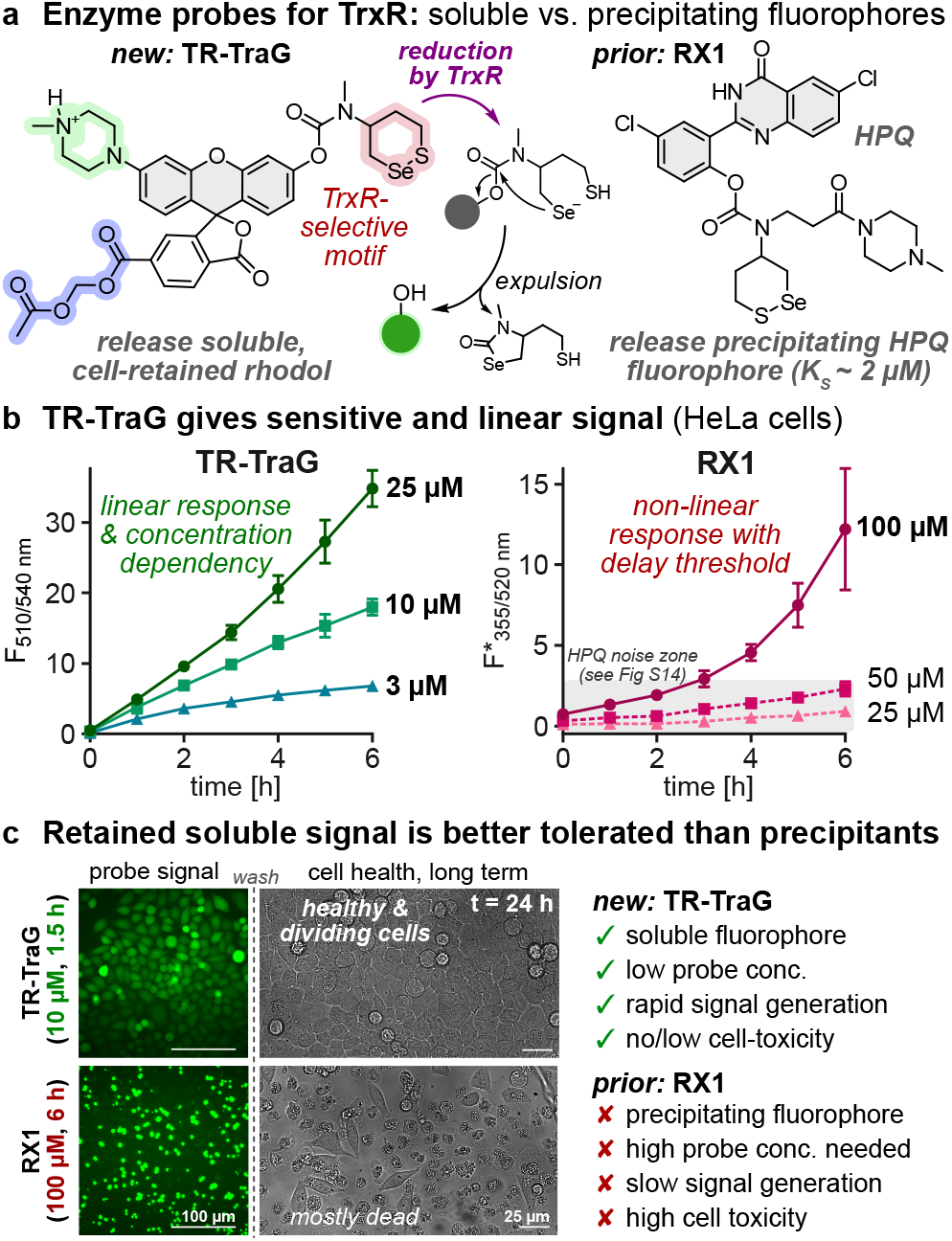
The TraG probe design can be modularly equipped with enzyme activation motifs to address different targets. **(a)** The TrxR1 probe **TR-TraG** (releases soluble **H-TraG**) compared to known probe **RX1** (precipitating fluorophore). **(b)** Cellular concentration-dependent fluorescence generation of **RX1** and **TR-TraG** in HeLa cells (*n*=3; error bars: SD). **(c)** Microscopy of HeLa cells treated with **TR-TraG** (10 µM) or **RX1** (100 µM) for 6 h, then washed, and kept for 24 h to assess cell viability (full data in **Fig S18**).

In cell-free experiments, **TR-TraG** was activated by even 20 nM of TrxR1 (vicinal selenolthiol); cell-free selectivity was decent over vicinal dithiols (thioredoxin 1 resisted up to 300 nM, lower resistance to glutaredoxin 1) or outstanding over thiol GSH (1000 mol equivalents of GSH reach only ∼15% activation after 4 h, i.e. the level reached by 0.002 mol eq of TrxR after 0.5 h; **Fig S15**). Pleasingly, in cellular assays **TR-TraG** signal mainly depended on TrxR activity: inhibition with electrophile TRi-1^[51]^ (HeLa and A549 cells, **Fig S16**) or genetic knockout (MEF cells, **Fig S17**) largely controlled its signal. Thus, the selenenylsulfide substrate does set the probe’s target-selectivity. We next examined some systematic benefits of the soluble cell-retained design.

A major technical drawback of precipitating fluorophores is their non-linear fluorescence response. In each cell, the released fluorophore concentration has to surpass K_S_ (HPQ: ∼2 µM) before true signal starts to be observable, whereas soluble fluorophores are theoretically detectable with linear activation response from the first molecule released. High-throughput plate reader assays with precipitating fluorophores also suffer from inter-cell variability since turnover must reach ∼2K_S_ *in the majority of cells* before overall signal becomes linear: again, an issue that does not affect soluble probes. Finally, the quenching in precipitation-based systems is often incomplete: even quenching one fluorescence channel (e.g. HPQ: ESIPT quenching by *O-*masking) does not suppress all channels (weak long-wavelength tail of normal emission by *O-*acylated probe, **Fig S18b**, 45 min); whereas xanthene spirocyclisation quenching can be complete. All these advantages were evident when comparing **TR-TraG** and RX1 in cellular assays. **TR-TraG** builds up signal linearly from time zero, proportional to its dosage, reaching usefully quantifiable signal even at ≤1 h at 3 µM (**Fig 4b**) while RX1 signal starts only at >3 h at 100 µM, with no true signal at lower times or doses. Such high RX1 exposure is incompatible with assays in tissues or organisms with limited, variable biodistribution: a limitation that **TR-TraG** escapes. The rhodol’s reproducible signal (**Fig S16**) also contrasts to the high variability in RX1 assays^[49]^ that results from their sensitivity to precipitation effects.

A major biological problem with precipitating fluorophores is that they cause cellular stress and cytotoxicity that limit or prevent long-term experiments and *in vivo* assays. Typical ways to run high-powered assays, e.g. first imaging and sorting by FACS to stratify cell populations, then further cultivation or parameter testing, are thus impossible. The rhodol **TR-TraG** instead allowed high-quality cell-resolved imaging at order(s) of magnitude lower probe exposure (10 µM, 90 min) than RX1 (100 µM, 6 h; see **Fig S18**), which should already result in far lower biological stress from the *probe*. Yet, we attribute the major difference to the *product* exposure. Cells treated with **TR-TraG** were healthy and continued dividing to confluency over 24 h (as did untreated controls), whereas RX1-treated cells stopped dividing and were <15% viable even after probe removal and culture for 24 h in fresh media (**Fig 4c**). Thus, **TraG** type probes will likely enable long-term cell tracking by bioactivity, e.g using FACS to resolve and study cell subpopulations: which at least in the context of TrxR probes is a novel and urgently needed advance.

## Conclusion

We designed a novel, modularly applicable fluorogenic probe scaffold for flexible use in sensitive biochemical activity imaging at low probe doses, that results in the linear generation of a biocompatible, cell-retained, bright fluorescence signal from the **H-TraG** fluorophore (λ_ex_ 504 nm, λ_em_ 531 nm). The combination of a basic amine and an intracellularly unmasked carboxylate on the spirocyclised rhodol precursor allows rapid cell loading and retention of the probe, as well as excellent post-wash retention of the fluorescent open-form rhodol product generated by *O-*unmasking, across different cell lines (hours in HEK and HeLa cells, up to 1 h in MEF, A549, and Hoxb8-derived macrophages). That the piperazine-rhodol seems to escape significant signal loss by passive diffusion or by active transport, contributes to the reliability of signal detection and confidence in signal quantification, even after long “post-wash” incubation times as would be encountered in multi-step cell biology assays (such as cell population sorting) or in situations with wash-in/wash-out (such as ADME kinetics during *in vivo* enzyme activity imaging). The signal is uniformly distributed across the whole cell with no compartmental accumulation, which is a further advantage for *in vivo* imaging in 3D environments.

Previous approaches to ensure cellular signal retention and thus biochemical activity integration have greatly relied on releasing precipitating fluorophore or intracellular alkylator products, which have biological as well as technical disadvantages. We applied our modular scaffold to generate two activity sensor probes showing the superior performance that a soluble fluorophore probe can achieve with a well-tempered cell-entry/exit profile.

(1) The hydrogen peroxide sensor **HP-TraG** senses exogenous and endogenous hydrogen peroxide in cells, adding a novel and milder cell-retained H_2_O_2_ probe to the probe toolbox. (2) **TR-TraG** images the cellular enzyme activity of thioredoxin reductase 1 (TrxR1), and outperforms the current probe RX1 in multiple respects: it more rapidly reaches higher sensitivity quantification, even at vastly lower probe loading, and delivers beneficial linear signal development, as well as allowing long-term cellular viability. We had assumed that the strong probe uptake across multiple cell lines indicates that cellular uptake relies mainly on passive diffusion, but it is also possible that these amine probes may profit from transporter-mediated uptake e.g. by organic cation transporters^[52,53]^. Nevertheless, the modular performance of this system, with two probes of rather different overall polarity that both perform strongly, promises the straightforward design and generation of a variety of other phenol-releasing probes centred on this scaffold (e.g. for *O-*unmasking by glycosidases or phosphatases, or by peptidases *via* self-immolative spacers), that can improve the sensitivity and biocompatibility of long-term-compatible cellular and *in vivo* molecular imaging.

## Supporting Information

Supporting notes and figures on the design and performance of lipidated fluorescein probes (**Figs S4−S8**); supplementary figures for the rhodol probes (**Figs S9−S18**); photocharacterisation (**Figs S19−S21, Table S1**); cell-free assays (**Figs S22−S27**) and all compound synthesis, analysis and biological methods. The authors have cited additional references within the Supporting Information.^[54-64]^

## Author Contributions

P.M. performed synthesis, chemical analysis, enzymatic cell-free studies, cell biology, and coordinated data assembly. L.D.-W., C.Z., N.A.V., D.B. and A.K. performed cell biology. L.D.-W., N.A.V. and D.B. performed confocal microscopy, image analysis and quantification. J.I.B. and T.K. performed synthesis. J.T.-S. and M.K. supervised cell biology and confocal microscopy. P.M. and O.T.-S. designed the concept and experiments. O.T.-S. supervised all other experiments. P.M. and O.T.-S. co-wrote the manuscript with input from all authors.

## Acknowledgements

This research was supported by funds from the German Research Foundation (DFG: Emmy Noether grant number 400324123 to O.T.-S.). The authors acknowledge support from the Joachim Herz Foundation (Research Fellowships to P.M. and J.T.-S.); the Studienstiftung des Deutschen Volkes (Ph.D. scholarships to P.M. and D.B.); and the Munich Graduate School of Systemic Neurosciences (D.B.). We are grateful to Dr. Johannes Morstein (Caltech) for collegial discussions around cellular uptake, retention, and localisation.

## Abbreviations

### Abbreviations

A549: human lung cancer cell line
ADME: absorption, distribution, metabolism, and excretion
AM: acetoxymethyl ester
DMEM: Dulbecco’s modified Eagle’s medium (cell culture media)
ESIPT: excited state intramolecular proton transfer
FACS: fluorescence-activated cell sorting (by flow cytometry)
FCS: foetal calf serum
Hoxb8: macrophage precursor cell line
HPQ: (2-(2’- hydroxyphenyl)-4(3*H*)-quinazolinone (fluorophore)
GSH: glutathione; HEK: human embryonic kidney cell line HEK293T
HeLa: human cervical cancer cell line
KS: solubility limit
MEF: mouse embryonic fibroblast cell line
PBS: phosphate buffered saline (buffer)
PMA: phorbol 12-myristate 13-acetate
RX1: molecular probe for TrxR activity with cell retained signal based on HPQ release and precipitation
SPiDER: molecular probe scaffold with cell retention of signal based on enzymatic unfurling of an alkylating ortho-quinone methide
TBAF: n-tetrabutyl-ammonium fluoride
TBAF: n-tetrabutyl-ammonium fluoride
TBAF: n-tetrabutyl-ammonium fluoride
TBAF: n-tetrabutyl-ammonium fluoride
TrxR: the mammalian selenoenzyme thioredoxin reductase

