## Supporting Information for "Cellularly-Retained Fluorogenic Probes for Sensitive Cell-Resolved Bioactivity Imaging"

**Author Contributions:** P.M. performed synthesis, chemical analysis, enzymatic cell-free studies, cell biology, and coordinated data assembly. L.D.-W., C.Z., N.A.V., D.B. and A.K. performed cell biology. L.D.-W., N.A.V. and D.B. performed confocal microscopy, image analysis and quantification. J.T.-S. and M.K. supervised cell biology and confocal microscopy. P.M. and O.T.-S. designed the concept and experiments. O.T.-S. supervised all other experiments. P.M. and O.T.-S. co-wrote the manuscript with input from all authors.

### 1 Overview of all fluorogenic probes and fluorophores

#### 1.1 Fig S1: Fluorescein-derived probes and fluorophores

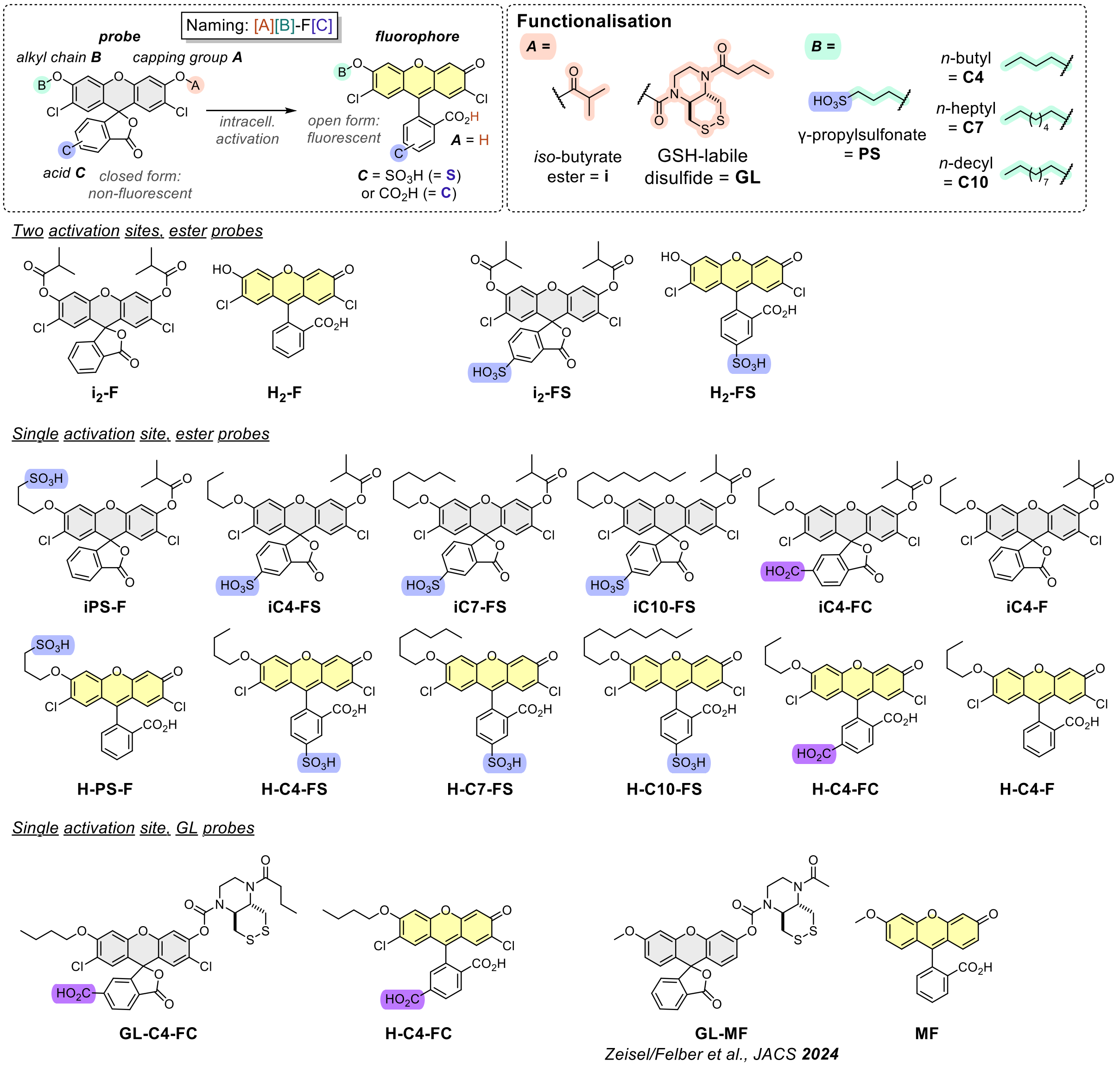

**Figure S1**: Compound naming rationale and structure overview of all fluorescein-derived probes and fluorophores.

#### 1.2 Fig S2: Rhodol-derived probes and fluorophores

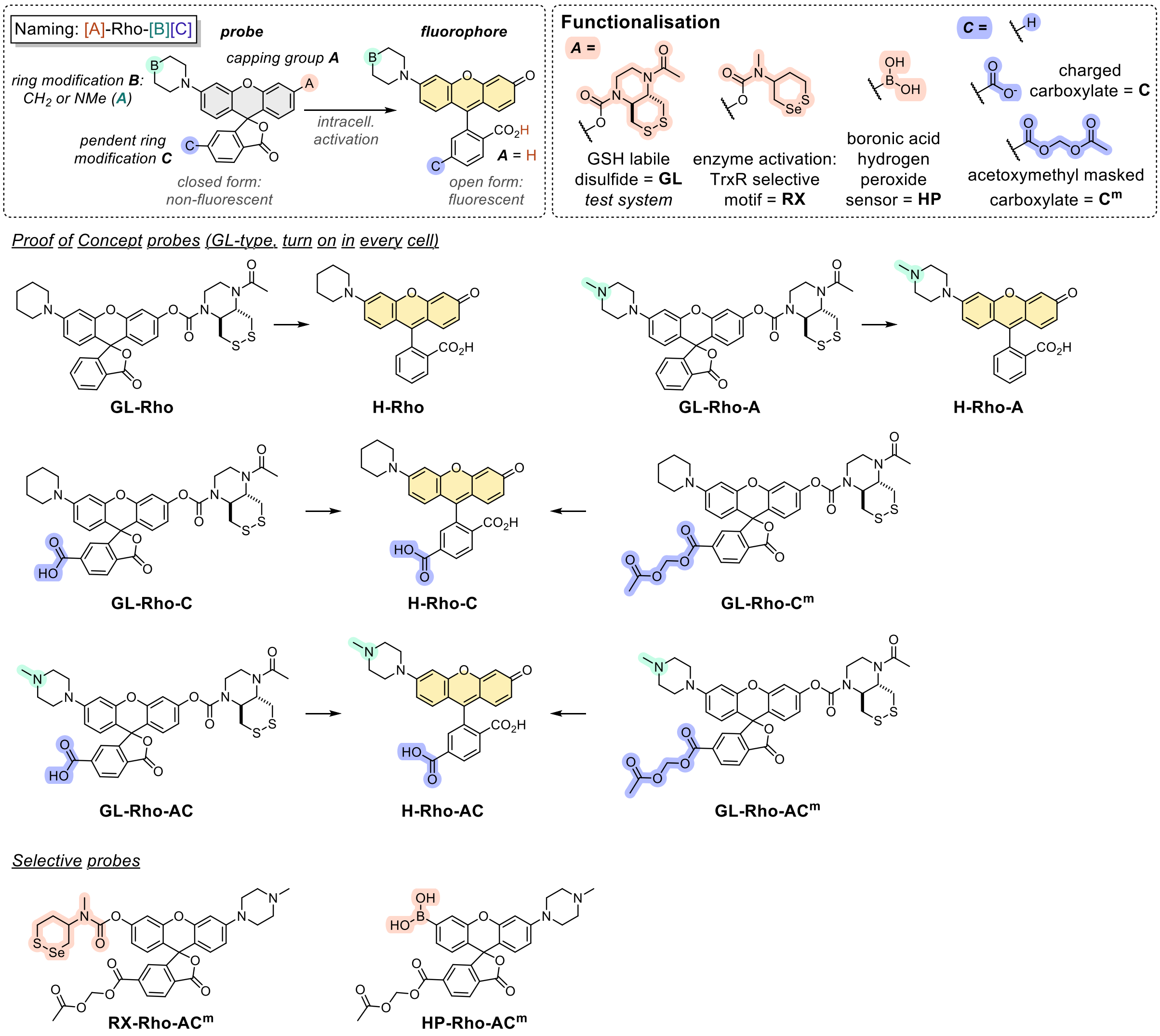

**Figure S2**: Compound naming rationale and structure overview of all rhodol probes and fluorophores.

#### 2 Fig S3: Cell retention approaches in the literature

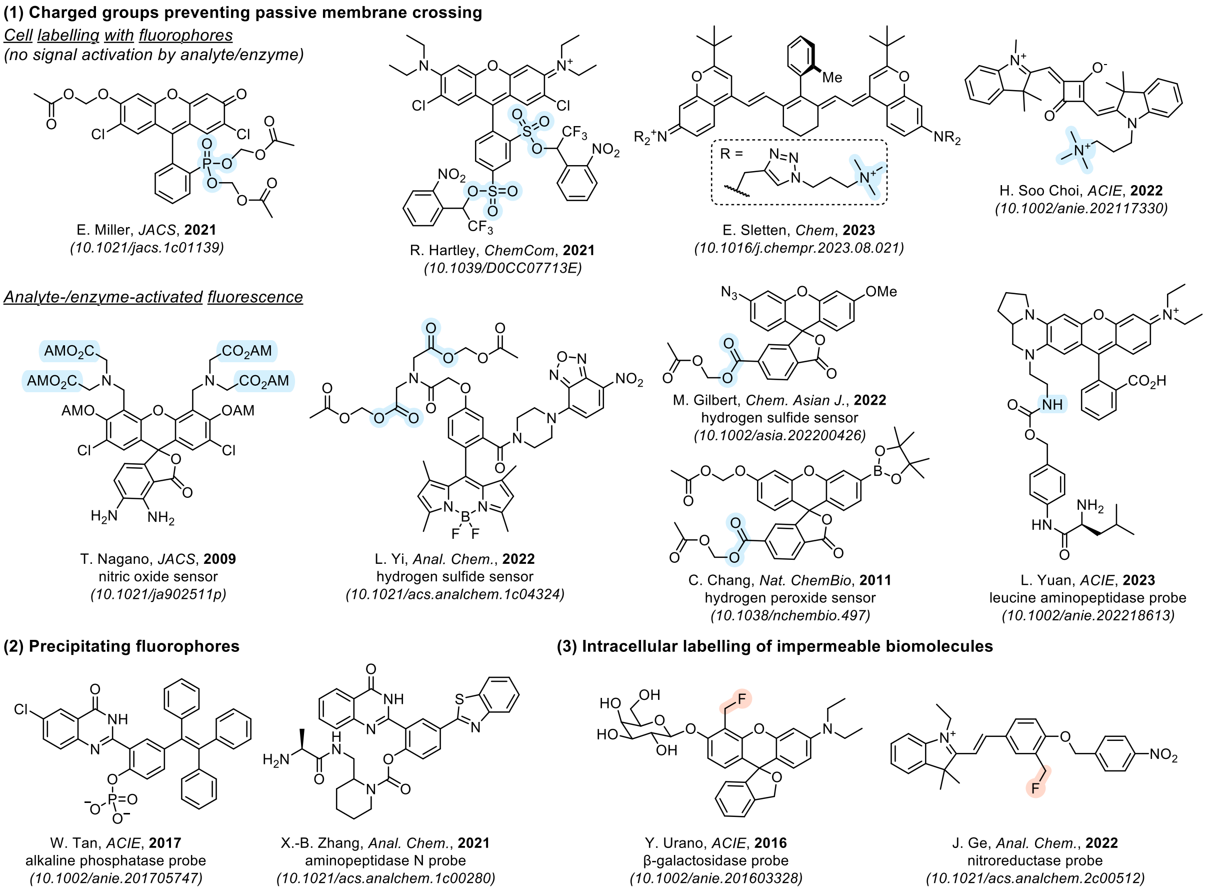

**Figure S3: Strategies for intracellular fluorophore and probe signal retention used in literature.**

The need for cell-retained probes has inspired many investigations of intracellular fluorophore trapping, with three general probe strategies emerging: (1) charge- and polarity-based impermeabilization by suppression of passive membrane crossing, (2) precipitation of the released fluorophore, and (3) intracellular labelling of impermeable biomolecules (e.g. proteins or glutathione) (overview of literature approaches in **Fig S3**).

(1) Charge-based cell retention is to date the most broadly utilised approach despite its limitations. Generally, these strategies use either cationic motifs such as tetraalkylammonium ions[1,2] or anionic groups such as carboxylates[3], phosphonates[4] or sulfonates[5] which cannot cross lipid bilayer by passive diffusion. The key challenge is cellular delivery of such polar groups which can be achieved in different ways: (i) cleavable masking groups that transform the charged functionality into lipophilic, membrane-permeable groups (e.g. acetoxymethyl carboxyl/phosphonate esters[3,4] or trifluoromethylbenzyl sulfonate esters[5,6]), (ii) endocytosis which can be induced by cell-penetrating peptides[7], and (iii) transporter-mediated uptake[1]. Most approaches focus on fluorophore delivery for cellular labelling and are not applicable for activity probes which generate fluorescence upon enzyme- or analyte-triggered activation. However, there are some examples investigating signal retention of activity probes which come with different limitations. Nagano and Yi developed sensors for nitric oxide[3] and hydrogen sulfide[8] which intracellularly release acetoxymethyl masked carboxylates and suppresses fluorescence by photoinduced electron transfer (PET) and FRET quenching before reacting with the analyte – an approach that limits applications to special reaction types and cannot be utilised for simple bond-cleavage reactions which would allow modular use for many types of analytes and enzymes. Other examples from Gilbert and Chang overcome this problem by generating phenol- and aniline-modified xanthenes for sensing hydrogen sulfide[9] and hydrogen peroxide[10] which would be translatable to other activating triggers but suffers from low fluorescence brightness or non-specific, partial intracellular signal generation which strongly reduces the sensitivity (opposing the goal main goal of retained probes: increased sensitivity and zero background). Yuan uses cationic retention releasing a basic amine for detecting hydrogen peroxide as well as leucine aminopeptidase and nitroreductase activity[11] and achieves signal turn on by PET quenching before activation requiring benzylic spacers whose 1,6-elimination influences signal turn-on kinetics and intracellularly releases electrophilic (aza)quinone-methides which can be cytotoxic, and the net positively charged fluorophore accumulates in lysosomes after activation. Overall, previously explored probe motifs come with different limitations in activation trigger modularity, fluorophore brightness, cellular uptake, release of reactive side products or undesired compartmentalisation (e.g. of basic amines to the lysosome).

(2) Precipitating fluorophores are a different approach to accomplish intracellular trapping of fluorescence. The water-insoluble fluorophore HPQ ((2-(2'-hydroxyphenyl)-4(3*H*)-quinazolinone) features excited-state intramolecular proton transfer (ESIPT)-based solid-state fluorescence with a large Stokes-Shift and bright signal.[12] HPQ-derived probes have been developed for alkaline phosphatase[13] and aminopeptidases[14] enabling not only cellular but even subcellular resolution of probe activation. Unfortunately, precipitating fluorophores come with several disadvantages: (i) their precipitation concentration threshold limits the sensitivity and renders activation below this threshold invisible, (ii) the water-insolubility of the fluorophore due to its lipophilicity and π--stacking limits the activating trigger to polar, solubilising motifs (such as phosphates or amino acids) to avoid pre-activation precipitation or membrane localisation, and – most problematically – (iii) the high cytotoxicity of intracellularly formed crystals which changes cell metabolism and makes visualisation of natural, biological activity impossible.

(3) Intracellular labelling of impermeable biomolecules as a cell retention approach was pioneered by Urano with the development of the so-called SPiDER probes. While previous approaches used active electrophiles to trap drugs[15] or fluorescent sensors[16], SPiDER probes are not electrophilic before activation and generate reactive quinone-methides upon activation which rapidly react with proteins and glutathione and have been successfully used for the development of glucosidase, peptidase, nitroreductase and hydrogen peroxide probes with durable cell retention.[17–21] However, the release of electrophiles can cause problems and influence the enzyme activity it is probing by reacting with the protein of interest[21] and the accumulation of reactive species can furthermore be toxic especially for high turnover cells.[19]

### 3 Figs S4-S6: Lipidated sulfo- and carboxyfluorescein probes

***This chapter expands on the main text's shorter description of the design and performance of the failed set of lipidated fluorescein probes.***

##### Sulfonated mono-capped fluorescein probes are not cell-retained

In a previous study we noticed that some mono-sulfonated fluoresceins can enter cells, which we used as a starting point for investigating and optimising cellular delivery and retention of such probes.[22] As an easy-to-synthesise test system for intracellular activation of phenolic probes, we used 2′,7′-dichlorofluorescein isobutyrate esters (**Fig S4a**) which are more stable against spontaneous ester hydrolysis in aqueous medium by sterical (isobutyrate vs. acetate) and electronic effects (n→π* interaction of chloride and ester) compared to ordinary fluorescein diacetate.[23] First, we investigated the previously described[22] diester **i$_2$-F** and its sulfonated analogue **i$_2$-FS** both featuring two esterase activation sites (resulting in non-linear fluorescence turn-on upon esterase activation as the first ester cleavage results in only 10% fluorescence activation with the other 90% coming from the second ester cleavage) and compared it to the sulfonated mono-ester **iPS-F** (one activation site, linear fluorescence turn-on). **i$_2$-F** releases dichlorofluorescein (**H$_2$-F**) with strong cellular signal which is actively exported from cells by anion transporters[24,25] and therefore shows reduced post-wash cell retention, its sulfonated analogue **i$_2$-FS** shows lower cellular signal but is able to escape the export pathway and is well retained after activation (**Fig S4b–d**). Unfortunately, these encouraging results did not translate to the single activation-site **iPS-F** which gives low cellular signal and is not cell retained. Thus, a different strategy was needed.

**Sulfonated diisobutyrate probe enters cells and is retained, single activation site sulfonate probe is not retained**

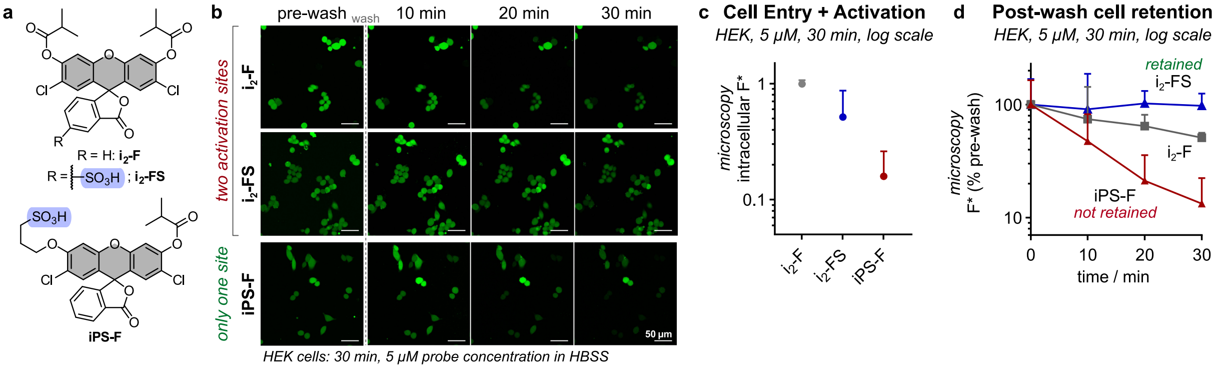

**Figure S4: Sulfonated mono-capped fluorescein probes are not retained. (a)** Structures of previously reported sulfonated fluorescein probes which served as entry for retention study; **(b)** Confocal microscopy: intracellular signal and post-wash retention of fluorescein diisobutyrate **i$_2$-F** compared to the sulfonated derivatives **i$_2$-FS** and **iPS-FS** (treatment with 5 µM probe in HBSS for 30 min, then wash (2× with HBSS); *scale bars: 50 µm*; transmission and CellTracker images: Fig S7); **(c)** Intracellular fluorescence signal quantified from microscopy images (treatment with 5 µM probe in HBSS for 30 min; *error bars: SD*; *n*=3); **(d)** Post-wash intracellular signal retention quantified from microscopy images (treatment with 5 µM probe in HBSS for 30 min, then wash (2× with HBSS), values normalised to pre-wash intensity; *error bars: SD*; *n*=3).

##### Lipidated sulfonate probes can be cell-retained

Nature uses medium-length lipids to enhance cell uptake and retention of natural products.[26] Inspired by this observation, we synthesised a set of *O*′-lipidated sulfofluorescein esters with different alkyl-chain lengths (*n*-butyl, *n*-heptyl and *n*-decyl: C4, C7 and C10) aiming for signal retention by combining charge introduction with lipidation (**Fig S5a**). The *O*′-alkyl fluoresceins show expected fluorescence properties (full discussion at **Fig S19**, **Table S1**), so next we investigated their cellular performance. Depending on the lipid tail length, different cellular uptake efficiency and post-wash retention is observed: **iC4-FS** and **iC7-FS** give low but decent cellular signal which is strongly reduced for **iC10-FS** with its much longer lipid tail, but all probes retain their fluorescence after washing (**Fig S5b–d**, the **iC10-FS** fluorescence was not quantified due to membrane-anchoring). The cellular signal distribution strongly depends on the lipid length: while **iC4-FS** (short tail) gives uniform signal distribution across the whole cell, **iC10-FS** (long tail) is anchored in the plasma membrane and slowly leaks into the cell after washing which renders it useless for activity imaging. The fluorescence of **iC7-FS** (medium-length tail) is mostly found in the cytosol and excluded from the nucleus. The signal intensity trend (**Fig S5c**) and the *in vitro* esterase activation kinetics (**Fig S24**) show lower signal for longer lipids, indicating reduced solubility and/or aggregation for more lipophilic probes with increased alkyl chain length, rendering **iC4-FS** the favourite candidate for optimising the cellular uptake.

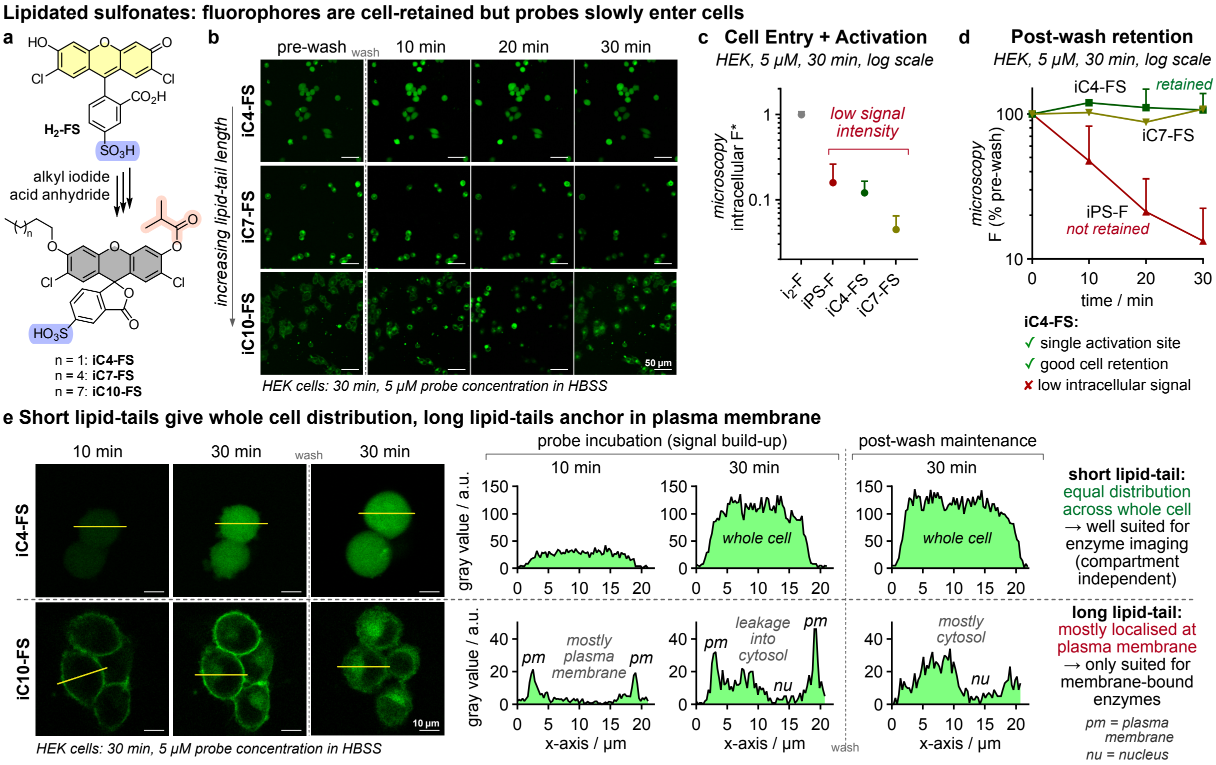

**Figure S5: Lipidated sulfonated fluorescein probes are cell-retained but slowly enter cells. (a)** Chemical structures and synthetic accessibility of lipidated sulfofluorescein isobutyrate probes **iC4-FS**, **iC7-FS** and **iC10-FS**; **(b)** Confocal microscopy: intracellular signal and post-wash retention of lipidated fluorescein probes (treatment with 5 µM probe in HBSS for 30 min, then wash (2× with HBSS); *scale bars: 50 µm*; transmission and CellTracker images: Fig S7); **(c)** Intracellular fluorescence signal quantified from microscopy images (treatment with 5 µM probe in HBSS for 30 min; *error bars: SD*; *n*=3); **(d)** Post-wash intracellular signal retention quantified from microscopy images (treatment with 5 µM probe in HBSS for 30 min, then wash (2× with HBSS), values normalised to pre-wash intensity; *error bars: SD*; *n*=3); **(e)** Confocal microscopy images of intracellular signal distribution for **iC4-FS** and **iC10-FS** and quantification of the gray values distribution across one cell (treatment with 5 µM probe in HBSS for 30 min, then wash (2× with HBSS); *scale bars: 10 µm*).

#### Lipidated carboxylate probes appeared to improve cell uptake but were ultimately found to rely on unphysiological membrane integrity destabilisation for this uptake, which stopped our development

To improve cellular uptake, we synthesised the short lipid tail carboxylate probe **iC4-FC** (**Fig S6a**) exchanging the permanently deprotonated sulfonate **iC4-FS** ($pK_a \approx -2$)[27] for a reversibly protonatable carboxylate ($pK_a \approx 4.3$). This less polar probe promisingly gave 20-fold higher cellular signal compared to its sulfonate-version while maintaining the post-wash retention of the fluorophore and the whole-cell signal distribution (**Fig S6b−d**). These results indicated that the signal retention is caused mostly by the lipid-tail while the acid-functionality influences the cellular uptake, so we investigated the cellular performance of a C4-tail probe without a charged group **iC4-F** (**Fig S6e**). Surprisingly, we found absolutely no cellular signal development which is in line with the cell-free esterase assay which also showed no turn-on of **iC4-F** presumably due to the high lipophilicity and hence insolubility and aggregation in aqueous medium as well as potentially membrane localisation in cells which overall makes the probe biologically unavailable.

In summary, the **C4-FC** motif seemed to deliver many of the desired features of a cell-retained probe with single-site activation, rapid cell entry, post-wash signal retention and uniform distribution in the whole cell. Problematically, with our proof-of-concept probe **iC4-FC** we observed severe changes in cell morphology such as cell rounding and blebbing (**Fig S6d**, **Fig S8**) which we first attributed to the use of HBSS buffer instead of DMEM supplemented with fetal calf serum (FCS) where the cell viability is improved. We used HBSS as default medium for our ester probes since the dichlorofluorescein isobutyrates are hydrolytically unstable in supplemented (nucleophile containing) cell culturing media such as DMEM supplemented with fetal calf serum (FCS) – and the hydrolytic instability even increases with solubilised FS- and FC-type probes compared to more lipophilic probes such as **$i_2$-F** (see cell-free hydrolytic stability in **Fig S22**).

To avoid spontaneous probe hydrolysis in FCS-supplemented DMEM, we synthesised the hydrolytically stable carbamate probe **GL-C4-FC** which is rapidly activated by (intracellularly abundant) glutathione (GSH, intracellular concentrations ~1−5 mM[28]; cell-free GSH activation in **Fig S25**) while being stable in typical cell culture media with low GSH concentrations (cell-free stability in cell culture media: **Fig S23**). Comparing the performance of **GL-C4-FC** in HBSS and in DMEM revealed an unexpected behaviour: cells treated in HBSS are strongly fluorescent but show impaired cell morphology (rounding and blebbing) at high probe concentration (10 μM), while negligible signal is generated when treated in DMEM where a healthy morphology is maintained. Cells cultured in either HBSS or DMEM without probe addition are also healthy, showing that HBSS itself does not impair the cells under the treatment conditons. Also, when the probe is treated in HBSS at lower concentrations (3 and 1 μM) the signal disproportionally decreases, while the cell morphology improves gradually (**Fig S8**) which clearly shows that our FS- and FC-type probes can only enter cells with disrupted membranes. Furthermore, these probes themselves damage the plasma membrane together with the simple buffer HBSS, while the nutrient-supplemented DMEM prevents the membrane disruption of the probes rendering them impermeable. We reason that the detergenic character of our negatively charged, lipidated fluoresceins together with the lack of nutrients in HBSS causes the membrane disruption which the probes rely on to enter the cells. Unfortunately, we did not see a strategy to avoid this problem with the carboxyfluorescein motif and therefore abandoned this approach and tested rhodol scaffolds instead. However, we gained some insights that guided us for the design of rhodol probes: (1) negative charges should be avoided, but if intracellularly released can assist cell retention if they are not actively translocated out of the cell by transport mechanisms; (2) solubility tags should be used to ensure bioavailability of the probe (since very lipophilic probes such as **iC4-F** are not even activated by isolated esterase); and (3) the proof-of-concept probes should allow testing in standard cell culturing media such as DMEM (with FCS), so hydrolytically stable carbamates are better suited than esters.

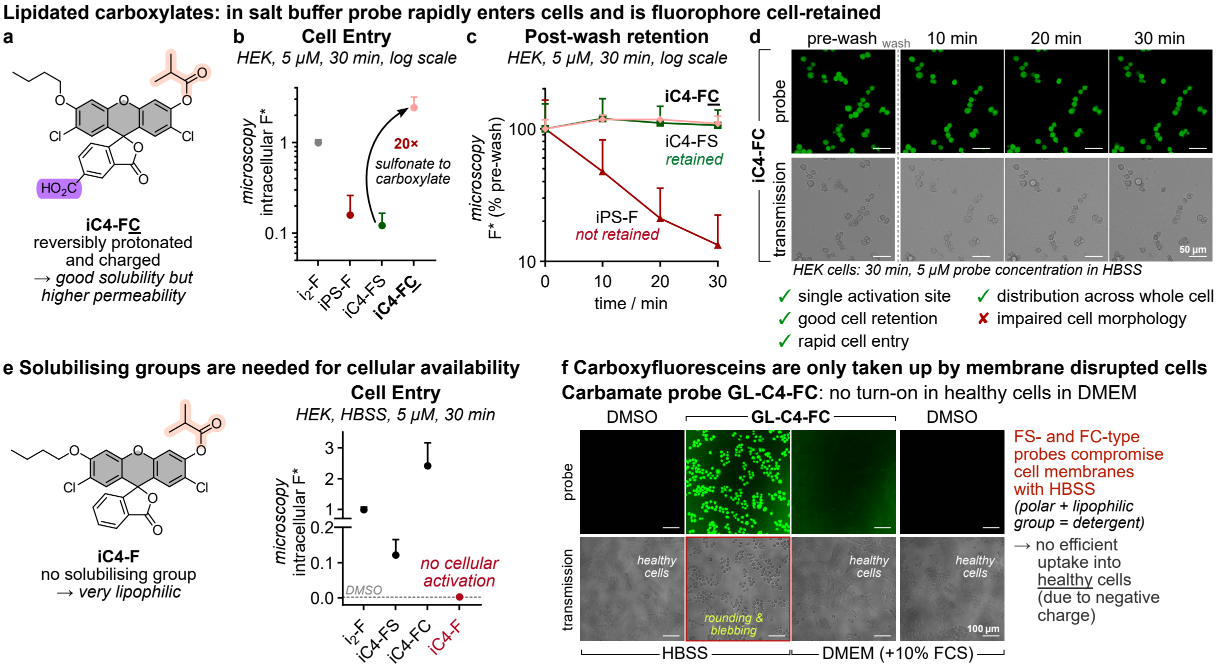

**Figure S6: Lipidated carboxy-fluorescein probes rapidly enter cells and are retained. (a)** Chemical structure of **iC4-FC** which is reversibly protonated and charged under physiological conditions; **(b)** Intracellular fluorescence signal quantified from microscopy images (treatment with 5 μM probe in HBSS for 30 min; *error bars: SD*; *n*=3); **(c)** Post-wash intracellular signal retention quantified from microscopy images (treatment with 5 μM probe in HBSS for 30 min, then wash (2× with HBSS), values normalised to pre-wash intensity; *error bars: SD*; *n*=3); **(d)** Confocal microscopy: intracellular signal and post-wash retention of **iC4-FC** (treatment with 5 μM probe in HBSS for 30 min, then wash (2× with HBSS); *scale bars: 50 μm*; CellTracker images: Fig S7); **(e)** Intracellular fluorescence signal of **iC4-F** (with no solubilising groups) quantified from microscopy images (treatment with 5 μM probe in HBSS for 30 min; *error bars: SD*; *n*=3); **(f)** The hydrolytically stable carbamate probe **GL-C4-FC** only generates signal when treated in HBSS but not in DMEM (with 10% FCS) in HEK293T cells (images after 30 min probe incubation (10 μM), no washing, *scale bars: 100 μm*).

### 4 Supplementary Figures

#### 4.1 Fig S7: Fluorescein probe microscopy: all channels

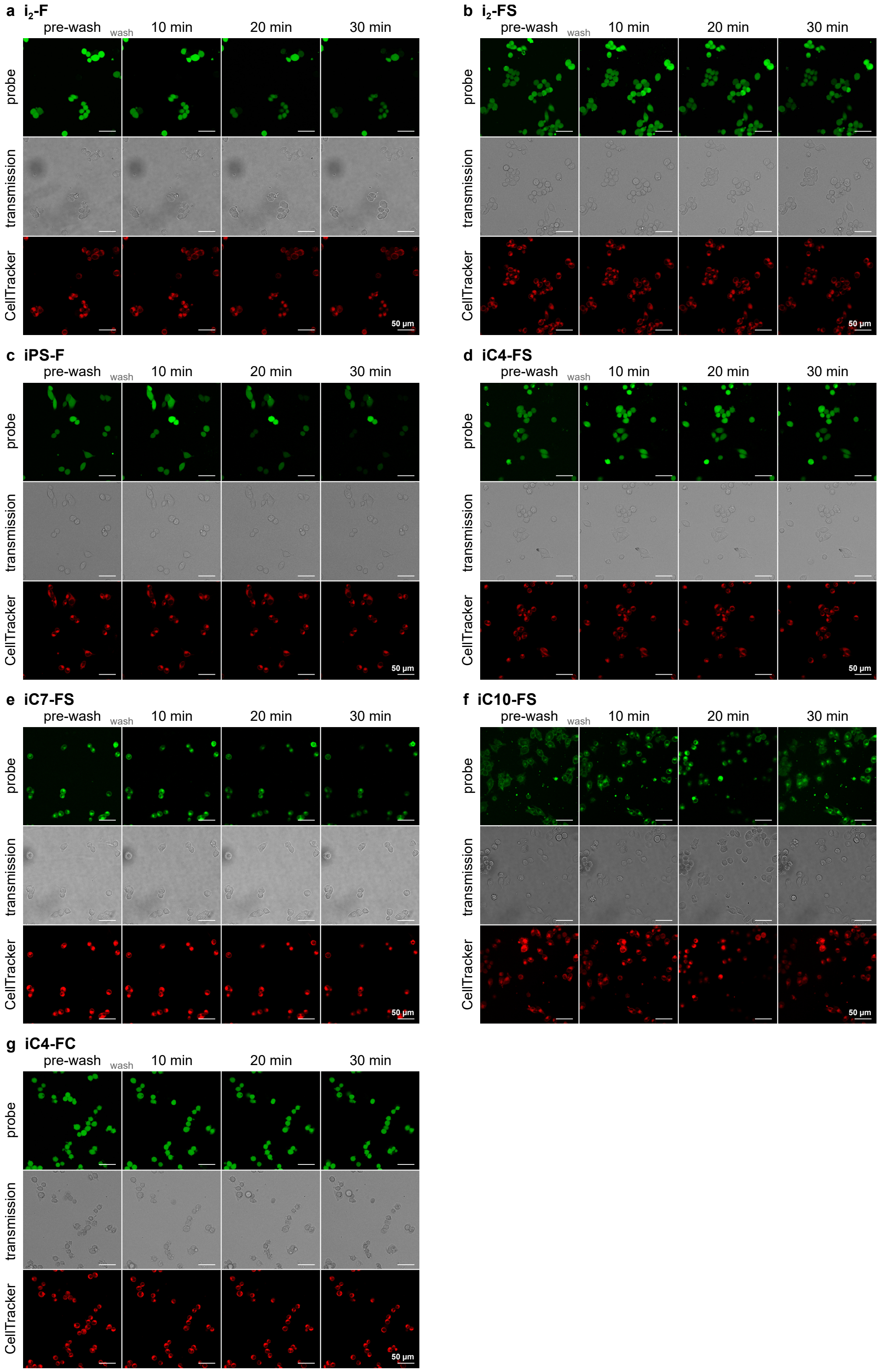

**Figure S7: Full figure for confocal microscopy of fluorescein probes** (all channels: probe, transmission and CellTracker Red). Cells were pre-treated with CMTPX (1μM for 30 min), then treated with the probes (5 μM probe for 30 min in HBSS) and imaged for the pre-wash image. Then the medium was removed (2× wash with HBSS) and post-wash images were acquired after 10, 20 and 30 min (*scale bars: 50 μm*).

#### 4.2 Fig S8: Comparison of GL-C4-FS and GL-C4-FC

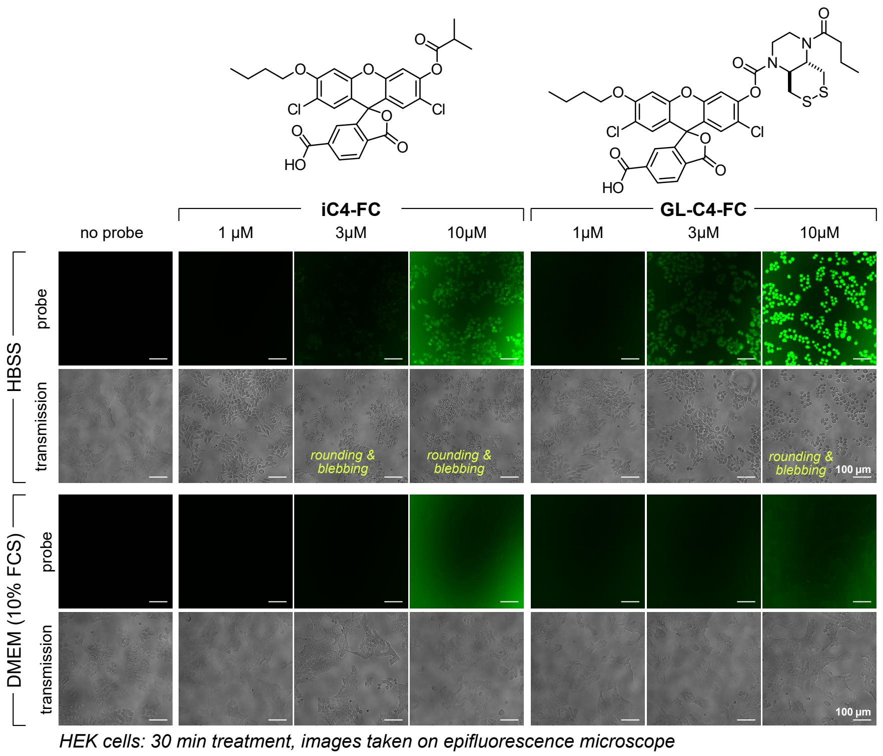

**Figure S8: Comparison of the carboxyfluorescein probes iC4-FC (ester) and GL-C4-FC (hydrolytically stable carbamate) in HBSS vs. DMEM (+10% FCS) at different concentrations (1, 3 and 10 µM).** Epifluorescence microscopy images of HEK293T cells treated 1–10 µM probe for 30 min in different media shows intracellular fluorescence but strongly impaired cell morphology after treatment in HBSS while cells remain healthy but non-fluorescent in DMEM (*scale bars: 100 µm*).

#### 4.3 Fig S9: Cell entry & retention of rhodol probes

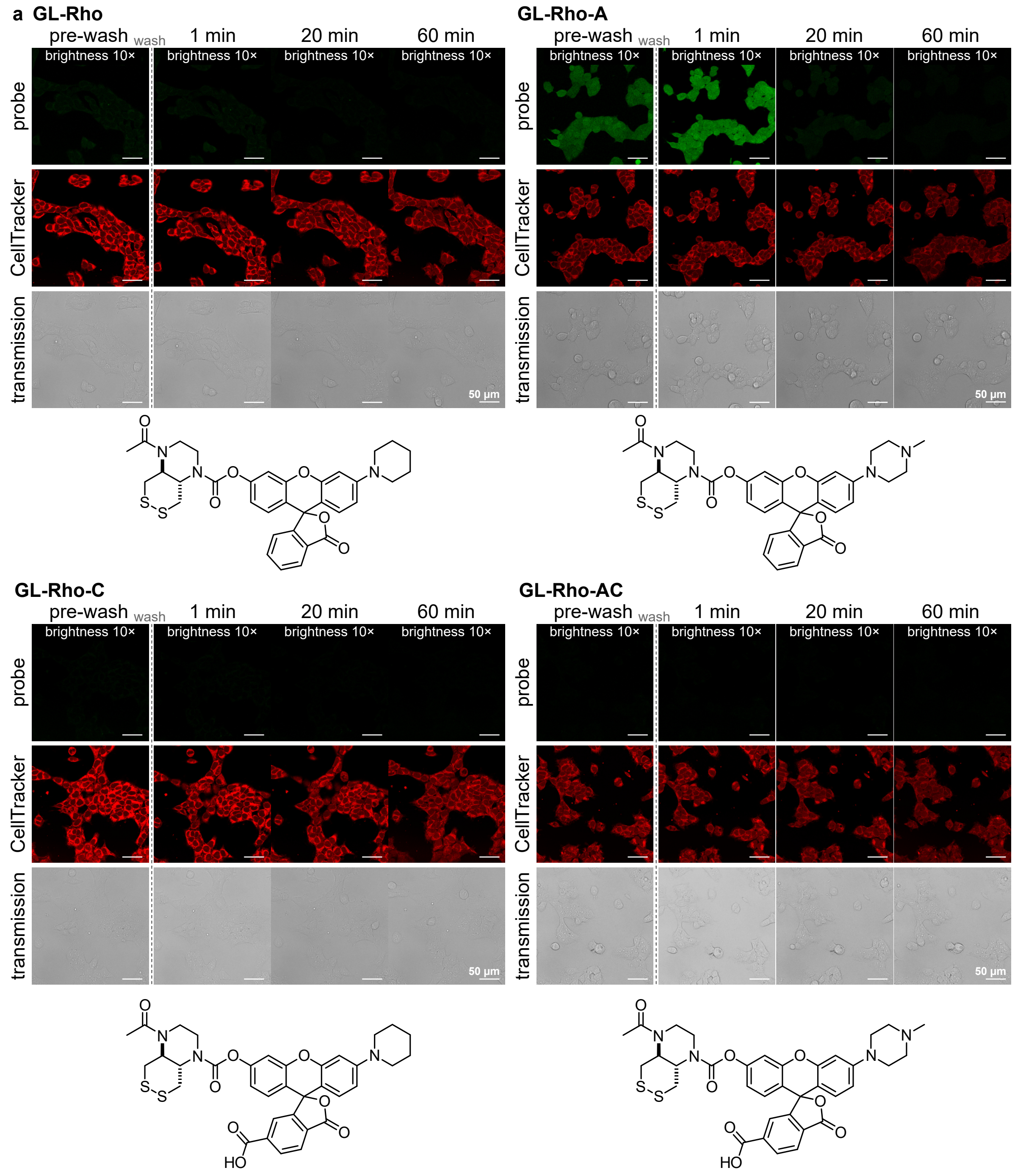

**Figure S9 Part 1: Comparison of cellular turn-on and retention for all rhodol probes by confocal microscopy** (full caption below at Part 2 of this figure).

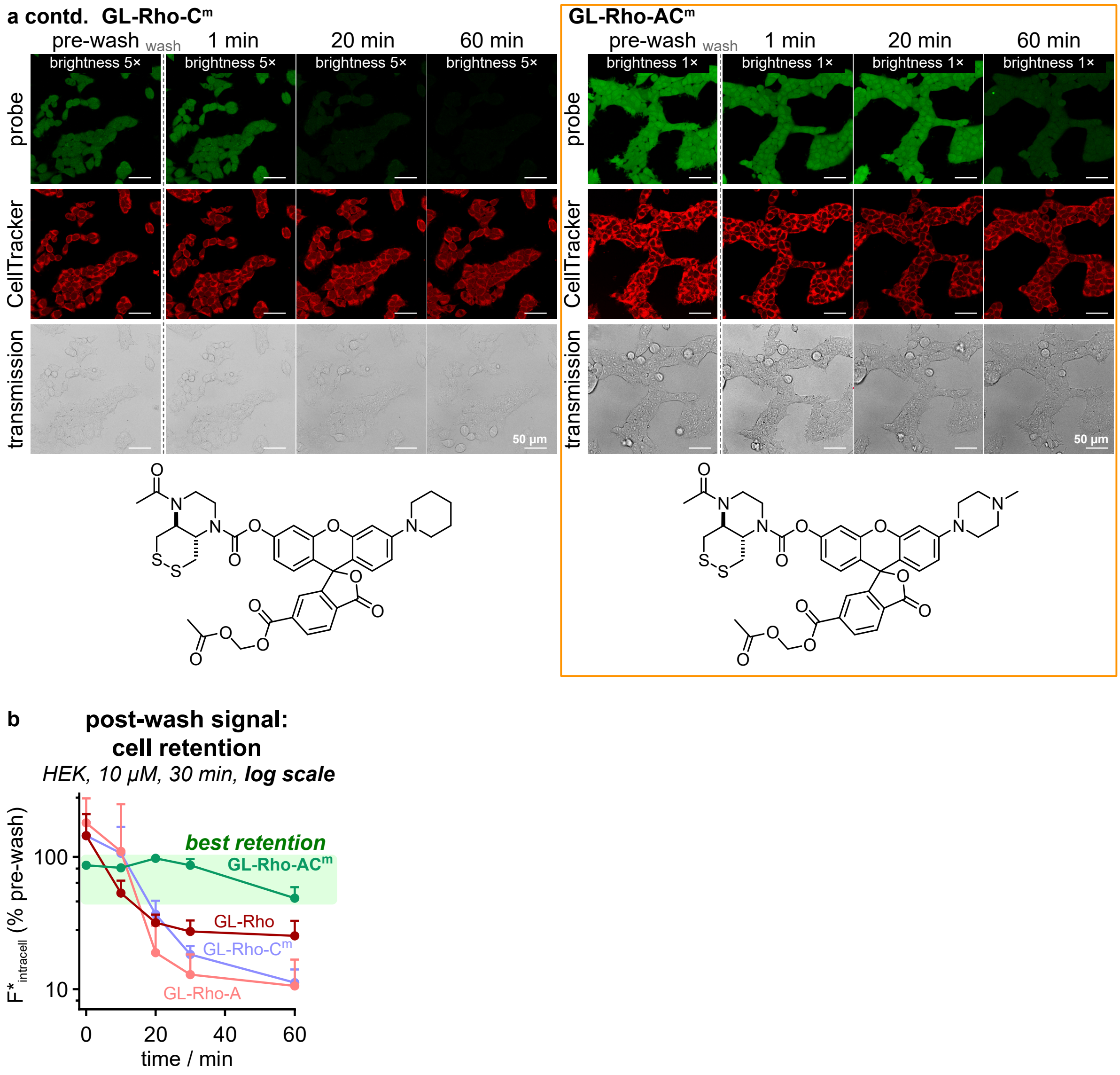

**Figure S9 Part 2: Comparison of cellular turn-on and retention for all rhodol probes by confocal microscopy:** **(a)** HEK293T cells; pre-wash images taken after 30 min treatment with 5 µM probe in DMEM (with 10% FCS); post-wash images after 2× wash with DMEM after 1, 20 and 60 min incubation (*scale bars: 50 µm*); **(b)** the intracellular fluorescence intensities were quantified to determine signal retention and are plotted in the graph below the images (n=3; error bars: SD).

Though **Fig 2c** (platereader assay, where signal is integrated over intra- and extra-cellular spaces) shows that **GL-Rho** enters cells and becomes activated, its exit (which **Fig 2d** timepoint 0 shows is ~80% complete before the cells have even been washed) is presumably too fast for any intracellular build-up to be seen in microscopy (**Fig 2d** later timepoints, and **Fig S9a**).

#### 4.4 Fig S10: Cellular distribution of GL-Rho-AC$^m$ signal

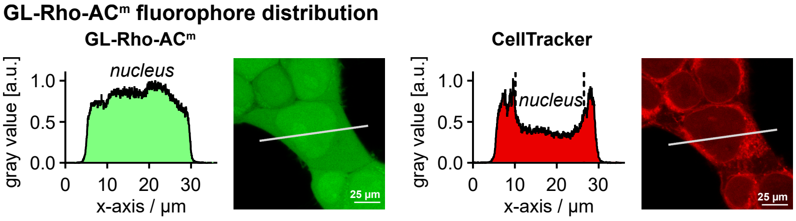

**Figure S10: Cellular fluorescence distribution of GL-Rho-AC$^m$** (after 30 min treatment, CellTracker signal for comparison (excluded from nucleus); *scale bars: 25 µm*).

#### 4.5 Fig S11: Entry and retention across several cell lines

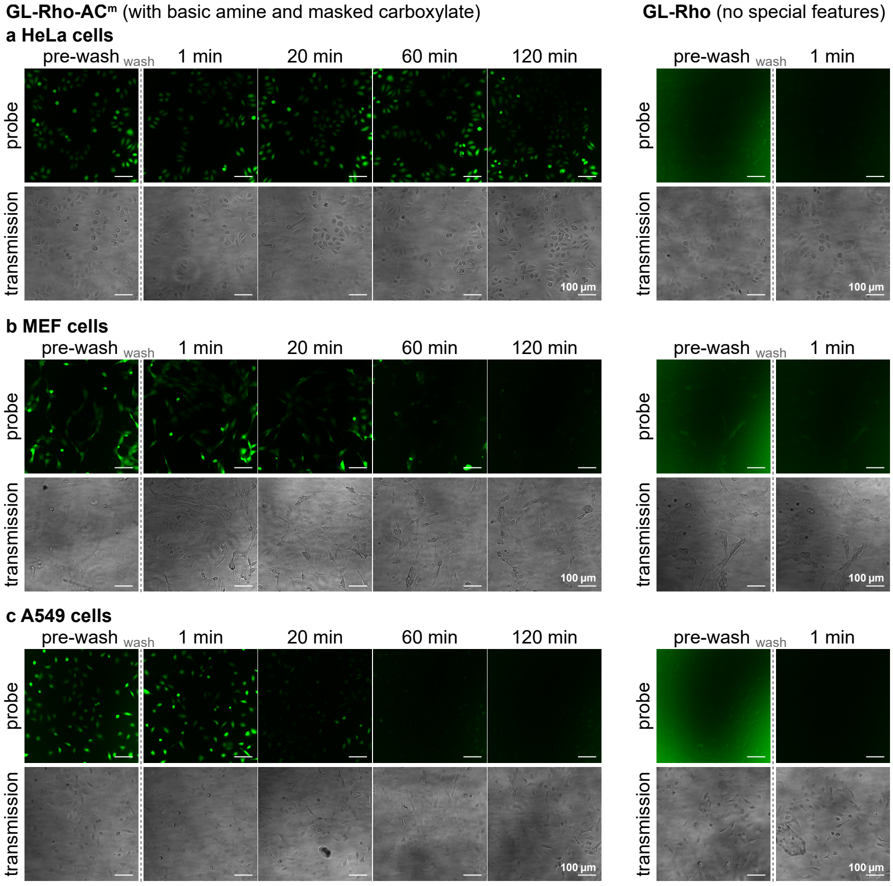

**Figure S11: Post-wash signal retention of GL-Rho-AC$^m$ compared to non-functionalised rhodol GL-Rho in different cell lines: (a)** HeLa, **(b)** MEF, or **(c)** A549 cells were treated with 5 µM probe in DMEM (+10% FCS) for 30 min (pre-wash image), then washed (2× with DMEM) and imaged after again for post-wash retention for up to 2 h (*scale bars: 100 µm*).

#### 4.6 Figs S12-S14: Characterisation of hydrogen peroxide sensor HP-TraG

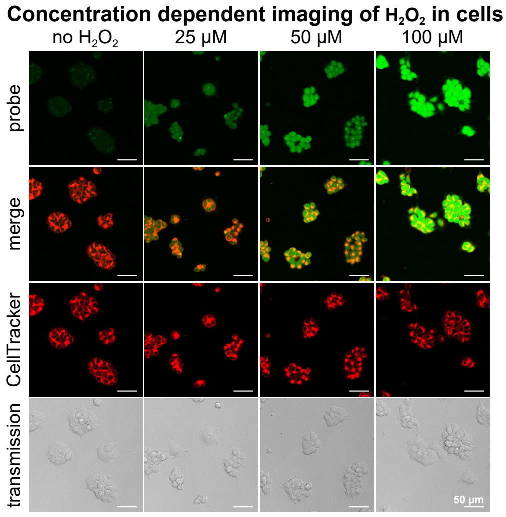

**Figure S12: Full figure panel of Figure 4c.** Concentration dependent activation of **HP-TraG** in cells (10 µM, 15 min loading before $H_2O_2$ addition, then 60 min $H_2O_2$ treatment; *scale bars: 50 µm*).

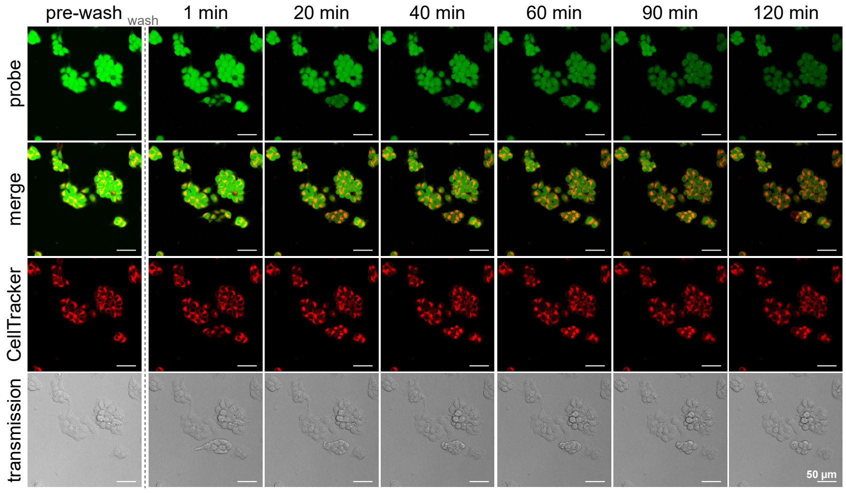

**Figure S13: Full figure panel of Figure 4e.** Confocal microscopy images of post-wash signal retention of **HP-TraG** (10 µM, 15 min loading before $H_2O_2$ addition) in HEK cells treated with $H_2O_2$ (100 µM for 1 h; *scale bars: 50 µm*).

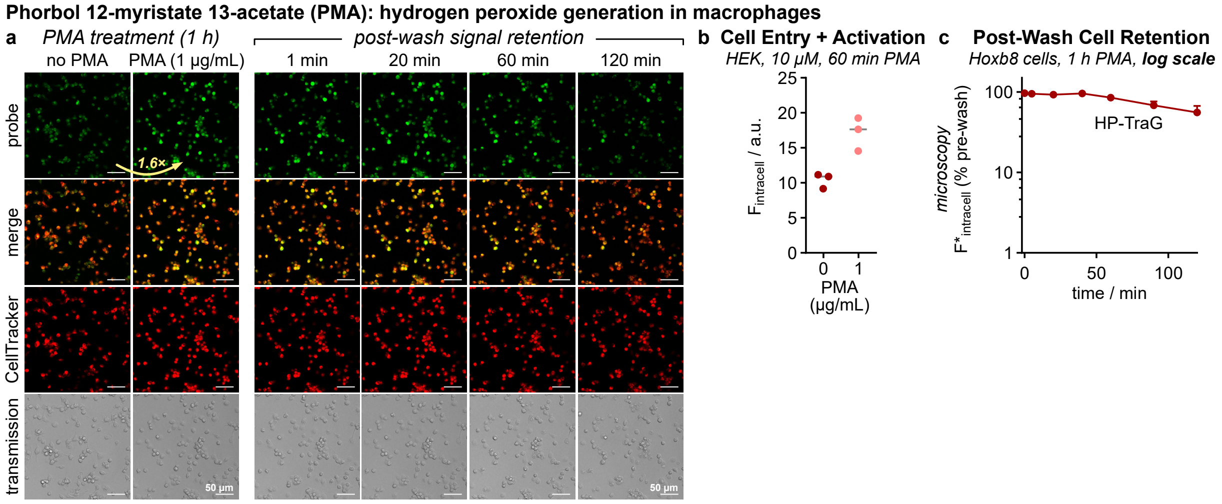

**Figure S14:** Hoxb8-derived macrophages activated with phorbol 12-myristate 13-acetate (PMA, 1 µg/mL, 1 h) that triggers $H_2O_2$ production which is visualised by **HP-TraG** (10 µM, 15 min loading before PMA addition): **(a)** microscopy images (*scale bars: 50 µm*); **(b)** intracellular **HP-TraG** signal after PMA treatment (1 h, 1 µg/mL, 10 µM probe loading for 15 min before PMA addition) and **(c)** the post-wash signal retention quantified from images in panel a ($n$=3; error bars: SD).

#### 4.7 Fig S15-S18: Characterisation of TrxR enzyme probe TR-TraG

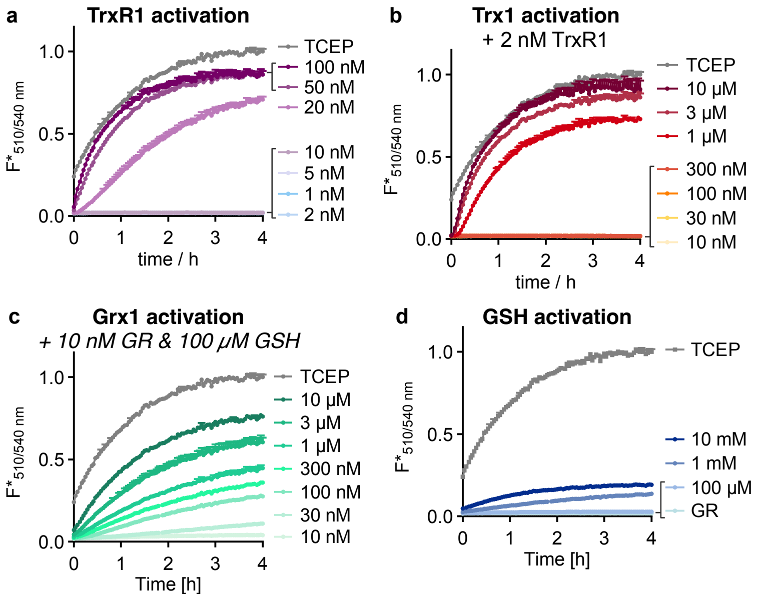

**Figure S15: Activation of TR-TraG by key redox enzymes and gluthathion (GSH): (a)** Target enzyme TrxR1, **(b)** Trx1 (+2 nM TrxR1 which is needed for Trx1 recovery by reduction), **(c)** Grx1 (+10 nM GR and 100 μM GSH which is needed for Grx1 recovery by reduction), and **(d)** GSH; Conditions: 10 μM **TR-TraG** in TE buffer at 37 °C for 4 h, TCEP: positive control (100 μM); NADPH co-factor concentration in all samples: 100 μM.

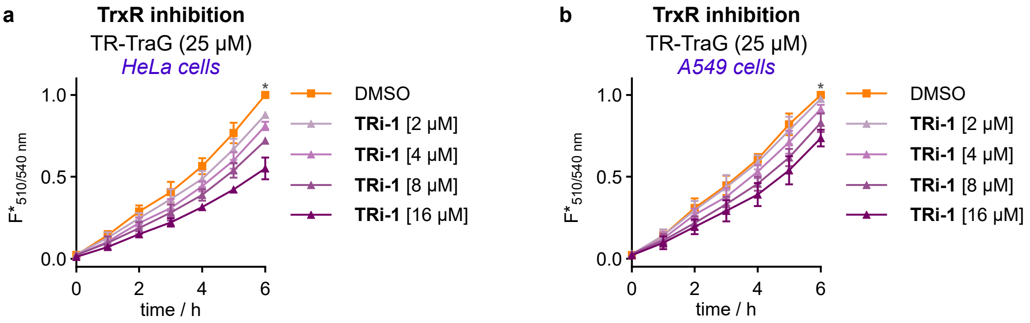

**Figure S16: TrxR inhibition dependent TR-TraG signal in HeLa and A549 cells.** Cells were treated with TrxR inhibitor TRi-1 at 2–16 μM for 3 h before medium exchange (2×) and probe addition (t = 0; 10 μM in DMEM (+10% FCS)), then the fluorescence was measured for 6 h (values normalised to DMSO-cells after 6 h, marked with asterisk; *error bars: SD*; *n*=3). The reproducibility of the TR-TraG fluorescence values throughout the cell culture experiments (small relative error, typ. ±<10%) contrasts to the stronger variability of the previously-published probe RX1 (typ. ±>20%)[29], which we assign to the reprodubility of its soluble (rather than precipitating) fluorogenic cargo.

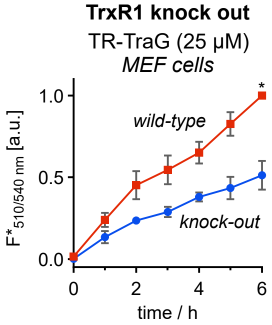

**Figure S17: TR-TraG** is activated in TrxR expressing MEF cells while the signal is strongly reduced when TrxR is knocked out (probe treatment at 25 μM and 37 °C in Se-supplemented DMEM (+ 10% FCS); *error bars: SD*; *individual experiments normalised to wild-type endpoint (marked with asterisk); n*=3).

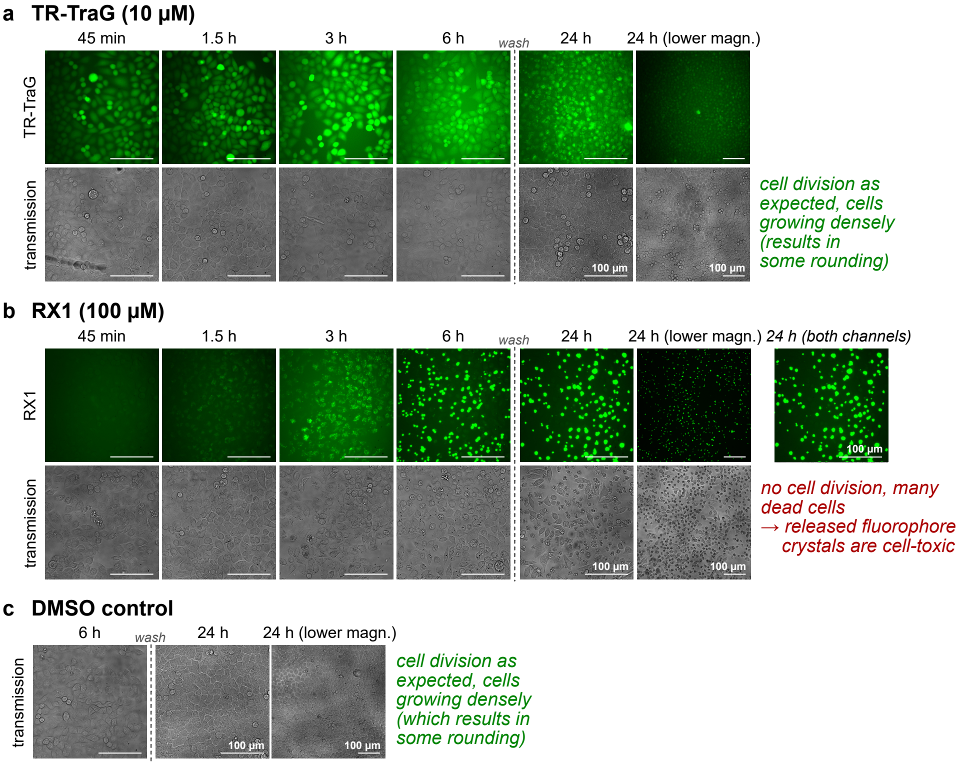

**Figure S18: Full figure: comparison of TR-TraG with RX1.** HeLa cells were treated with **TR-TraG** (10 µM, panel a) or **RX1** (100 µM, panel b) and imaged after 45 min and 1.5, 3 and 6 h, then washed (2×) and imaged again after 24 h to show post-treatment cell viability compared to untreated cells (DMSO control, panel c) (*scale bars: 100 µm*).

### 5 Cell-free photocharacterisation, probe stability, and probe activation

#### 5.1 Figs S19-S21: Photocharacterisation

##### 5.1.1 Fluoresceins: Absorption and emission spectra

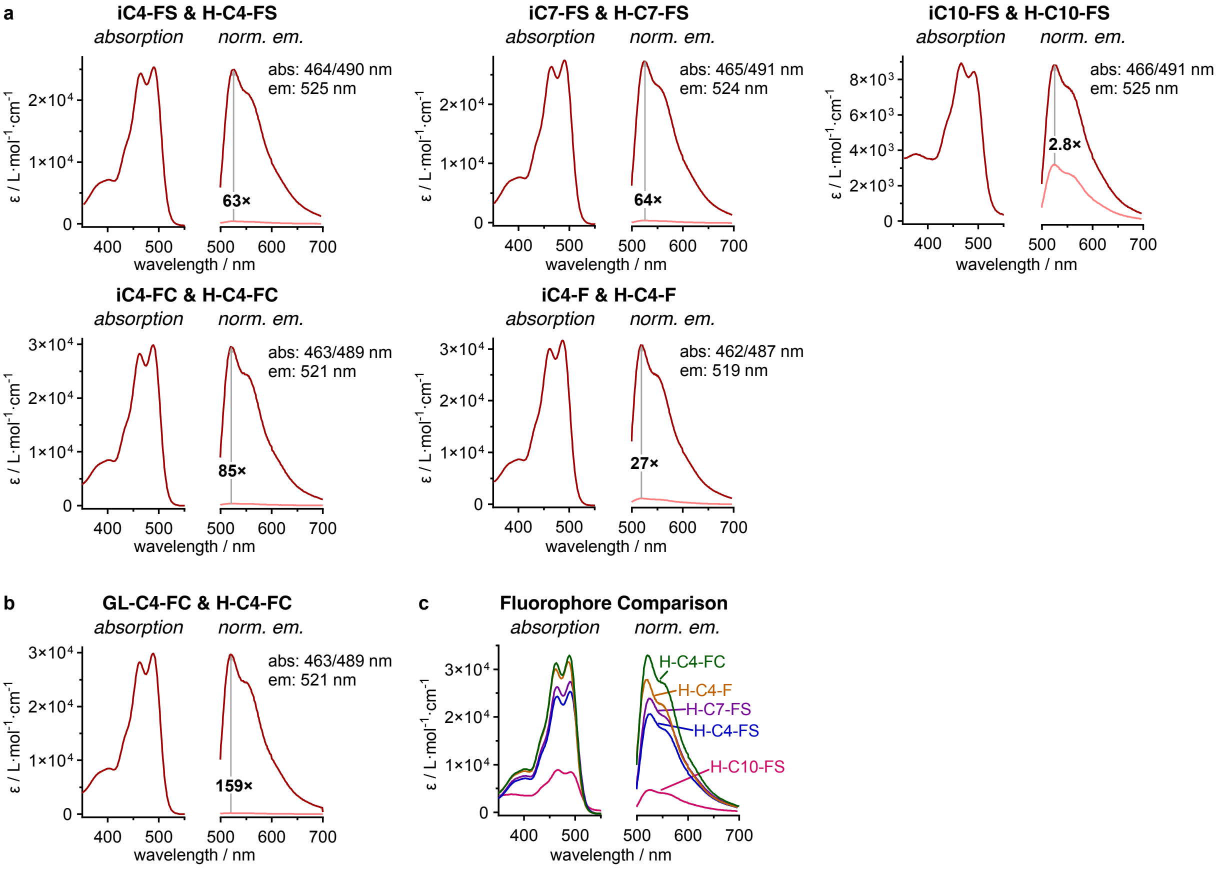

**Figure S19**: Absorption and emission spectra of fluorescein probes and fluorophores: **(a)** *iso*-butyrate ester probes and **(b)** GL-probe **GL-C4-FC** (10 µM in PBS, pH = 7.4; *dark red line: fluorophore; light red line: probe; grey, emission spectra:* $\lambda_{ex}$ *= 495 nm*); **(c)** Comparison of absorption and fluorescence emission of all fluorescein fluorophores.

The monoalkylated fluoresceins have two absorption maxima at ca. 465 and 490 nm, with slight red-shifting for sulfonate and carboxylate probes as expected for electron withdrawing substituents.[30] The probes have 3-fold weaker absorbance at 485 nm compared to unmodified, symmetric fluorescein, and their fluorescence quantum yields ($\lambda_{max}$ ~525 nm) are reduced from ~0.8 to ~0.2 (**Table S1**), which are expected results for the unsymmetric chromophore.[31] Very low absorption and emission is observed for the lipophilic **H-C10-FS**, presumably due to aggregation effects. All (spirocyclised) probes are non-fluorescent and feature very good turn-on ratios to the fluorophore (except **iC10-FS** with its very weak fluorescence of the fluorophore).

##### 5.1.2 Rhodols: Absorption, excitation and emission spectra

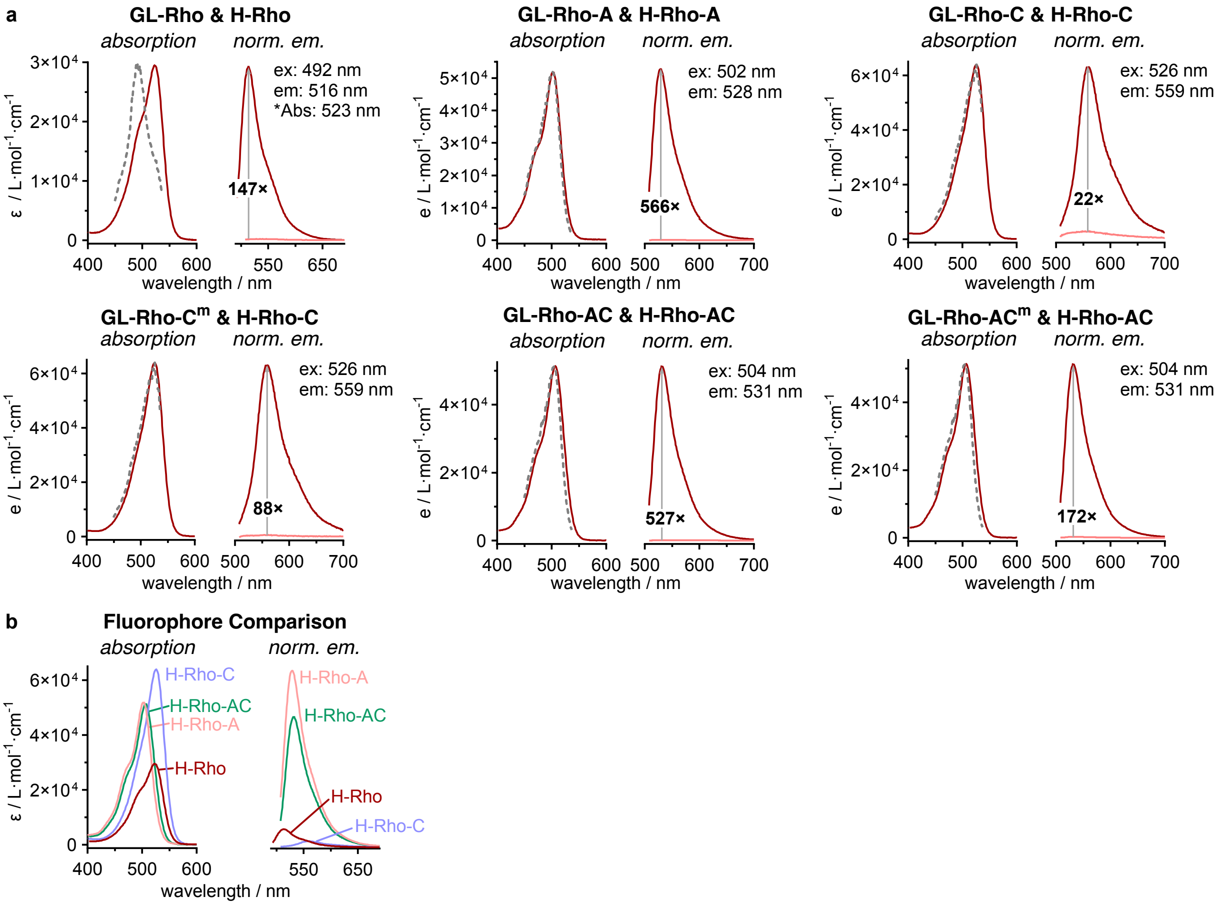

**Figure S20**: **(a)** Absorption, excitation and emission spectra of rhodols fluorophores and GL-probes (10 µM in PBS, pH = 7.4; *dark red line: <u>fluorophore</u>; light red line: <u>probe</u>; grey, dashed line represents excitation spectra: $\lambda_{em}$ = 550 nm; emission spectra: $\lambda_{ex}$ = 495 nm*); **(b)** Comparison of absorption and fluorescence emission of all rhodol fluorophores (*$\lambda_{ex}$ = 495 nm*).

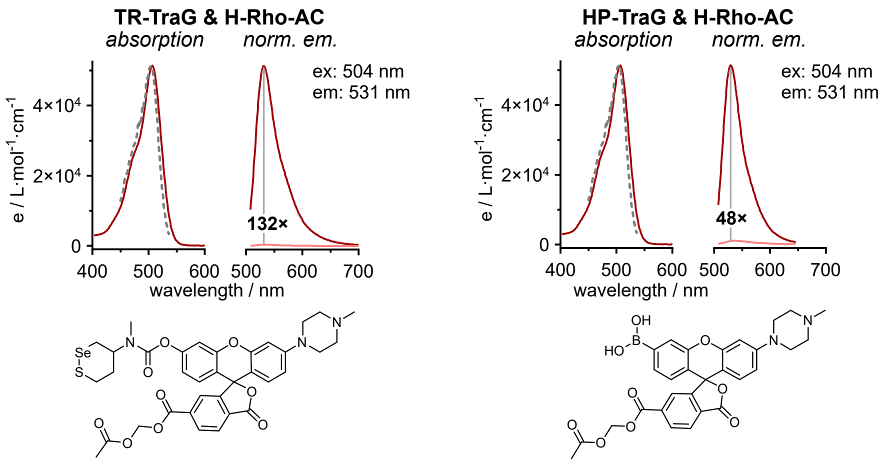

**Figure S21**: Absorption, excitation and emission spectra of **TR-TraG** (10 µM) and **HP-TraG** (5 µM) in PBS (pH = 7.4; *dark red line: <u>fluorophore</u>; light red line: <u>probe</u>; grey, dashed line represents excitation spectra: $\lambda_{em}$ = 550 nm; emission spectra: $\lambda_{ex}$ = 495 nm*).

We suppose that the lower max fluorogenicity for some of the probes derives from <2% contamination by species that can enter an open-closed equilibrium, e.g. hydrolysed or rhodamine byproducts of synthesis, that is inconvenient but also unnecessary to remove to high purity.

#### 5.1.3 Table S1: Photophysical properties overview

| | | Fluorophore | $\lambda_{Abs,max}$ / nm | $\lambda_{emission,max}$ / nm | Stokes Shift / nm | $\varepsilon_{\lambda,max}$ / L mol$^{-1}$ cm$^{-1}$ | $\Phi$ | brightness / L mol$^{-1}$ cm$^{-1}$ |
|---|---|---|---|---|---|---|---|---|
| **fluoresceins** | *two activation sites* | fluorescein | 490 | 513 | 23 | $84 \cdot 10^3$ | 0.85 | $72 \cdot 10^3$ |
| | | $H_2$-F | 503 | 523 | 20 | $68 \cdot 10^3$ | 0.78 | $53 \cdot 10^3$ |
| | | $H_2$-FS | 507 | 526 | 19 | $51 \cdot 10^3$ | 0.75 | $38 \cdot 10^3$ |
| | *single activation site* | H-C4-F | 487 | 521 | 34 | $32 \cdot 10^3$ | 0.18 | $5.7 \cdot 10^3$ |
| | | H-C4-FS | 490 | 524 | 34 | $25 \cdot 10^3$ | 0.16 | $4.0 \cdot 10^3$ |
| | | H-C7-FS | 491 | 523 | 32 | $27 \cdot 10^3$ | 0.17 | $4.7 \cdot 10^3$ |
| | | H-C10-FS | 491 | 523 | 32 | $8.4 \cdot 10^3$ | 0.09 | $0.73 \cdot 10^3$ |
| | | H-C4-FC | 489 | 521 | 32 | $33 \cdot 10^3$ | 0.21 | $7.1 \cdot 10^3$ |
| **rhodols** | *single activation site* | H-Rho* | 492* | 514 | 22 | $18 \cdot 10^3$ * | 0.26 | $4.7 \cdot 10^3$ |
| | | H-Rho-A | 503 | 530 | 27 | $52 \cdot 10^3$ | 0.64 | $33 \cdot 10^3$ |
| | | H-Rho-C | 525 | 560 | 35 | $64 \cdot 10^3$ | 0.04 | $2.6 \cdot 10^3$ |
| | | H-Rho-AC | 507 | 531 | 24 | $51 \cdot 10^3$ | 0.51 | $26 \cdot 10^3$ |

**excitation maximum instead of absorption maximum*

**Table S1: Photophysical properties of fluorescein and rhodol fluorophores.**

Quantum yields of the novel fluorophores were determined the following equation (Resch-Genger and co-workers[32]):

$$\Phi_{f,x} = \Phi_{f,st} \cdot \frac{F_x}{F_{st}} \cdot \frac{1 - 10^{-A_{st}(\lambda_{ex})}}{1 - 10^{-A_x(\lambda_{ex})}} \cdot \frac{n_x(\lambda_{em})^2}{n_{st}(\lambda_{em})^2}$$

Fluorescein was used as a reference fluorophore with a quantum yield of $\Phi_{f,st}$ = 0.85 (in PBS, pH=7.4).[33]

#### 5.2 Fig S22-S23: Cell-free probe stability in cell culture media

##### 5.2.1 Ester probes

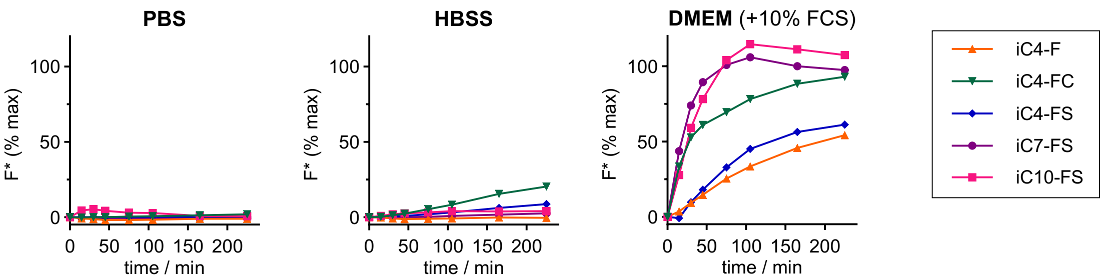

**Figure S22: Probe stability in different cell culture media.** Spontaneous probe hydrolysis time-courses of ester probes in PBS, HBSS and DMEM (probe conc.: 10 µM, incubation at 37 °C).

We examined the probe stabilities in PBS, HBSS and the standard cell culture medium DMEM (with 10% FCS). PBS is a very simple cell buffer containing sodium and potassium chloride as well as sodium hydrogen- and dihydrogen-phosphates. HBSS we used contains the same ingredients (in different amounts) plus additional salts (calcium and magnesium chloride and sulfate, sodium hydrogencarbonate), as well as glucose, to ensure longer cell viability in experiments as compared to PBS, although we expected that the stronger Lewis acids could better promote ester hydrolysis. The cell culture medium DMEM additionally contains amino acids, which we expected to give even higher ester cleavage by trans-acylation, as well as vitamins. We thus expected PBS to show the lowest spontaneous probe hydrolysis and DMEM to show the highest, while cellular viability during longer-term experiments should increase in the same order.

Indeed, in PBS we observe no relevant hydrolysis within 4 h, while in HBSS the least hydrophilic probes **iC4-FC** and **iC4-FS** show some activation (20 and 10% respectively) but overall the probes are very stable and within a typical imaging experiment timeframe (<60 min) the hydrolysis is very low. In FCS-supplemented DMEM however, all probes are rapidly activated, interestingly the lipophilic sulfonated probes (**iC10-FS** and **iC7-FS**) have the fastest turn-on, we assume that this is mainly due to assisted solubility by proteins and potential nucleophilic attack of protein surface amines (lysines). Therefore, we decided to perform all experiments with ester probes in HBSS.

##### 5.2.2 GL probes

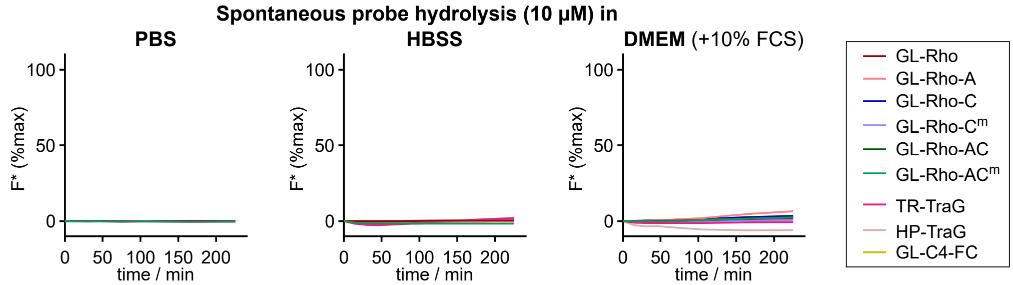

**Figure S23: Probe stability in different cell culture media.** Spontaneous probe hydrolysis time-courses of carbamate capped GL-probes in PBS, HBSS and DMEM (probe conc.: 10 µM, incubation at 37 °C).

We chose the GSH-labile GL-trigger for non-specific intracellular proof-of-concept probe activation as we expected much better hydrolytic stability of the carbamate (compared to the ester probes). Indeed, we observe no probe activation in PBS and HBSS within 4 h, and only negligible activation in DMEM (with 10% FCS). This allowed us to perform experiments with GL-probes in FCS-supplemented DMEM which is an ideal medium for cell viability.

#### 5.3 Figs S24-S25: *in vitro* activation of proof-of-concept probes

##### 5.3.1 Esterase activation of ester probes

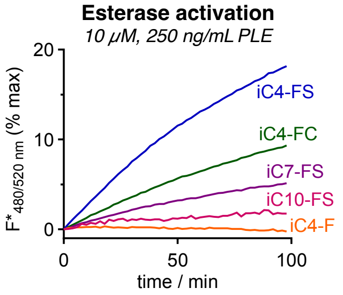

**Figure S24: Esterase activation of ester probes.** Time-course of probe activation by porcine liver esterase (250 ng/mL) above spontaneous hydrolysis in PBS (values were corrected for PBS activation, probe conc.: 10 µM, incubation at 37 °C).

The probes must be rapidly processed upon cellular entry to generate a fluorescent signal. We assessed probe activation by the model enzyme porcine liver esterase (PLE, 250 ng/mL) in PBS as the least hydrolysing buffer (**Fig S22**). All probes (except **iC4-F**) are activated above spontaneous hydrolysis in PBS proving the activation by esterases as desired. However, depending on the lipophilicity strong kinetics differences are observed: charged sulfonate **iC4-FS** with a short lipid tail gives rapid activation which is drastically slowed down for the longer lipid tails (**iC7-FS** and **iC10-FS**). Carboxylate **iC4-FC** is activated slower than is sulfonated version but still with good turn-on, uncharged **iC4-F** on the other hand shows no detectable signal generation, presumably due to insolubility and/or self-aggregation resulting in bio-unavailability.

##### 5.3.2 Glutathione activation of GL-probes

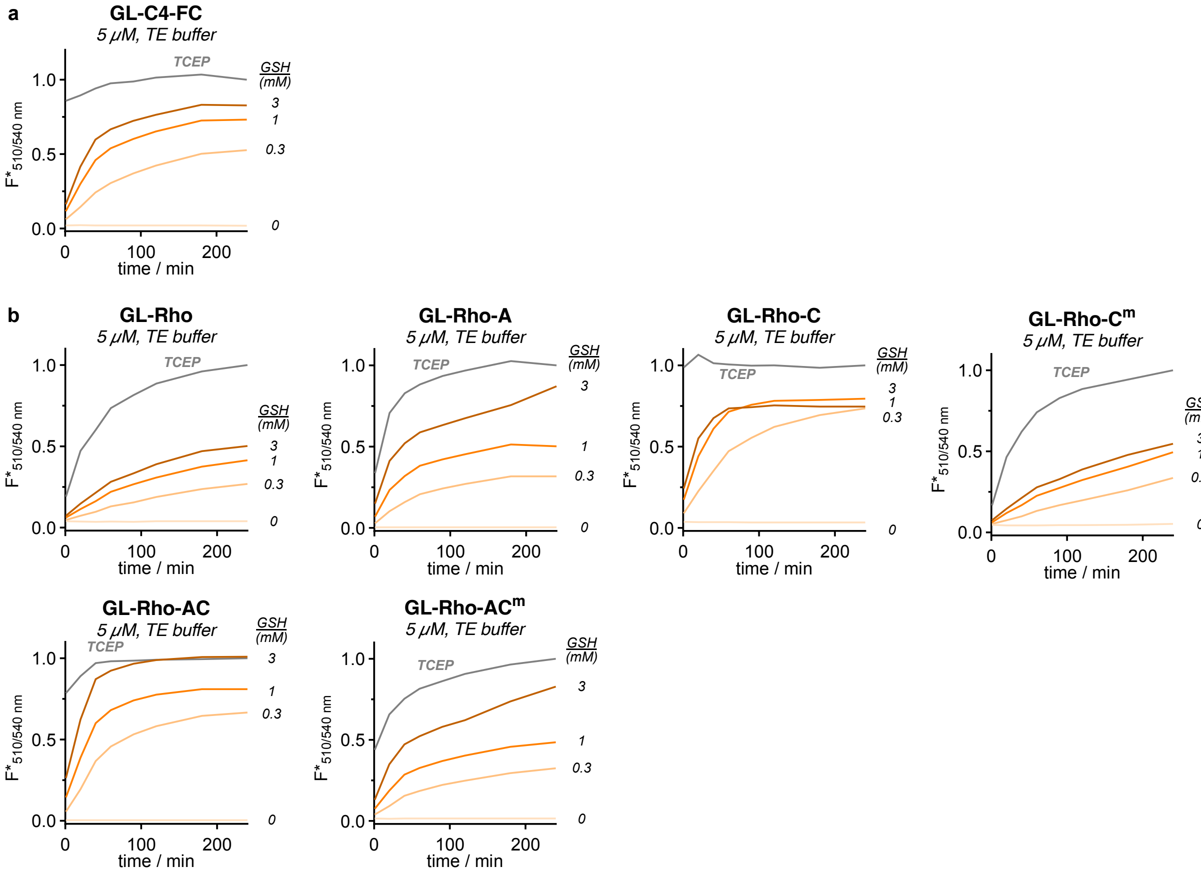

**Figure S25: Glutathione (GSH) activation of the GL-type probes.** Activation of **(a)** fluorescein-based **GL-C4-FC** and **(b)** rhodol-based probes **GL-Rho**, **GL-Rho-A**, **GL-Rho-C**, **GL-Rho-C$^m$**, **GL-Rho-AC** and **GL-Rho-AC$^m$** with GSH (0−3 mM) in TE buffer (pH = 7.4).

#### 5.4 Figs S26-S27: *in vitro* activation of HP-TraG

##### Oxidation of HP-TraG with hydrogen peroxide

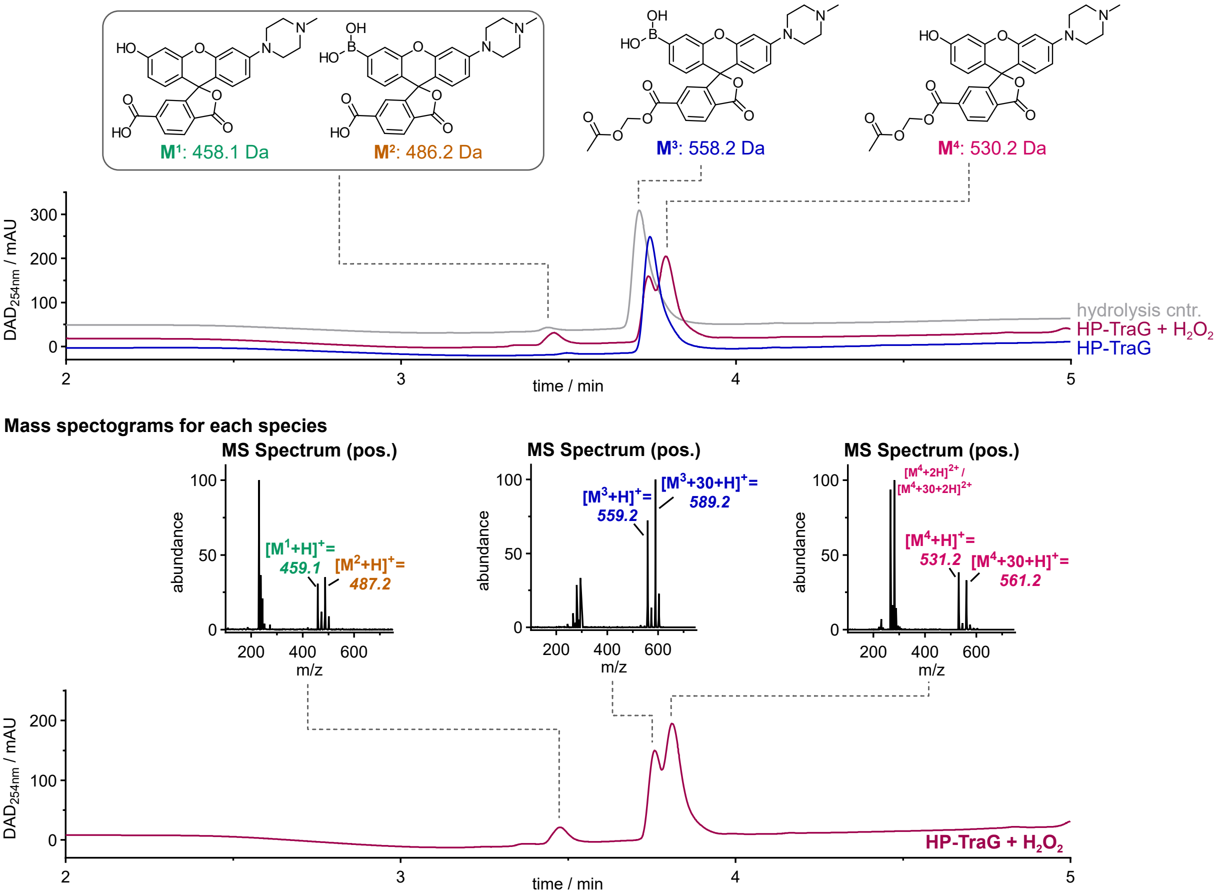

**Figure S26: Oxidation of HP-TraG with hydrogen peroxide to the corresponding rhodol** (50 µM probe with 500 µM $H_2O_2$ in PBS for 15 min at room temperature (22 °C)).

We tested the $H_2O_2$ sensing ability of **HP-TraG** by HPLC-MS to confirm the desired oxidation pathway. **HP-TraG** (**Fig S26**, molecule **M**$^3$, 50 µM) was treated with 10 eq $H_2O_2$ (0.5 mM) in PBS and incubated at room temperature for 15 min and then analysed by HPLC-MS. **HP-TraG** is well oxidised to the corresponding rhodol (**M**$^4$) with ca. 60% conversion while a small proportion of the acetoxymethyl esters of both **HP-TraG** and **M**$^4$ hydrolyse presumably due to $H_2O_2$ nucleophilicity, to give **M**$^1$ and **M**$^2$. This **HP-TraG** sample contained an extra, co-eluting species with an additional formaldehyde unit (m/z = x+30) which was still present in the **M**$^4$ oxidation product, proving that the modification was not located on the boronic acid, and it did not change the functional performance for oxidation-induced fluorogenicity (see below).

##### Acetoxymethyl ester cleavage by esterase (PLE)

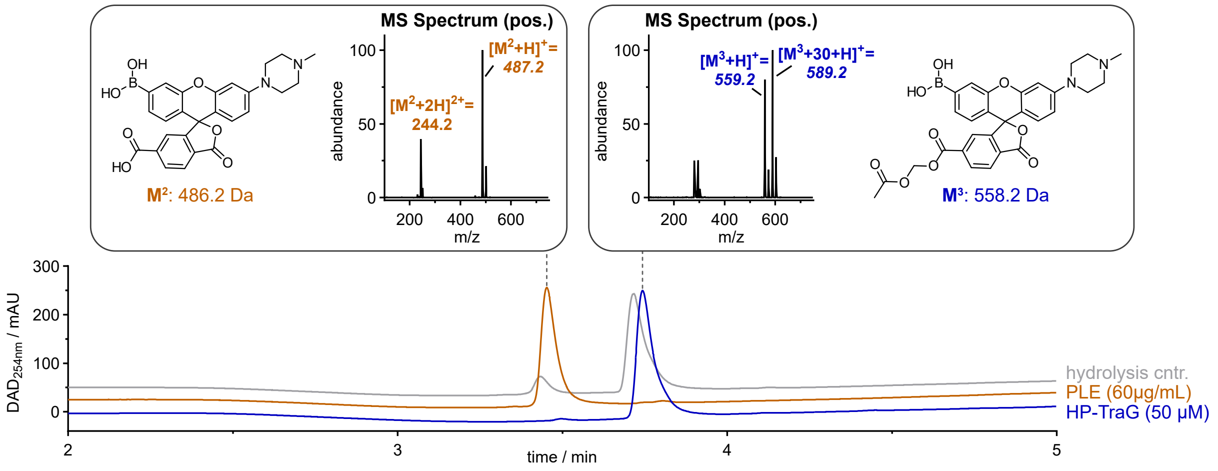

**Figure S27: Acetoxymethyl ester cleavage of HP-TraG with porcine liver esterase (PLE)** (50 µM probe with 60 µg/mL PLE at 37 °C for 1.5 h.

We then investigated the esterase unmasking of the acetoxymethyl ester which is required for reliable intracellular trapping of the fluorophore (**Fig S27**). **HP-TraG** (molecule **M**$^3$, 50 µM) was incubated with porcine liver esterase (PLE, 60 µg/mL, 15 U/mg) at 37 °C for 1.5 h which cleanly converted **HP-TraG** (as well as the M+30 impurity) into the same carboxylate **M**$^2$ with no +30 shift in the mass spectrum. This shows that the extra formaldehyde unit in the **HP-TraG** side product is located on the pendant carboxylate, and is converted to the same boronic acid product (**M**$^2$) by intracellular esterases. Taken together, the *in vitro* hydrogen peroxide oxidation and the esterase catalysed ester cleavage assays show that **HP-TraG** follows the expected pathway for oxidation to the phenol, and that the co-eluting contaminant which arises during synthesis is likely to have the same cellular performance (ester cleavage to give the same cell-trapped boronate which is the active sensing species). We therefore used the mixtures with variable residual amounts of M+30 contaminant for the cellular evaluations since separation (C18 and normal-phase column chromatography) was infeasible and since the outcomes were identical.

### 6 Biological materials and methods

#### 6.1 Cell culture and cell lines

Cells were grown in high glucose Dulbecco's modified Eagle's medium (DMEM, Sigma-Aldrich, D1145) supplemented with 10% heat-inactivated fetal bovine serum (Biochrom S0615), 1% L-glutamine (Sigma-Aldrich, G7513), 1 mM Sodium Pyruvate (Sigma-Aldrich, S8636) and 100 nM sodium selenite (Sigma-Aldrich, 214485-5G) at 37 °C and 5% $CO_2$. Washing was performed with Dulbecco's phosphate buffered saline (PBS, Sigma-Aldrich, D8537), cell detachment was performed using Trypsin-EDTA solution (Sigma-Aldrich, T4174) diluted to 1× Dulbecco's PBS (Sigma-Aldrich, D8537). Cell growth was monitored using an inverted microscope (Nikon Eclipse Ti), passage was kept between 2 and 20.

HeLa (DSMZ Cat No. ACC57) and A549 (DSMZ; ACC 107) cells were obtained from German Collection of Microorganisms and Cell Cultures. HEK293T cells were obtained from ATCC (Cat. No. crl-3216), TrxR knockout and reference mouse embryonic fibroblasts (MEF) were a kind gift from Marcus Conrad. All cell lines were tested regularly for mycoplasma contamination and only mycoplasma negative cells were used in assays.

In all experiments, probes and inhibitors were treated from DMSO stocks with 1% final DMSO concentration in the experiment (unless stated otherwise).

**Cell media**

- Phosphate buffered saline (PBS) from Sigma Aldrich (Cat. No. D8537)
- Hanks´ balanced salt solution (HBSS) from Sigma Aldrich (Cat. No. H6648)
- Dulbecco´s modified eagle´s medium (DMEM) from Sigma Aldrich (Cat. No. D1145), used with the following supplements (unless stated otherwise): 10% FCS (Biochrom S0615), 1 mM Sodium Pyruvate (Sigma-Aldrich, S8636) and 100 nM sodium selenite (Sigma-Aldrich,214485-5G)

#### 6.2 Cell-free characterisation

Whole well fluorescence measurements were performed with a BMG Labtech FluoStar Omega plate reader (fluorescein settings: ex: 485bp10 and em: 520lp) or a Tecan M1000 plate reader: rhodol settings: excitation 510 nm, emission 540 nm; fluorescein settings: excitation 498 nm, emission 518 nm. All plate reader experiments were performed in coated black F-bottom 96-well plates (greiner, 655077).

**Probe stability assay**

Water (10 µL) was placed in a black 96 well plate and the probes/fluorophores were added from a 1 mM stock solution in DMSO (1 µL). Then the different media PBS, HBSS or DMEM (89 µL) were added to the probes with a multipipette (for smallest possible differences in the starting time) to reach a final concentration of 10 µM. All samples in technical duplicates. The fluorescence intensity in each well was measured after 1, 15, 30 min and 1, 2, 3, 4, 5, 6 h using a Tecan Infinite M1000 plate reader (fluoresceins: ex/em 480bp5/520bp5; rhodols: ex/em 510bp5/540bp5). Between measurements the samples were incubated at 37 °C under air atmosphere. At the end of the experiment, a *n*-butyl amine solution (50 mM, 100 µL) was added to each well to determine the maximum fluorescence after full probe activation in each well. The recorded data were background subtracted (fluorescence at 0 min) and normalised to their maximum fluorescence (determined after 2.5 h incubation with *n*-butylamine, corrected for concentration and pH changes by calculating the fluorescence ratio for the fluorophores upon *n*-butylamine addition).

**Esterase activation of fluorescein esters**

PBS (49 µL) was placed in a black 96 well plate and the probes/fluorophores were added from a 1 mM stock solution in DMSO (1 µL). Porcine liver esterase (lyophilized powder, ≥15 units/mg, purchased from Sigma

Aldrich, Cat.-No.: E3019) was diluted in PBS to give a stock solution of 500 ng/mL and warmed to 37 °C. The esterase stock solution was added to the probes with a multipipette (for smallest possible differences in the starting time) to reach a final probe/fluorophore concentration of 10 µM and an esterase concentration of 250 ng/mL (all samples in technical triplicates). The probes/fluorophores were placed in the pre-heated plate reader and incubated at 37 °C for the whole experiment. The fluorescence intensity in each well was measured every 2 min for the total duration of 100 min. At the end of the experiment, a *n*-butylamine solution (50 mM, 100 µL) was added to each well to determine the maximum fluorescence after full probe activation in each well. The recorded data were background subtracted (fluorescence at 0 min) and normalised to their maximum fluorescence (determined after 2.5 h incubation with *n*-butylamine, corrected for concentration and pH changes by calculating the fluorescence ratio for the fluorophores upon *n*-butylamine addition).

**GSH activation of GL probes**

The probes were placed in the respective wells of a 96-well plate (50 µL, 10 µM in TE buffer, final conc. 5 µM), then GSH (50 µL, 2× stock solutions in TE buffer, final conc 0, 0.3, 1.0, 3.0 mM) or TCEP (50 µL, 0.6 mM in TE buffer, final conc 0.3 mM; positive control for maximum activation) was added with a multipipette (for smallest possible differences in the starting time). The fluorescence intensity in each well was measured after 1, 20, 40, 60, 90, 120, 180 and 240 min using a Tecan Infinite M1000 plate reader (GL-C4-FC: ex/em 480bp5/520bp5; rhodols: ex/em 510bp5/540bp5). Between measurements the samples were incubated at 37 °C under air atmosphere. The recorded data were normalised to their maximum fluorescence (determined at the endpoint of the TCEP positive control).

#### 6.3 Cellular characterisation

**General**

Whole well fluorescence measurements were performed with a FluoStar Omega plate reader from BMG Labtech, Ortenburg (Germany) (fluorescein settings: ex: 485bp10 and em: 520lp) or on a M1000 plate reader (Tecan): rhodol settings: excitation 510 nm, emission 540 nm; fluorescein settings: excitation 498 nm, emission 518 nm. All plate reader experiments were performed in coated black F-bottom 96-well plates (greiner, 655077). The data was assembled and sorted in Microsoft Excel (version 16.88) and plotted in GraphPad Prism (version 10.4.1).

Epifluorescence microscopy images were acquired on an inverted microscope (Nikon Eclipse Ti), using the CFI Plan Achro DL 10× objective (Nikon). For fluorescence images the probes were excited with a pE-4000 (CoolLED) (rhodols: ex. 490 nm at 100% intensity with 530/50 nm emission filter; RX1: ex. 365 nm with 420lp emission filter).

##### 6.3.1 Proof-of-concept probes: uptake and retention

**Comparison of FC-probe uptake in DMEM and HBSS**

We compared the cellular signal generation of **GL-C4-FC** in HBSS and DMEM. HEK293T cells (10.000 cells/well) were seeded on Poly-D-Lysine (gibco, A3890401) coated clear 96-well plates (TPP, Z707902) in 100 µL. After 1 d the cells were washed (3× medium change with DMEM or HBSS respectively) and **GL-C4-FC** and **iC4-FC** were added at 1, 3 and 10 µM from 10× stock solutions using a D300e Digital Dispenser (Tecan). The probes were incubated for 30 min and imaged by epifluorescence microscopy. During incubation, cells in HBSS were kept at 37 °C under air atmosphere and cells in DMEM were kept at 37 °C under 5% $CO_2$ atmosphere.

**Uptake and post-wash retention (plate reader)**

Fluorescence measurements were performed on a M1000 plate reader (Tecan) (rhodol settings: excitation 510 nm, emission 540 nm; fluorescein settings: excitation 498 nm, emission 518 nm).

HEK293T cells (10.000 cells/well) were seeded on Poly-D-Lysine (gibco, A3890401) coated black F-bottom 96-well plates (greiner, 655077). After 1 d the medium was removed and the rhodol probes (GL-Rho-AC$^m$, GL-Rho, GL-Rho-A, GL-Rho-AC, GL-Rho-C$^m$, GL-Rho-C) and fluorescein probes (GL-MF) were added in DMEM (100 µL, probe conc. 10 µM) with a multipipette (for smallest possible differences in the starting time).

The samples were incubated at 37 °C for 30 min under 5% $CO_2$ and the fluorescence was measured ("pre-wash"). The cells were washed (2× medium change) and the fluorescence was measured ("0 min post-wash"). The cells were incubated at 37 °C for 60 min under 5% $CO_2$ and the cells were washed (2×) and measured after 20 and 60 min ("20/60 min post-wash" respectively).

*Controls*:
(1) Untreated cells for subtraction of the autofluorescence (of cells and medium)
(2) Probes in cell medium for subtraction of the residual probe fluorescence in DMEM (10 µM probe conc.)
(3) Full activation of probes for determination of maximum probe fluorescence: TCEP (300 µM final concentration, BLDpharm, BD155793) was added to the probes in DMEM (5 µM final probe conc.) and the fluorescence was measured after 90 min (when the signal was constant)

The raw data was assembled and sorted using Microsoft Excel. "Cell entry & activation" values were calculated from "pre-wash" fluorescence subtracted by the residual fluorescence of probes in cell medium (Control 2) which was divided by the max. fluorescence of fully activated probe (Control 3). "Cell retention" values were calculated from cellular post-wash fluorescence subtracted by the cellular autofluorescence (Control 1) which was divided by the total amount of activated probe ("pre-wash" fluorescence minus control 2). Data was plotted using GraphPad Prism. For all resulting plots, one data point represents one biological replicate.

##### Confocal microscopy imaging

*Fluorescein probes*

Instrument: Leica SP8 point scanning confocal microscope using a 20× air objective, at 2× zoom, 1024x1024 pixel, 200 Hz scan speed, Pinhole: 1.00 AU. Environmental conditions were controlled using a stage top incubator (OKO Lab H301 K-frame) at 37 °C without $CO_2$ (for HBSS cell medium). The fluoresceins were excited at 488 nm (10% intensity), and emission was detected at 504-548 nm using a HyD detector. Cell Tracker Red CMTPX was excited with 594 nm (1% intensity) and emission was detected at 605−645 nm using a HyD detector. All probes were imaged in 2−3 independent biological runs.

HEK293T cells were plated on poly-D-lysine coated (gibco, A3890401) 8-well imaging dishes (No. 1.5, µ-Slide, 8-well, ibidiTreat, ibidi USA Inc., Wisconsin) and left to adhere in DMEM for 16-24 h. The DMEM was removed and warm HBSS was added. Cells were kept at 37 °C without $CO_2$ atmosphere and treated with Cell Tracker Red (CMTPX, Invitrogen, C34552, final conc. 1 µM) in HBSS for 30 min at 37 °C. The cells were washed with warm HBSS (2× medium change), then the probes were added (2 µL from 200× stock into 400 µL HBSS on the cells, final probe conc. 5 µM). Pre-wash images were acquired after 10, 20 and 30 min, then the cells were washed with warm HBSS (3× medium change), and post-wash images were acquired every 10 min for 60 min.

Quantification of fluorescence intensity was performed using Fiji/imageJ. All cells were selected from raw, unprocessed images based on the transmitted light image and the fluorescence intensity of the probe channel was measured. All cells were selected, overlapping areas were counted only for one of the cells. Typically, more than 50 cells were selected for each compound per time point over three locations within the same well.

*Rhodol probes (except HP-TraG)*

Instrument: Leica SP8 point scanning confocal microscope using a water 40× objective was used, no digital zoom, 2048x2048 pixels, 200 Hz scan speed, Pinhole: 1.00 AU. Environmental conditions were controlled using a stage top incubator (OKO Lab H301 K-frame) at 37 °C with 5% $CO_2$. The rhodols were excited at 518 nm (0.2% intensity), and emission was detected at 523−593 nm using a HyD detector. Cell Tracker Red CMTPX was excited with 598 nm (0.5% intensity) and emission was detected at 603−789 nm using a HyD detector. All probes were imaged in three independent experiments ($n$ = 3).

HEK293T cells were seeded at 40.000 cells/well in Poly-D-Lysine coated (gibco, A3890401) 8-well imaging dishes (No. 1.5, µ-Slide, 8-well, ibidiTreat, ibidi USA Inc., Wisconsin) in 200 µL DMEM and cultured over night. The cells were stained with CellTracker Red (CMTPX, 4 µM, Invitrogen A3890401) for 45 min. The cells were washed (2× medium change), treated with the rhodol probe (5 µM, final DMSO concentration 0.5%) for 30 min and imaged to get pre-wash images. The cells were washed to remove the extracellular probe (2× medium change) and imaged after 1, 10, 20, 30 and 60 min.

For the cellular signal distribution of **GL-Rho-AC$^m$**, representative images were taken after 30 min of probe incubation (5 µM) using a 63× oil objective with otherwise identical conditions as above.

Images were analysed using Fiji ImageJ. For cellular fluorescence quantification, images were segmented by thresholding on the CellTracker channel. The resulting mask was then applied to the probe channel to obtain a mean fluorescence intensity value. Data was plotted using GraphPad Prism.

##### Retention in different cell lines

Experiments were performed in high glucose DMEM. HeLa (10.000 cells/well), MEF (5.000 cells/well) and A549 (5.000 cells/well) cells were seeded on Poly-D-Lysine (gibco, A3890401) coated black, clear F-bottom 96-well-plates (greiner, 655096) in 100 µL medium. After 1 d the medium was removed and the compounds **GL-Rho-AC$^m$** and **GL-Rho** were added in DMEM (5 µM probe concentration), incubated for 30 min and pre-wash images were acquired by epifluorescence microscopy. Then the cells were washed (2× medium change with DMEM) and imaged after 1, 20, 60 and 120 min while keeping the cells at 37 °C under $CO_2$ atmosphere between measurements.

##### 6.3.2 TR-TraG characterisation

**TRi-1 inhibition assay**

The activation of **TR-TraG** was quantitatively assessed depending on TrxR inhibition with Tri-1 by microplate reader fluorescence measurement. HeLa and A549 cells were seeded at 20.000 cells/well on 96-well plates (microplates, 96-well, F-bottom, µCLEAR®, black, Fluotrack, high binding; Greiner bio-one GmbH, 655087) in 100 µL medium (DMEM, 10% FCS) and cultured over night. Then the cells were preincubated with TRi-1 (2, 4, 8 and 16 µM final concentrations from DMSO stock solutions) for 3 h, cells were washed with PBS (2×) and the probe (**TR-TraG**: 25 µM; RX1: 100 µM) was added in fresh DMEM. The fluorescence intensity was measured hourly for 6 h using a Tecan Infinite M1000 plate reader (**TR-TraG**: ex/em 510bp5/540bp5; RX1: ex/em 355bp5/520bp5). In between measurements, cells were kept at 37 °C and 5% $CO_2$ atmosphere.

**Probe dose dependent activation**

The activation of **TR-TraG** and RX1 was quantitatively assessed depending on the probe concentration by microplate reader fluorescence measurement. HeLa cells were seeded at 20.000 cells/well on 96-well plates (microplates, 96-well, F-bottom, µCLEAR®, black, Fluotrack, high binding; Greiner bio-one GmbH, 655087) in 100 µL medium (DMEM, 10% FCS) and cultured over night. Then the cells were treated with **TR-TraG** at 3, 10 and 25 µM or RX1 at 25, 50 and 100 µM (1% final DMSO concentration). The fluorescence intensity was measured hourly for 6 h using a Tecan Infinite M1000 plate reader (**TR-TraG**: ex/em 510bp5/540bp5; RX1: ex/em 355bp5/520bp5). In between measurements, cells were kept at 37 °C and 5% $CO_2$ atmosphere.

**Cell-free enzyme assays**

All experiments were performed in TE buffer (Tris-HCl (50 mM), EDTA (1 mM), pH = 7.4). and in black 96-well plates (microplates, 96-well, F-bottom, black, Fluotrack, high binding; Greiner bio-one GmbH, 655077) in technical triplicates. For all measurements the fluorescence intensity was measured every 150 s at 37 °C for 4 h using a Tecan Infinite M1000 plate reader (**TR-TraG**: ex/em 510bp5/540bp5).

*TrxR1*

TrxR1 (40 µL from 2.5× stock in TE buffer, final conc. 1−100 nM) was placed in the respective wells, **TR-TraG** (50 µL from 2× stock in TE buffer, final conc. 10 µM) was added and the reaction was started by adding NADPH (10 µL from 1 mM stock in TE buffer, final conc. 100 µM).

*Trx1*

Trx1 (20 µL from 5× stock in TE buffer, final conc. 0.01−10 µM) and TrxR1 (20 µL from 5× stock in TE buffer, final conc. 20 nM) were placed in the respective wells, **TR-TraG** (50 µL from 2× stock in TE buffer, final conc. 10 µM) was added and the reaction was started by adding NADPH (10 µL from 10× stock in TE buffer, final conc. 100 µM).

*Grx1*

Grx1 (20 µL from 5× stock in TE buffer, final conc. 0.01−10 µM), GR (10 µL from 5× stock in TE buffer, final conc. 10 nM) and GSH (10 µL from 5× stock in TE buffer, final conc. 100 µM) were placed in the respective wells, **TR-TraG** (50 µL from 2× stock in TE buffer, final conc. 10 µM) was added and the reaction was started by adding NADPH (10 µL from 10× stock in TE buffer, final conc. 100 µM).

*Controls*: The following negative controls were also measured: NADPH only (100 µL NADPH at 100 µM in TE-buffer), NADPH with **TR-TraG** (100 µM NADPH with 10 µM **TR-TraG** in TE-buffer), **TR-TraG** only (10 µM in TE-buffer), **TR-TraG** with Trx1 and Grx1 without the respective reductases (10 µM Trx1/Grx1 with 10 µM **TR-TraG** in TE-buffer) and **TR-TraG** with GSH (0.1−10 mM GSH with 10 µM **TR-TraG** in TE-buffer). To determine the maximum probe fluorescence, TCEP (final concentration of 200 µM) was added to the probes.

Human recombinant thioredoxin 1 (Trx 1) (lyophilized), human recombinant glutaredoxin 1 (Grx 1) (lyophilized from 10 µL TE-buffer, pH 7.5), human thioredoxin reductase (TrxR1) (1.5 mg/mL in 50% glycerol/TE-buffer, pH 7.5) and baker's yeast glutathione reductase (GR) (100 µM in 50% glycerol/TE-buffer, pH 7.5) were a kind gift by Elias Arnér and produced as previously described.[34,35]

**Comparison of TR-TraG and RX1 (microcopy)**

To qualitatively evaluate the effects of the **TR-TraG** and RX1 regarding signal generation and cell toxicity we imaged both probes by epifluorescence microscopy ((**TR-TraG**: ex/em 490bp5/510lp; RX1: ex/em 365bp5/410lp). HeLa cells were seeded at 20.000 cells/well on 96-well plates (microplates, 96-well, F-bottom, µCLEAR®, black, Fluotrack, high binding; Greiner bio-one GmbH, 655087) in 100 µL medium. Cells were treated with **TR-TraG** (10 µM) and RX1 (100 µM) and imaged after 45 min, 1.5 h, 3 h and 6 h. After 6 h of incubation, the were washed (2× medium change) and imaged again after 24 h.

##### 6.3.3 HP-TraG characterisation

**General**

All cells were grown at 37 °C and 5% $CO_2$. Cell growth was monitored using an inverted microscope (Leica DMi1). HEK 293T cells were grown in DMEM (ThermoFisher 21885108) supplemented with 10% FBS (Biochrom S0615) and 1% Penicillin/Streptomycin (ThermoFisher 15140122). Hoxb8 cells were grown in RPMI (Thermo Fisher 61870010) supplemented with 10% FBS (Biochrom S0615), 1% Penicillin/Streptomycin (ThermoFisher 15140122), 0.1% 2-Mercaptoethanol (Thermo Fisher 31350010), 1 µM β-estradiol (Sigma E2758), and supernatant from a Flt3L-producing B16 melanoma cell line to a final concentration of 35 ng/mL. For macrophage differentiation, Hoxb8 cells were grown in RPMI supplemented with 10% FBS, 1% Penicillin/Streptomycin and 10-20 ng/ml M-CSF (PeproTech 315-02) for 5 days before seeding.

Confocal live cell imaging was performed at the Core Facility Bioimaging of the Biomedical Center with an inverted Leica SP8X microscope, equipped with Argon laser, WLL2 laser (470−670 nm) and acusto-optical beam splitter. Live cells were treated and recorded at 37 °C. For the duration of the assay, cells were kept in Hank’s Balanced Salt Solution (HBSS, ThermoFisher 14025092) to allow incubation without $CO_2$. Assays were performed in 8-well glass bottom chambered coverslips (ibidi 80827).

The microscope was programmed to take three images of different fields of view per condition and time point tested, which were focused using reflection-based adaptive focus control. Images were acquired with a 20× 0.75 objective and additional 2× optical zoom. Image pixel size was 284 nm. The following fluorescence settings were used: **HP-TraG** excitation 488 nm (Argon), emission 500−570 nm; CellTracker Red excitation 594 nm (WLL), emission 605−645 nm. Recording was performed sequentially to avoid bleed-through. **HP-TraG** and CellTracker Red were recorded with hybrid photo detectors (HyDs), a transmitted light image was generated with a conventional photomultiplier tube.

Images were analysed using Fiji ImageJ. For cellular fluorescence quantification, images were segmented by thresholding on the CellTracker channel. The resulting mask was then applied to the **HP-TraG** channel to obtain a mean fluorescence intensity value. Data was plotted using GraphPad Prism.

**$H_2O_2$ assays with HEK cells**

Chambered coverslips were coated 2 d before the experiment by applying a 0.1 mg/mL Poly-D-Lysine (Sigma-Aldrich P7280) solution for 2 h at 37 °C. HEK cells were seeded into the coated wells at a density of 20.000 cells per cm².

Cells were stained with 0.5 µM CellTracker™ Red CMTPX Dye (ThermoFisher C34552) at 37 °C for 15 min. The medium was then changed to 10 µM **HP-TraG** solution in HBSS. After 10 min loading time, pre-treatment images were recorded. After 15 min loading time, cells were washed with HBSS once and the medium then changed once more to $H_2O_2$ solutions of the indicated molarities in HBSS or HBSS only for the negative control wells. The different $H_2O_2$ concentrations were obtained by serial dilution. Images were acquired at 15, 30 and 60 min of treatment. All wells were then washed twice with HBSS and post-wash images acquired at the indicated times thereafter.

**PMA assays with Hoxb8-derived macrophages**

Chambered coverslips were coated one day before experiment by applying a 0.1 mg/ml Poly-D-Lysine (Sigma-Aldrich P7280) solution for 2 h at 37 °C. Hoxb8-derived macrophages were seeded into the coated wells at a density of 300.000 cells per cm² and their differentiation medium was additionally supplemented with TNF-α (10 µg/ml, PeproTech 315-01A) and IFN-γ (10 µg/mL, PeproTech 315-05) for M1 polarisation.

Cells were stained with 0.5 µM CellTracker™ Red CMTPX Dye (ThermoFisher C34552) at 37 °C for 15 min. The medium was then changed to 10 µM **HP-TraG** solution in HBSS. After 10 min loading time, pre-treatment images were recorded. After 15 min loading time, cells were washed with HBSS once and the medium then changed once more to 1 µg/ml PMA in HBSS or HBSS only for the negative control wells. Images were acquired at 15, 30 and 60 min of treatment. All wells were then washed twice with HBSS and post-wash images acquired at the indicated times thereafter.

### 7 Synthetic Chemistry

#### 7.1 Chemistry methods and techniques

##### 7.1.1 Analytical methods

High resolution mass spectrometry (**HRMS**) was conducted on the following instruments: (1) a *Thermo Finnigan LTQ FT Ultra FourierTransform* ion cyclotron resonance spectrometer from *ThermoFisher Scientific GmbH* applying electron spray ionisation (ESI) with a spray capillary voltage of 4 kV at temperature 250 °C with a method dependent range from 50 to 2000 u;(2) a *Finnigan MAT 95 from Thermo Fisher Scientific* applying electron ionisation (EI) at a source temperature of 250 °C and an electron energy of 70 eV with a method dependent range from 40 to 1040 u; and (3) a Waters Xevo G2-XS Q-TOF applying electron spray ionisation (ESI) with a spray capillary voltage of 2 kV at a source temperature of 140 °C with a method dependent range from 50 to 1200 u.

Nuclear magnetic resonance (**NMR**) spectroscopy was performed using the following instruments: (1) a *Bruker Avance* (600/150 MHz, with TCI cryoprobe) or (2) a *Bruker Avance III HD Biospin* (400/100 MHz, with BBFO cryoprobe™) from Bruker Corp. or (3) a Bruker Avance III HD (800 MHz, with cryoprobe) or (4) a Bruker Avance Neo (600/150 MHz, with cryoprobe). NMR-spectra were measured at 298 K, unless stated otherwise, and were analysed with the program *MestreNova 12* developed by *MestreLab Ltd.* $^{1}$H-NMR spectra chemical shifts (δ) in parts per million (ppm) relative to tetramethylsilane (δ = 0 ppm) are reported using the residual protic solvent ($CHCl_3$ in $CDCl_3$ : δ = 7.26 ppm, DMSO-$d_5$ in DMSO-$d_6$: δ = 2.50 ppm, $CHD_2OD$ in $CD_3OD$: δ = 3.31 ppm) as an internal reference. For $^{13}$C-NMR spectra, chemical shifts in ppm relative to tetramethylsilane (δ = 0 ppm) are reported using the central resonance of the solvent signal ($CDCl_3$: δ = 77.16 ppm, DMSO-$d_6$: δ = 39.52 ppm, $CD_3OD$: δ = 49.00 ppm) as an internal reference. For $^{1}$H-NMR spectra in addition to the chemical shift the following data is reported in parenthesis: multiplicity, coupling constant(s) and number of hydrogen atoms. The abbreviations for multiplicities and related descriptors are s = singlet, d = doublet, t = triplet, q = quartet, or combinations thereof, m = multiplet and br = broad. When rotamers were observed in the NMR spectra, the corresponding signals are separated by a slash ("/"). Where known products matched literature analysis data, only selected data acquired are reported.

Analytical high performance liquid chromatography (**HPLC**) analysis was conducted either using an *Agilent 1100* system from *Agilent Technologies Corp.,* Santa Clara (USA) equipped with a DAD detector and a *Hypersil Gold* HPLC column from *ThermoFisher Scientific GmbH*, Dreieich (Germany) or a *Agilent 1200 SL* system *Agilent Technologies Corp.,* Santa Clara (USA) equipped with a DAD detector, a *Hypersil Gold* HPLC column from *ThermoFisher Scientific GmbH*, Dreieich (Germany) and consecutive low-resolution mass detection using a LC/MSD IQ mass spectrometer applying ESI from *Agilent Technologies Corp.,* Santa Clara (USA). For both systems mixtures of water (analytical grade, 0.1 % formic acid) and MeCN (analytical grade, 0.1 % formic acid) were used as eluent systems.

**UV-Vis** spectra were recorded on an Cary 60 UV-Vis spectrophotometer from *Agilent Technologies Inc.*, Santa Clara (USA) using 1 cm quartz or PMMA cuvettes. The scan rate was set to 600 nm/min and 2.5 nm slit width was used. Unless stated otherwise, the probes and fluorophores were dissolved in PBS (pH = 7.4, 1 % DMSO) at 10 µM concentration.

**Fluorescence spectroscopy** was performed on a Cary Eclipse Fluorescence Spectrometer from *Agilent Technologies Inc.*, Santa Clara (USA) using quartz cuvettes (scan rate: 120 nm/min, 5 nm slit width) or on a Tecan Infinite M1000 plate reader. Unless stated otherwise, the samples were measured at 10 µM concentration.

##### 7.1.2 Synthetic techniques

Unless stated otherwise, all reactions were performed without precautions regarding potential air- and moisture-sensitivity and were stirred with Teflon-coated magnetic stir bars. For work under inert gas (nitrogen) atmosphere, a Schlenk apparatus and a high vacuum pump from Vacuubrand GmbH, Wertheim (Germany) were used. For solvent evaporation a *Laborota 400* from Heidolph GmbH, Schwabach (Germany) equipped with a vacuum pump was used. Flash column chromatography was conducted with a Biotage® Isolera One Chromatograph with Biotage® Sfär Silica D columns (10 g or 25 g silica) for normal-phase (np) chromatography or with Biotage® Sfär C18 D columns (12 g or 30 g silica) for reversed-phase (rp) chromatography. Reactions were monitored by thin layer chromatography (TLC) on TLC plates (*Si 60 F254 on aluminium sheets*) provided by Merck GmbH and visualised by UV irradiation and by analytical HPLC-MS. The procedures and yields are not optimised.

##### 7.1.3 Chemicals

All chemicals, which were obtained from BLDpharm, Sigma-Aldrich, TCI, Alfa Aesar, Acros, abcr or carbolution were used as received and without purification. Tetrahydrofuran (THF), dichloromethane (DCM) and dimethylformamide (DMF) were provided by Acros and were stored under argon atmosphere and dried over molecular sieves. TLC control, extractions and column chromatography were conducted using distilled, technical grade solvents. Whenever the term *hexanes (Hex)* is used, the applied solvent actually comprised isomeric mixtures of hexane (2-methylpentane, 3-methylpentane, 2,2-dimethylbutane, 2,3-dimethylbutane).

#### 7.2 Synthetic procedures

##### 7.2.1 Literature procedures

The following molecules were synthesised according to literature procedures:

- **$i_2$-F** (Chyan *et al.*[23], compound 3b)
- **$i_2$-FS** (Mauker *et al.*[22], compound $i_2$-$FS_1$)
- **iPS-F** (Mauker *et al.*[22], compound iPS-$FS_1$)
- **GL-MF** (Zeisel, Felber *et al.*[36], compound Ac-SS66C-MF)
- **Compound 2** (Zeisel, Felber *et al.*[36], compound 7T)

##### 7.2.2 General Procedures

**General Procedure A: *O´*-alkyl fluorescein ester probes**

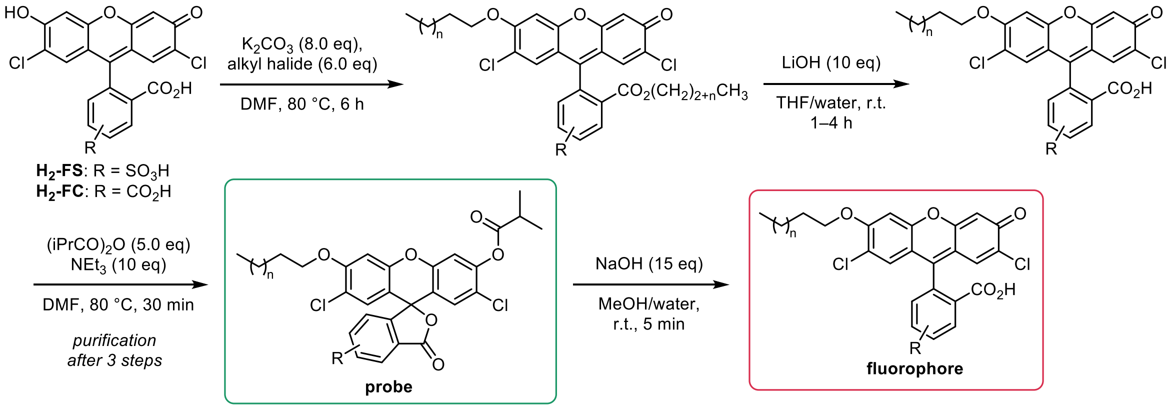

<u>*Step 1: Alkylation*</u>
**$H_2$-FS** or **$H_2$-FC** (1.0 eq) was dissolved in anhydrous DMF (0.02−0.15 M). Potassium carbonate (8.0 eq) and the respective **alkyl halide** (6.0 eq) were added and the reaction was heated 80 °C for 2−15 h until full conversion to the di-alkyl product (for sulfofluoresceins) or to the tri-alkyl product (for carboxyfluoresceins) (monitored by HPLC/MS). The reaction mixture was filtered to remove insoluble salts and the solvent was removed *in vacuo*.

<u>*Step 2: Ester cleavage*</u>
The crude product was dissolved in THF/water (0.004−0.05 M) (*Note: at high salt/compound concentrations THF and water are not miscible*). Lithium hydroxide (10 eq) was added and the reaction mixture was stirred at room temperature for 1−4 h. Aqueous hydrochloric acid (2 M, 12 eq) was added to acidify the solution (checked by pH paper) and the volatiles were removed *in vacuo* and optionally semi-purified by reversed-phase flash column chromatography affording the *O´*-alkylated fluorophore (*Note: acidification before evaporation is recommended to avoid decomposition of the fluorescein to the corresponding benzophenones*).

<u>*Step 3: Acylation*</u>
The crude fluorophore was dissolved in DMF (0.01−0.05 M), triethylamine (10 eq) and isobutyric anhydride (5.0 eq) were added and the reaction mixture was heated to 80 °C for 30 min. The volatiles were removed *in vacuo* and the crude product was purified by reversed-phase flash column chromatography (acetonitrile/water, 0.1% FA). From the column fractions containing the product, the acetonitrile was removed at the rotary evaporator (bath temperature: 40 °C to avoid probe hydrolysis), then the aqueous solution was lyophilised overnight.

*Note*: If the probe cannot be afforded in the desired purity by this procedure, we recommend capping the purified fluorophores (see General Procedure B)

**General Procedure B: *O*-alkyl fluorescein fluorophores**

The corresponding probe (1.0 eq) was dissolved in methanol/THF (1:1, 0.008−0.02 M). Sodium hydroxide (aqueous, 2 M, 10 eq) was added and the reaction mixture was stirred at r.t. for 5 min. Hydrochloric acid (aqueous, 2 M, 12 eq) was added to acidify the solution and the volatiles were removed *in vacuo*. The crude product was purified by reversed-phase flash column chromatography (acetonitrile/water, 0.1% FA). The solvent of the product fractions was removed *in vacuo*.

<u>*Note*</u>: We found that the purification of the fluorophores was simplest to perform by preparing the probes in three steps only purifying the product and then cleaving the probe to the fluorophore again; however, purification of the fluorophore before capping is also feasible.

##### General Procedure C: *O*-Aryl carbamate formation

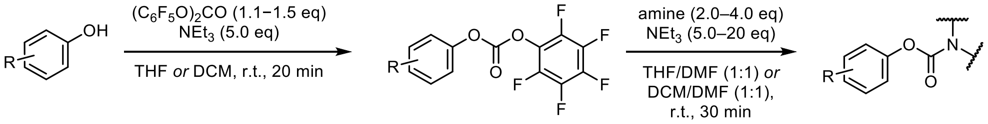

The corresponding phenol (1.0 eq) was dissolved in anhydrous THF or DCM (0.01−0.03 M) under nitrogen atmosphere, triethylamine (5.0 eq) and bis(pentafluorophenyl)carbonate (1.1−1.5 eq) were added and the reaction mixture was stirred at room temperature for 20 min. The desired amine (2.0−4.0 eq) was added as solid to the mixture, then anhydrous DMF was added. The reaction mixture was stirred at room temperature for 30 min. For isolation, the volatiles were removed *in vacuo* and the crude product was purified by reversed-phase flash column chromatography (acetonitrile/water, 0.1% formic acid).

##### General Procedure D: Buchwald-Hartwig cross coupling

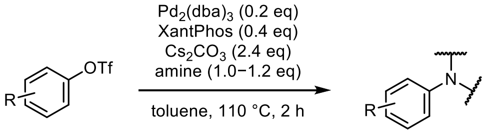

The corresponding triflate (1.0 eq) was placed in a round-bottom flask (dried in the dry-oven at 80 °C over night) and dissolved in anhydrous toluene (0.05 M) under nitrogen atmosphere. The respective amine (1.0−1.2 eq) and cesium carbonate (2.4 eq) were added and the solution was degassed by bubbling nitrogen through the solution for at least 5 min. XantPhos (20 mol%) and $Pd_2(dba)_3$ (10 mol%) were added and the reaction was heated to reflux for 2 h. The crude product was obtained by removing the volatiles *in vacuo*.

##### General Procedure E: Triflate cleavage with lithium hydroxide or TBAF

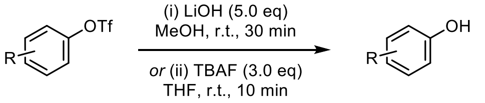

(i) The corresponding triflate was dissolved in methanol (0.07 M) and lithium hydroxide (5.0 eq) was added. The reaction mixture was stirred at room temperature for 20 min. Then aqueous HCl (2 M, 7.0 eq) was added before removing the volatiles *in vacuo*.

(ii) The corresponding triflate was dissolved in anhydrous THF (0.05−0.08 M) and TBAF (1 M solution in THF, 3.0 eq) was added. The reaction mixture was stirred at room temperature for 20 min.

##### General Procedure F: *tert*-Butyl carboxylate deprotection

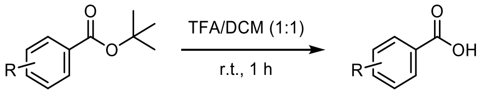

The corresponding *tert*-butyl carboxylate was dissolved in anhydrous DCM (0.01−0.02 M), trifluoroacetic acid (final solution: DCM:TFA = 1:1) was added and the reaction mixture was stirred at room temperature for 1 h. The crude product was afforded by removing the volatiles *in vacuo*.

##### General Procedure G: Acetoxymethylation

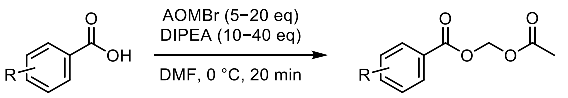

The corresponding carboxylic acid was dissolved in anhydrous DMF (0.01−0.02 M) at 0 °C, DIPEA (10−40 eq) and acetoxymethyl bromide (AOMBr, 5−20 eq) were added and the mixture was stirred at 0 °C for 20 min. The product was purified by preparative HPLC.

*Note: methyl piperazine probes also form the acetoxymethyl ammonium byproduct if reacted for longer or at room temperature.*

##### 7.2.3 Proof-of-concept fluorescein probes and fluorophores

**iC4-FS**

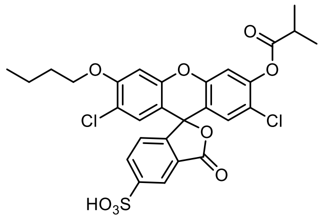

Prepared according to General Procedure A from **$H_2$-FS** (30 mg, 63 µmol, 1.0 eq; 0.10 M) and *n*-butyl iodide (43 µL, 0.38 mmol, 6.0 eq). Reaction time for alkylation: 6 h. Ester cleavage: 0.033 M in THF/water = 1:1 for 2 h, semi-purified by rp-column chromatography: 10→50% MeCN. Acylation: 0.01 M, purification by rp-column chromatography: 15→70% MeCN.

The product **iC4-FS** (18 mg, 30 µmol, 47% (3 steps)) was obtained as light yellow solid.

**TLC** *Rf* = 0.79 (*rp*, 60% MeCN)
**$^1$H-NMR** (400 MHz, DMSO-*d*$_6$): *δ* (ppm) = 8.10 (s, 1H), 8.00 (dd, J = 8.0, 1.2 Hz, 1H), 7.51 (s, 1H), 7.35 (d, J = 8.0 Hz, 1H), 7.18 (s, 1H), 7.12 (d, J = 2.4 Hz, 1H), 6.88 (s, 1H), 4.23 – 4.12 (m, 2H), 2.91 (hept, J = 6.9 Hz, 1H), 1.75 (p, J = 6.5 Hz, 2H), 1.46 (h, J = 7.3 Hz, 2H), 1.27 (d, J = 7.0 Hz, 6H), 0.94 (t, J = 7.4 Hz, 3H).
**$^{13}$C-NMR** (101 MHz, DMSO-*d*$_6$): *δ* (ppm) = 174.2, 168.3, 156.3, 151.9, 151.2, 150.4, 150.0, 148.6, 133.8, 129.4, 128.6, 125.9, 124.2, 122.3, 122.1, 118.2, 118.2, 113.5, 111.0, 102.3, 80.8, 69.5, 33.8, 30.8, 19.1, 19.1, 14.1.
**HRMS** (ESI+): m/z calc. for $C_{28}H_{25}Cl_2O_9S^+$ $[M+H]^+$: 607.0591, found: 607.0590.

**iC7-FS**

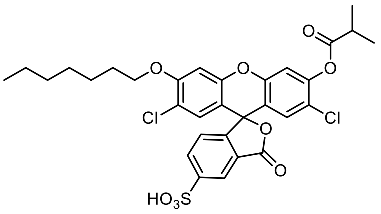

Prepared according to General Procedure A from **$H_2$-FS** (27 mg, 56 µmol, 1.0 eq; 0.04 M) and *n*-heptyl iodide (55 µL, 0.34 mmol, 6.0 eq). Reaction time for alkylation: 3 h. Ester cleavage: 0.02 M in THF/water = 2:1 for 3.5 h, semi-purified by rp-column chromatography: 15→70% MeCN. Acylation: 0.02 M, purification by rp-column chromatography: 30→100% MeCN.

The product **iC7-FS** (25 mg, 39 µmol, 74% (3 steps)) was obtained as light yellow solid.

**TLC** *Rf* = 0.67 (*rp*, 60% MeCN)
**$^1$H-NMR** (800 MHz, DMSO-*d*$_6$): *δ* (ppm) = 8.11 (s, 1H), 8.02 (d, J = 8.0 Hz, 1H), 7.51 (s, 1H), 7.35 (d, J = 7.9 Hz, 1H), 7.18 (s, 1H), 7.12 (s, 1H), 6.88 (s, 1H), 4.22 – 4.14 (m, 2H), 2.92 (hept, J = 6.8 Hz, 1H), 1.77 (p, J = 6.6 Hz, 2H), 1.44 (p, J = 7.8, 7.4 Hz, 2H), 1.35 (p, J = 7.3 Hz, 2H), 1.28 (d, J = 7.0 Hz, 6H), 1.32 – 1.21 (m, 4H), 0.87 (t, J = 6.8 Hz, 3H).
**$^{13}$C-NMR** (201 MHz, DMSO-*d*$_6$): *δ* (ppm) = 174.2, 168.2, 156.3, 151.9, 151.3, 150.4, 150.0, 148.6, 133.8, 129.3, 128.6, 125.9, 124.2, 122.3, 122.1, 118.2, 118.2, 113.5, 111.1, 102.3, 80.8, 69.8, 33.8, 31.7, 28.8, 28.7, 25.8, 22.5, 19.1, 14.4.
**HRMS** (ESI−): m/z calc. for $C_{31}H_{29}Cl_2O_9S^-$ $[M-H]^-$: 647.0915, found: 647.0917.

**iC10-FS**

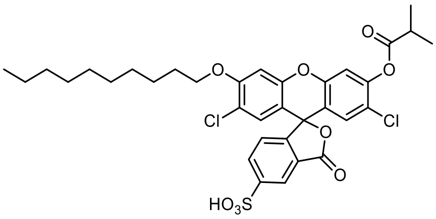

Prepared according to General Procedure A from **$H_2$-FS** (50 mg, 0.10 mmol, 1.0 eq; 0.02 M) and *n*-decyl iodide (0.13 mL, 0.62 mmol, 6.0 eq). Reaction time for alkylation: 2 h. Ester cleavage: 0.004 M in THF/water = 3:2 for 4 h, no purification. Acylation: 0.03 M, purification by rp-column chromatography: 40→100% MeCN.

The product **iC10-FS** (32 mg, 46 µmol, 45% (3 steps)) was obtained as colourless solid.

**TLC** *Rf* = 0.50 (*rp*, 60% MeCN)
**$^1$H-NMR** (800 MHz, DMSO-*d*$_6$): *δ* (ppm) = 8.10 (s, 1H), 8.01 (dd, J = 7.9, 1.2 Hz, 1H), 7.50 (s, 1H), 7.34 (d, J = 7.9 Hz, 1H), 7.17 (s, 1H), 7.11 (s, 1H), 6.87 (s, 1H), 4.21 – 4.13 (m, 2H), 2.91 (hept, J = 7.0 Hz, 1H), 1.76 (p, J = 6.6 Hz, 2H), 1.43 (p, J = 7.6, 7.2 Hz, 2H), 1.33 (p, J = 6.9 Hz, 2H), 1.27 (d, J = 7.0 Hz, 6H), 1.30 – 1.20 (m, 10H), 0.84 (t, J = 7.0 Hz, 3H).
**$^{13}$C-NMR** (201 MHz, DMSO-*d*$_6$): *δ* (ppm) = 174.2, 168.2, 156.3, 151.9, 151.3, 150.4, 150.0, 148.6, 133.8, 129.3, 128.6, 125.9, 124.2, 122.3, 122.1, 118.2, 118.2, 113.5, 111.1, 102.3, 80.8, 69.8, 33.8, 31.8, 29.4, 29.4, 29.1, 29.0, 28.7, 25.8, 22.6, 19.1, 14.4.
**HRMS** (ESI−): m/z calc. for $C_{34}H_{35}Cl_2O_9S^-$ [M−H]$^-$: 689.1384, found: 689.1387.

**iC4-FC**

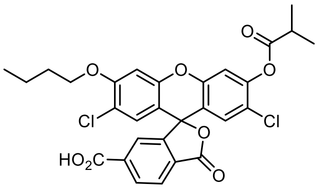

Prepared according to General Procedure A from **$H_2$-FC** (29 mg, 65 µmol, 1.0 eq; 0.07 M) and *n*-butyl iodide (60 µL, 0.52 mmol, 8.0 eq). Reaction time for alkylation: 3 h. Ester cleavage: 0.01 M in THF/water = 3:2 for 13 h, semi-purified by rp-column chromatography: 25→80% MeCN. Acylation: 0.01 M, purification by rp-column chromatography: 40→100% MeCN.

The product **iC4-FC** (10 mg, 17 µmol, 26% (3 steps)) was obtained as colourless solid.

**TLC** *Rf* = 0.41 (*rp*, 60% MeCN)
**$^1$H-NMR** (800 MHz, DMSO-*d*$_6$): *δ* (ppm) = 8.25 (d, J = 7.9 Hz, 1H), 8.11 (d, J = 7.6 Hz, 1H), 7.82 (s, 1H), 7.51 (s, 1H), 7.19 (s, 1H), 7.14 (s, 1H), 6.90 (s, 1H), 4.22 – 4.14 (m, 2H), 2.92 (hept, J = 7.0 Hz, 1H), 1.75 (p, J = 6.4 Hz, 2H), 1.47 (dt, J = 12.4, 6.2 Hz, 2H), 1.28 (d, J = 7.0 Hz, 6H), 0.95 (t, J = 7.4 Hz, 3H).
**$^{13}$C-NMR** (201 MHz, DMSO-*d*$_6$): *δ* (ppm) = 174.2, 168.0, 166.7, 156.3, 152.0, 150.9, 150.6, 150.1, 148.6, 131.8, 129.3, 128.8, 128.6, 125.8, 124.8, 122.1, 118.2, 118.1, 113.5, 110.9, 102.4, 81.1, 69.5, 33.8, 30.8, 19.1, 19.1, 14.1.
**HRMS** (ESI−): m/z calc. for $C_{29}H_{23}Cl_2O_8^-$ [M−H]$^-$: 569.0775, found: 569.0778.

**iC4-F**

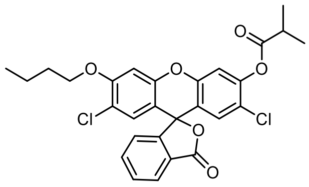

iC4-F

Prepared according to General Procedure A from **$H_2$-F** (82 mg, 0.20 mmol, 1.0 eq; 0.04 M) and *n*-butyl iodide (0.14 mL, 1.2 mmol, 6.0 eq). Reaction time for alkylation: 3 h. Ester cleavage: 0.01 M in THF/water = 3:2 for 13 h, semi-purified by rp-column chromatography: 10→50% MeCN. Acylation: 0.01 M, purification by np-column chromatography: Hex/EtOAc 0→20%.

The product **iC4-F** (41 mg, 78 µmol, 38% (3 steps)) was obtained as colourless solid.

**TLC** *Rf* = 0.26 (*np*, Hex:EtOAc 9:1)
**$^1$H-NMR** (400 MHz, $CD_2Cl_2$): $\delta$ (ppm) = 8.03 (dd, J = 7.3, 1.1 Hz, 1H), 7.75 (td, J = 7.5, 1.3 Hz, 1H), 7.70 (td, J = 7.5, 1.1 Hz, 1H), 7.24 – 7.18 (m, 1H), 7.14 (s, 1H), 6.87 (s, 1H), 6.86 (s, 1H), 6.77 (s, 1H), 4.12 – 4.05 (m, 2H), 2.88 (hept, J = 7.0 Hz, 1H), 1.90 – 1.79 (m, 2H), 1.53 (t, J = 8.7 Hz, 2H), 1.33 (dd, J = 7.0, 2.6 Hz, 6H), 0.99 (t, J = 7.4 Hz, 3H).
**$^{13}$C-NMR** (101 MHz, $CD_2Cl_2$): $\delta$ (ppm) = 174.5, 168.9, 156.8, 152.2, 150.9, 150.6, 148.9, 136.0, 130.8, 129.4, 128.9, 126.6, 125.7, 124.3, 122.6, 119.1, 118.3, 112.9, 111.1, 101.5, 81.5, 69.7, 34.5, 31.2, 19.5, 19.0, 19.0, 13.9.
**HRMS** (ESI+): m/z calc. for $C_{28}H_{24}Cl_2NaO_6^+$ $[M+Na]^+$: 549.0848, found: 549.0843.

**H-C4-FS**

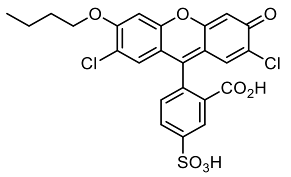

Prepared according to General Procedure B from **iC4-FS** (18 mg, 30 µmol, 1.0 eq, 0.008 M). Purification by rp column chromatography: 10→60% MeCN.

The product **H-C4-FS** (15 mg, 28 µmol, 94%) was obtained as orange solid.

**TLC** *Rf* = 0.71 (*rp*, 50% MeCN)
**$^1$H-NMR** (400 MHz, DMSO-*d*$_6$): $\delta$ (ppm) = 11.12 (s (br), 1H), 8.08 (s, 1H), 8.00 (d, J = 8.0 Hz, 1H), 7.28 (d, J = 7.9 Hz, 1H), 7.19 (s, 1H), 6.92 (s, 1H), 6.81 (s, 1H), 6.75 (s, 1H), 4.22 – 4.09 (m, 2H), 1.74 (p, J = 6.5 Hz, 2H), 1.45 (h, J = 7.4 Hz, 2H), 0.94 (t, J = 7.4 Hz, 3H)
**$^{13}$C-NMR** (101 MHz, DMSO-*d*$_6$): $\delta$ (ppm) = 168.4, 156.1, 155.7, 152.1, 151.1, 150.7, 150.4, 133.7, 128.9, 128.6, 126.1, 124.2, 122.0, 117.7, 116.9, 111.3, 110.6, 104.0, 102.4, 81.7, 69.4, 30.8, 19.1, 14.1.
**HRMS** (ESI−): m/z calc. for $C_{24}H_{17}Cl_2O_8S^-$ $[M-H]^-$: 535.0027, found: 535.0025.

**H-C7-FS**

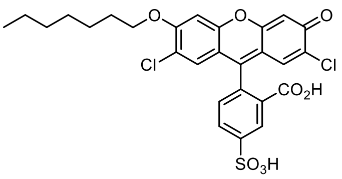

Prepared according to General Procedure B from **iC7-FS** (25 mg, 39 µmol, 1.0 eq, 0.01 M). Purification by rp column chromatography: 10→70% MeCN.

The product **H-C7-FS** (10 mg, 17 µmol, 45%) was obtained as orange solid.

**TLC** *Rf* = 0.66 (*rp*, 50% MeCN)
**$^{1}$H-NMR** (400 MHz, DMSO-*d*$_6$): *δ* (ppm) = 8.08 (s, 1H), 8.00 (dd, J = 8.0, 1.4 Hz, 1H), 7.27 (d, J = 7.9 Hz, 1H), 7.18 (s, 1H), 6.92 (s, 1H), 6.81 (s, 1H), 6.75 (s, 1H), 4.19 – 4.09 (m, 2H), 1.75 (p, J = 6.5 Hz, 2H), 1.48 – 1.38 (m, 2H), 1.38 – 1.30 (m, 2H), 1.30 – 1.23 (m, 4H), 0.90 – 0.81 (m, 3H).
**$^{13}$C-NMR** (101 MHz, DMSO-*d*$_6$): *δ* (ppm) = 168.0, 155.7, 155.3, 151.7, 150.6, 150.3, 149.9, 133.3, 128.5, 128.2, 125.7, 123.8, 121.6, 117.3, 116.5, 110.9, 110.2, 103.6, 102.0, 81.2, 69.3, 31.3, 28.4, 28.3, 25.4, 22.1, 14.0.
**HRMS** (ESI−): m/z calc. for $C_{27}H_{23}Cl_2O_8S^-$ [M−H]$^-$: 577.0496, found: 577.0496.

**H-C10-FS**

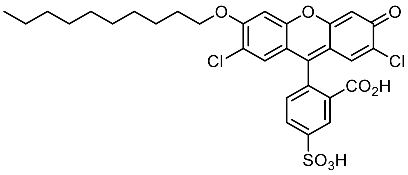

Prepared according to General Procedure B from **iC10-FS** (32 mg, 46 µmol, 1.0 eq, 0.008 M). Purification by rp column chromatography: 40→100% MeCN.

The product **H-C10-FS** (12 mg, 19 µmol, 41%) was obtained as orange solid.

**TLC** *Rf* = 0.50 (*rp*, 50% MeCN)
**$^{1}$H-NMR** (400 MHz, DMSO-*d*$_6$): *δ* (ppm) = 8.12 – 8.05 (m, 1H), 8.00 (dd, J = 8.0, 1.5 Hz, 1H), 7.26 (d, J = 7.9 Hz, 1H), 7.18 (s, 1H), 6.91 (s, 1H), 6.80 (s, 1H), 6.73 (s, 1H), 4.19 – 4.09 (m, 2H), 3.23 – 3.15 (m, 2H), 1.75 (p, J = 6.5 Hz, 2H), 1.67 – 1.55 (m, 2H), 1.48 – 1.38 (m, 2H), 1.24 (s, 6H), 0.89 – 0.80 (m, 5H).
**$^{13}$C-NMR** (101 MHz, DMSO-*d*$_6$): *δ* (ppm) = 168.0, 155.7, 150.7, 150.4, 150.0, 133.2, 128.4, 128.1, 125.7, 123.8, 121.7, 117.3, 117.0, 111.0, 109.8, 103.6, 101.9, 82.7, 69.3, 62.8, 31.3, 29.0, 28.7, 25.8, 25.4, 22.2, 21.7, 14.0.
**HRMS** (ESI−): m/z calc. for $C_{30}H_{29}Cl_2O_8S^-$ [M−H]$^-$: , found: 619.0963.

**H-C4-FC**

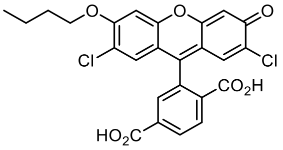

Was obtained purely during the preparation of iC4-FC according to General Procedure A from **H$_2$-FC** (0.30 g, 0.67 mmol, 1.0 eq, 0.07 M). Purification by rp column chromatography: 25→80% MeCN.

The product **H-C4-FC** (0.10 g, 0.21 mmol, 31%) was obtained as orange solid.

**TLC** *Rf* = 0.61 (*rp*, 60% MeCN)
**$^{1}$H-NMR** (400 MHz, DMSO-*d*$_6$): *δ* (ppm) = 11.14 (s, 1H), 8.24 (dd, J = 8.0, 1.2 Hz, 1H), 8.12 (d, J = 8.0 Hz, 1H), 7.75 (s, 1H), 7.20 (s, 1H), 6.93 (s, 1H), 6.87 (s, 1H), 6.80 (s, 1H), 4.15 (tt, J = 9.9, 5.0 Hz, 2H), 1.74 (p, J = 6.5 Hz, 2H), 1.46 (h, J = 7.4 Hz, 2H), 0.94 (t, J = 7.4 Hz, 3H).
**$^{13}$C-NMR** (101 MHz, DMSO-*d*$_6$): *δ* (ppm) = 168.0, 166.5, 156.1, 155.7, 152.3, 150.8, 150.4, 138.0, 131.7, 129.7, 128.9, 128.6, 126.3, 124.9, 117.7, 116.9, 111.1, 110.3, 104.0, 102.4, 82.1, 69.4, 30.8, 19.1, 14.1.
**HRMS** (ESI−): m/z calc. for $C_{25}H_{17}Cl_2O_7^-$ [M−H]$^-$: 499.0357, found: 499.0361.

**H-C4-F**

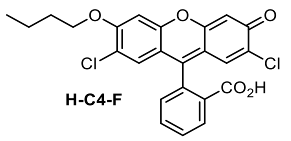

Prepared according to General Procedure B from **iC4-F** (30 mg, 57 µmol, 1.0 eq, 0.01 M). Purification by rp column chromatography: 40→100% MeCN.

The product **H-C4-F** (22 mg, 48 µmol, 85%) was obtained as orange solid.

**TLC** *Rf* = 0.37 (*rp*, 60% MeCN)
**$^1$H-NMR** (400 MHz, DMSO-*d*$_6$): *δ* (ppm) = 8.02 (d, J = 7.5 Hz, 1H), 7.82 (t, J = 7.2 Hz, 1H), 7.75 (t, J = 7.4 Hz, 1H), 7.33 (d, J = 7.6 Hz, 1H), 7.20 (s, 1H), 6.92 (s, 1H), 6.74 (s, 1H), 6.68 (s, 1H), 4.22 – 4.08 (m, 2H), 1.74 (p, J = 6.5 Hz, 2H), 1.52 – 1.36 (m, 2H), 0.93 (t, J = 7.4 Hz, 3H).
**$^{13}$C-NMR** (101 MHz, DMSO-*d*$_6$): *δ* (ppm) = 168.3, 155.6, 155.4, 151.5, 150.4, 150.1, 136.0, 130.6, 128.3, 128.0, 125.9, 125.2, 124.0, 117.2, 116.5, 111.2, 110.3, 103.6, 102.0, 81.4, 69.0, 30.4, 18.7, 13.7.
**HRMS** (ESI−): m/z calc. for $C_{24}H_{17}Cl_2O_5^-$ [M−H]$^-$: 455.0459, found: 455.0461.

**GSH activatable fluorescein probe GL-C4-FC**

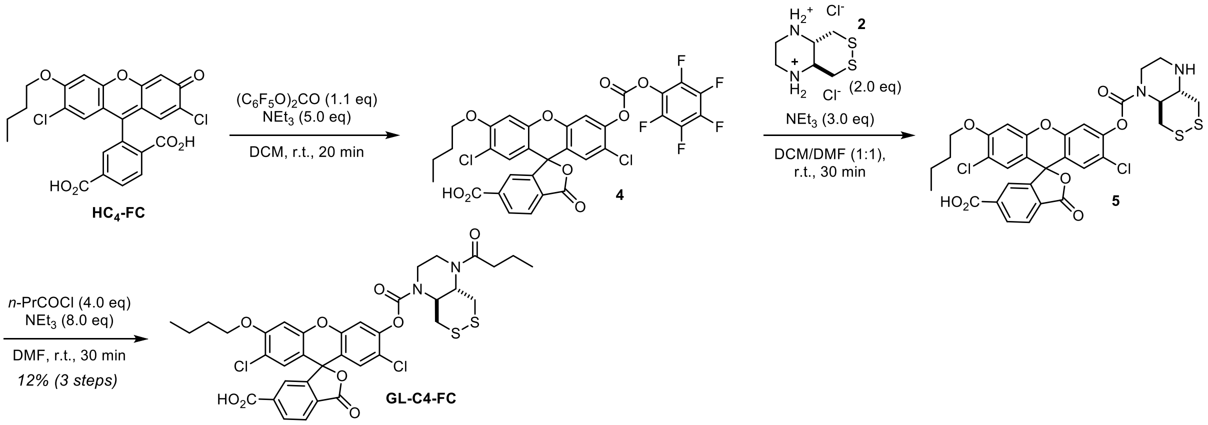

Compound **2** was prepared according to a known procedure (Zeisel, Felber *et al.*[36], compound 7T).

**H-C4-FC** (10 mg, 20 µmol, 1.0 eq, 0.007 M) was transformed to **5** following General Procedure C in DCM with 1.1 eq bis(pentafluorophenyl)carbonate and 2.0 eq of amine **2** with 5.0 eq triethylamine (purification: 15→100% MeCN). The semi-purified intermediate **5** was dissolved in anhydrous DMF (3 mL), triethylamine (22 µL, 0.16 mmol, 8.0 eq) and *n*-butyryl chloride (8.4 µL, 80 µmol, 4.0 eq) were added and the reaction mixture was stirred at room temperature for 30 min. The volatiles were removed *in vacuo* and the crude product was purified by preparative HPLC (acetonitrile/water, 0.1% formic acid; 50→100% MeCN, 20 min) affording **GL-C4-FC** (1.8 mg, 2.3 µmol, 12% over 3 steps) as colourless solid.

_Note_: dichloromethane should be used as solvent for the transformation of **H-C4-FC** to **4** since the same reagents in THF resulted in significant activation of the 6-carboxylate to form a mix of the desired carbamate and the byproducts 6-carboxy amide and both carbamate and carboxy-amide (and unreacted starting material) in equal amounts with primary/secondary amines (reaction in DMF almost cleanly affords the carboxy-amide and no carbamate).

**TLC** *Rf* = 0.37 (*rp*, 60% MeCN)
**$^1$H-NMR** (800 MHz, DMSO-*d*$_6$): *δ* (ppm) = 8.25 (dd, J = 7.9, 2.5 Hz, 1H), 8.11 (d, J = 7.8 Hz, 1H), 7.80 (s, 1H), 7.44 (s, 1H), 7.20 (s, 1H), 7.13 (s, 1H), 6.91 (s, 1H), 4.29 (s, 1H), 4.22 – 4.15 (m, 2H), 4.14 – 4.00 (m, 1H), 4.03 – 3.88 (m, 1H), 3.88 – 3.72 (m, 2H), 3.72 – 3.35 (m, 4H), 3.25 – 2.94 (m, 1H), 2.38 – 2.21 (m, 2H), 1.75 (p, J = 6.6 Hz, 2H), 1.51 – 1.43 (m, 4H), 0.95 (t, J = 7.4 Hz, 3H), 0.86 (t, J = 6.4 Hz, 3H).
**$^{13}$C-NMR** (201 MHz, DMSO-*d*$_6$): *δ* (ppm) = 172.3, 168.0, 166.5, 162.8, 156.3, 152.1, 152.1, 150.6, 150.1/150.0, 148.9, 131.9, 129.2, 129.2, 128.6, 126.3, 124.8, 122.3, 118.2, 117.8, 113.5, 110.9, 102.4, 81.1, 69.5, 58.4, 57.9, 46.8, 38.9, 38.4, 37.1, 35.2, 30.8, 19.1, 18.5, 14.2, 14.1.
**HRMS** (ESI−): m/z calc. for $C_{36}H_{33}Cl_2N_2O_9S_2^-$ [M−H]$^-$: 771.1010, found: 771.1022.

##### 7.2.4 Proof-of-concept rhodol probes and fluorophores

###### Synthetic Route for GL-Rho

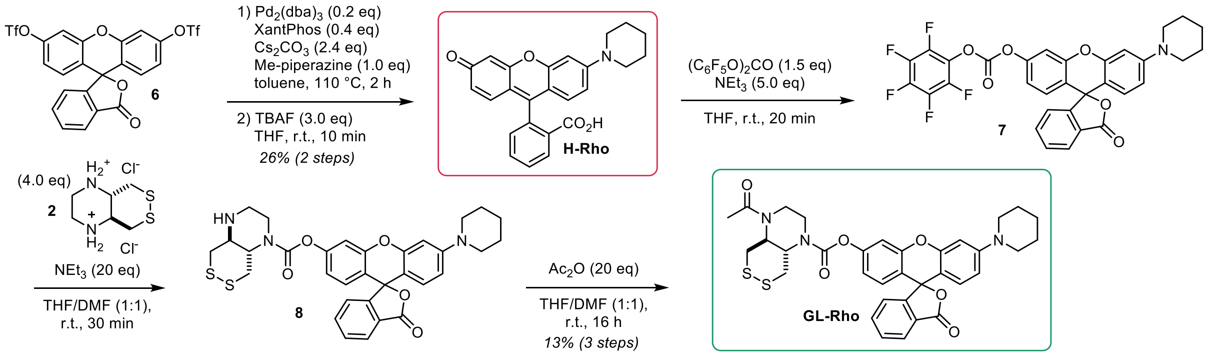

Fluorescein ditriflate **6** (He *et al.*[37], compound 3) and compound **2** (Zeisel, Felber *et al.*[36], compound 7T) were prepared according known procedures.

###### H-Rho

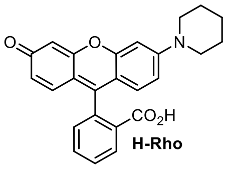

Buchwald-Hartwig coupling according to General Procedure D from fluorescein ditriflate **6** (0.15 g, 0.25 mmol, 1.0 eq, 0.05 M) with piperidine (25 µL, 0.25 mmol, 1.0 eq) followed by triflate cleavage according to General Procedure E(ii) (0.08 M). Purification by rp column chromatography: 10→45% MeCN.

The product **H-Rho** (26 mg, 66 µmol, 26% over 2 steps) was obtained as red solid.

**$^1$H-NMR** (400 MHz, MeOD-$d_4$): $\delta$ (ppm) = 8.07 (d, $J$ = 7.3 Hz, 1H), 7.77 (td, $J$ = 7.5, 1.3 Hz, 1H), 7.71 (td, $J$ = 7.5, 1.1 Hz, 1H), 7.23 (d, $J$ = 7.4 Hz, 1H), 6.86 – 6.78 (m, 2H), 6.75 – 6.68 (m, 3H), 6.60 (dd, $J$ = 8.8, 2.3 Hz, 1H), 3.40 – 3.35 (m, 4H), 1.75 – 1.64 (m, 6H).
**HRMS** (ESI+): m/z calc. for $C_{25}H_{22}NO_4^+$ [M+H]$^+$: 400.1543, found: 400.1531.

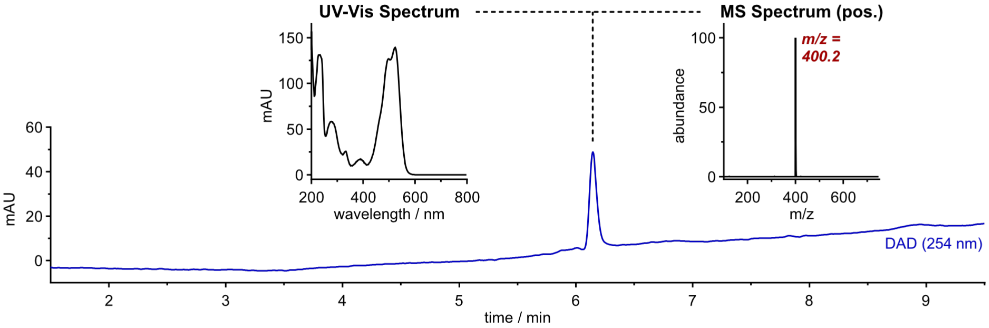

#### GL-Rho

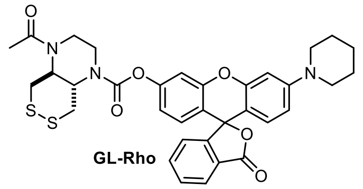

**H-Rho** (24 mg, 60 µmol, 1.0 eq, 0.03 M) was transformed into **8** following General Procedure C in THF with 1.5 eq bis(pentafluorophenyl)carbonate and 4.0 eq of amine **2** with 20 eq triethylamine. Then, acetic anhydride (0.11 mL, 1.2 mmol, 20 eq) was added to the reaction mixture of the intermediate product and stirred at room temperature for 16 h. The volatiles were removed *in vacuo* and the crude product was purified by preparative HPLC (acetonitrile/water, 0.1% formic acid; 20→100% MeCN, 20 min) affording **GL-Rho** (3.5 mg, 5.5 µmol, 13% over 3 steps) as light-red solid.

**$^1$H-NMR** (400 MHz, DMSO-$d_6$): $\delta$ (ppm) = 8.03 (d, J = 7.5 Hz, 1H), 7.81 (td, J = 7.5, 1.2 Hz, 1H), 7.74 (td, J = 7.5, 0.9 Hz, 1H), 7.31 (d, J = 7.6 Hz, 1H), 7.16 (s, 1H), 6.85 (t, J = 8.7 Hz, 1H), 6.80 (d, J = 8.7 Hz, 1H), 6.79 – 6.72 (m, 3H), 6.55 (d, J = 8.7 Hz, 1H), 4.27 (t, J = 10.6 Hz, 1H), 3.96 (s, 2H), 3.69 (dq, J = 35.9, 18.0 Hz, 5H), 3.26 (s, 4H), 3.05 (d, J = 11.7 Hz, 1H), 2.06 (s, 3H), 1.58 (s, 6H).
**$^{13}$C-NMR** (101 MHz, DMSO-$d_6$): $\delta$ (ppm) = 170.7, 169.1, 153.4, 152.8, 151.8, 136.2, 130.8, 129.4, 128.9, 126.4, 125.2, 124.5, 118.3, 116.8, 112.7, 110.5, 107.2, 101.2, 82.8, 58.9, 58.2, 48.8, 25.3, 24.4, 22.4.
**HRMS** (ESI+): m/z calc. for $C_{34}H_{34}N_3O_6S_2^+$ [M+H]$^+$: 644.1884, found: 644.1881.

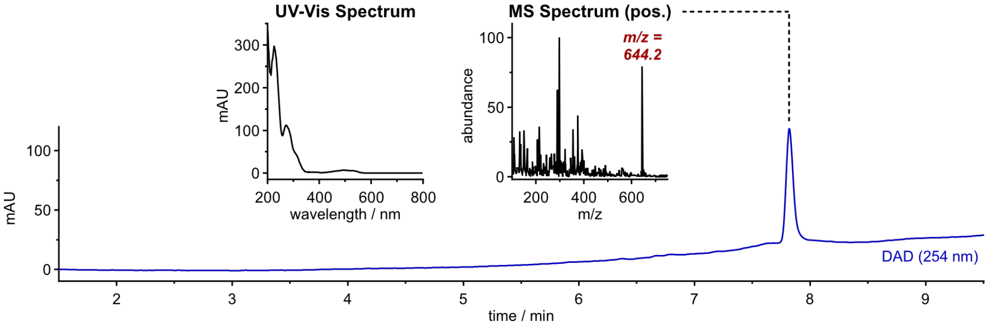

#### Synthetic Route for GL-Rho-A

**GL-Rho-A**

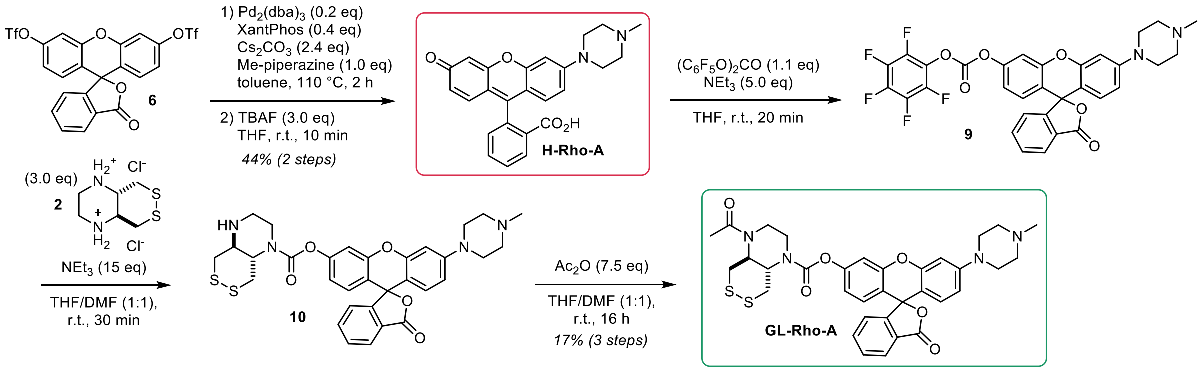

Fluorescein ditriflate (**6**) was prepared according to a known procedure (He *et al.*[37], compound 3).

#### H-Rho-A

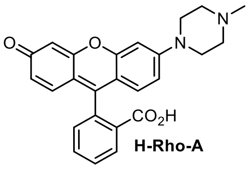

Buchwald-Hartwig coupling according to General Procedure D from fluorescein ditriflate **6** (0.15 g, 0.25 mmol, 1.0 eq, 0.05 M) with methyl piperazine (28 µL, 0.25 mmol, 1.0 eq) followed by triflate cleavage according to General Procedure E(ii) (0.08 M). Purification by rp column chromatography: 10→40% MeCN.

The product **H-Rho-A** (46 mg, 0.11 mmol, 44% over 2 steps) was obtained as red solid.

**$^1$H-NMR** (400 MHz, MeOD-$d_4$): $\delta$ (ppm) = 8.40 (s, 1H), 8.01 (d, J = 7.6 Hz, 1H), 7.77 (td, J = 7.4, 2.4 Hz, 1H), 7.70 (td, J = 7.3, 2.7 Hz, 1H), 7.19 (dd, J = 7.5, 4.1 Hz, 1H), 6.88 (dd, J = 24.0, 2.2 Hz, 1H), 6.77 (ddd, J = 11.7, 8.9, 2.3 Hz, 1H), 6.69 – 6.62 (m, 2H), 6.62 – 6.57 (m, 1H), 6.54 (dd, J = 8.7, 2.0 Hz, 1H), 3.67 – 3.58 (m, 3H), 3.44 (s, 2H), 3.26 (s, 2H), 3.14 – 3.05 (m, 2H), 2.72 (s, 2H).
**HRMS** (ESI+): m/z calc. for $C_{25}H_{23}N_2O_4^+$ [M+H]$^+$: 415.1652, found: 415.1641

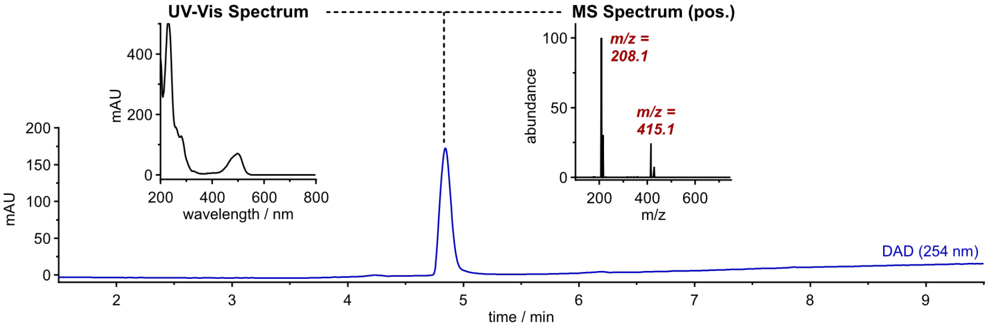

#### GL-Rho-A

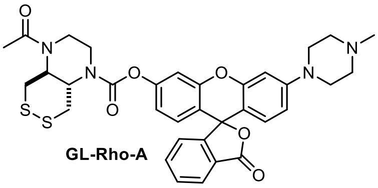

**H-Rho-A** (20 mg, 48 µmol, 1.0 eq, 0.02 M) was transformed into **10** following General Procedure C in THF with 1.1 eq bis(pentafluorophenyl)carbonate and 3.0 eq of amine **2** with 15 eq triethylamine. Then, acetic anhydride (34 µL, 0.36mmol, 7.5 eq) was added to the reaction mixture of the intermediate product and stirred at room temperature for 16 h. The volatiles were removed *in vacuo* and the crude product was purified by preparative HPLC (acetonitrile/water, 0.1% formic acid; 20→75% MeCN, 20 min) affording **GL-Rho-A** (5.3 mg, 8.1 µmol, 17% over 3 steps) as colourless solid.

**$^1$H-NMR** (400 MHz, DMSO-$d_6$): $\delta$ (ppm) = 8.10 (d, J = 7.5 Hz, 1H), 7.88 (t, J = 7.4 Hz, 1H), 7.81 (t, J = 7.4 Hz, 1H), 7.38 (d, J = 7.5 Hz, 1H), 7.23 (s, 1H), 6.94 (d, J = 8.4 Hz, 1H), 6.87 (dd, J = 12.2, 7.8 Hz, 3H), 6.65 (d, J = 8.8 Hz, 1H), 4.34 (t, J = 10.2 Hz, 1H), 4.04 (s, 2H), 3.87 – 3.61 (m, 4H), 3.60 – 3.45 (m, 2H), 3.12 (d, J = 12.3 Hz, 1H), 2.58 (s, 8H), 2.29 (s, 3H), 2.13 (s, 3H).
**$^{13}$C-NMR** (101 MHz, DMSO-$d_6$): $\delta$ (ppm) = 170.7, 169.1, 153.2, 152.8, 152.4, 152.0, 151.7, 136.2, 130.8, 129.4, 128.9, 126.3, 125.3, 124.5, 118.4, 116.8, 112.5, 110.6, 108.0, 101.3, 82.7, 59.9, 58.9, 54.8, 47.5, 46.2, 38.8, 37.3, 22.4.
**HRMS** (ESI+): m/z calc. for $C_{34}H_{35}N_4O_6S_2^+$ [M+H]$^+$: 659.1993, found: 659.1993

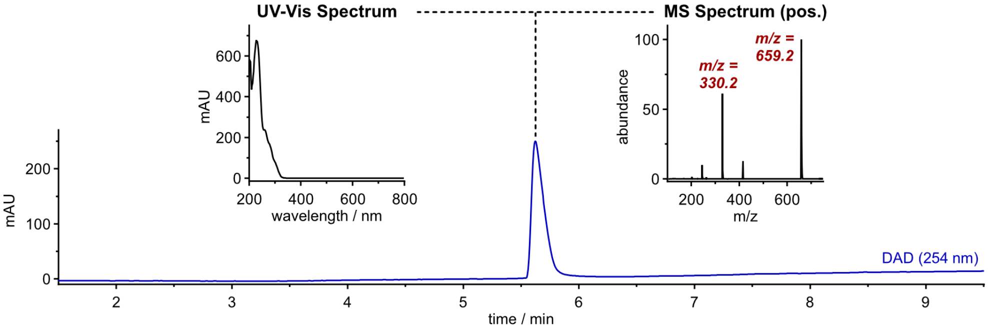

#### Synthetic Route for GL-Rho-C and GL-Rho-C$^{m}$

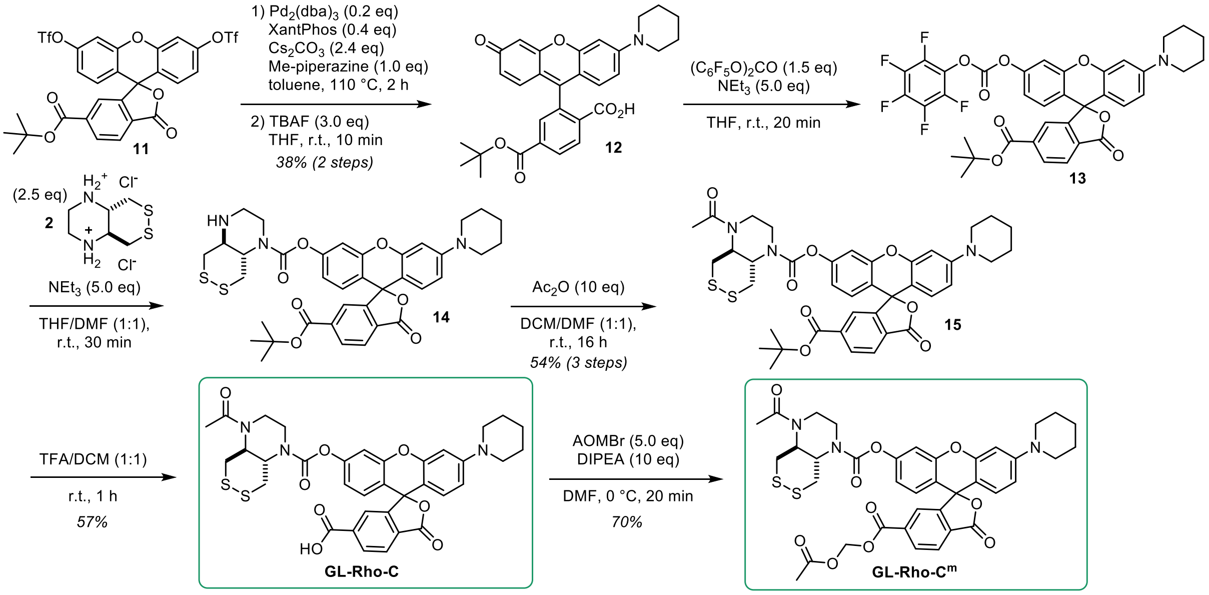

6-*tert*-Butoxycarbonylfluorescein (**11**) ditriflate was prepared according to a previously described procedure (Grimm *et al.*[38], compound S4).

##### Compound 12

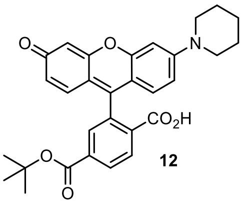

Buchwald-Hartwig coupling according to General Procedure D from ditriflate **11** (50 mg, 72 µmol, 1.0 eq, 0.04 M) with piperidine (7 µL, 72 µmol, 1.0 eq) followed by triflate cleavage according to General Procedure E(ii) (0.05 M). Semi-purification by rp column chromatography: 0→60% MeCN.

The product **12** (14 mg, 27 µmol, 38% over 2 steps) was obtained as red solid.

**$^{1}$H-NMR** (400 MHz, MeOD-*d*$_4$): $\delta$ (ppm) = 8.25 (dd, J = 8.1, 1.4 Hz, 1H), 8.12 (d, J = 8.1 Hz, 1H), 7.73 – 7.70 (m, 1H), 6.91 – 6.84 (m, 2H), 6.82 – 6.73 (m, 3H), 6.63 (dd, J = 8.9, 2.3 Hz, 1H), 3.44 (s, 4H), 1.69 (s, 6H), 1.55 (s, 9H).
**LRMS** (ESI+): m/z calc. for $C_{30}H_{30}NO_6^{+}$ [M+H]$^{+}$: 500.2, found: 500.2.

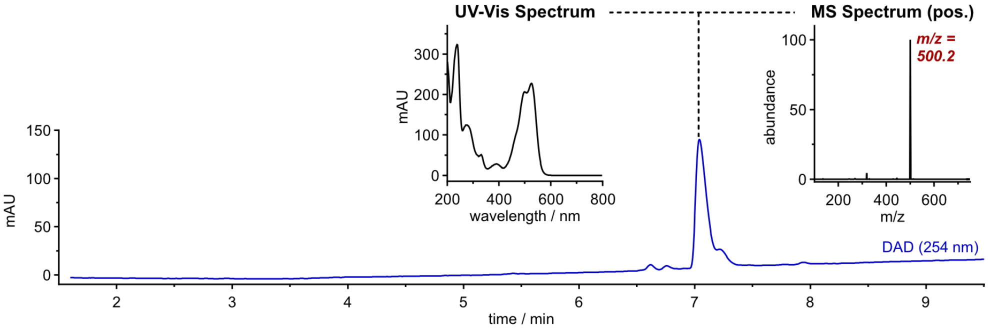

**Compound 15**

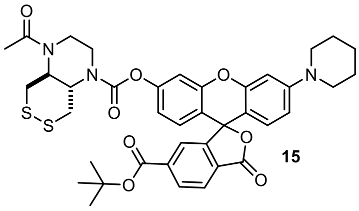

15

**12** (5 mg, 10 µmol, 1.0 eq, 0.005 M) was transformed into **14** following General Procedure C in THF with 1.5 eq bis(pentafluorophenyl)carbonate and 2.5 eq of amine **2** with 5.0 eq triethylamine. Then, acetic anhydride (9.4 µL, 0.10mmol, 10 eq) was added to the reaction mixture of the intermediate product and stirred at room temperature for 16 h. The volatiles were removed *in vacuo* and the crude product was purified by np flash column chromatography (DCM/MeOH, 0→3% MeOH) affording **15** (4 mg, 5.4 µmol, 54% over 3 steps) as colourless solid.

**TLC** *Rf* = 0.70 (*np*, DCM:MeOH 19:1)
**HRMS** (ESI+): m/z calc. for $C_{39}H_{42}N_3O_8S_2^+$ [M+H]$^+$: 744.2408, found: 744.2396.

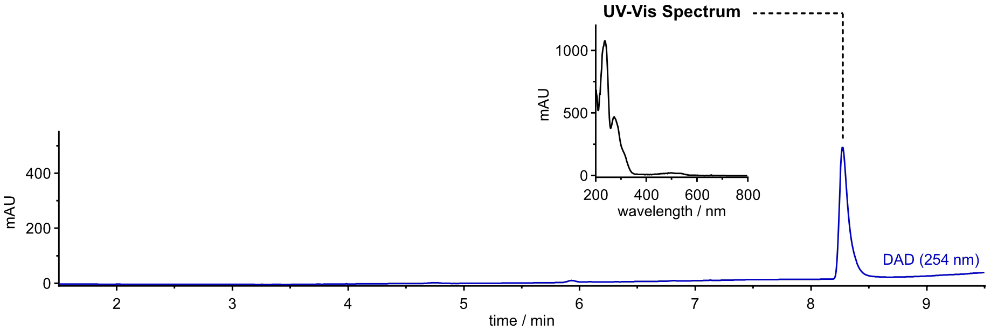

**GL-Rho-C**

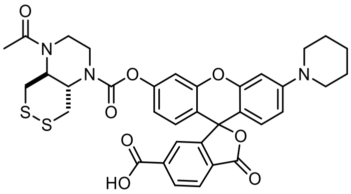

**GL-Rho-C**

Prepared according to General Procedure F from **15** (4 mg, 5.4 µmol, 1.0 eq, 0.01 M). The crude product was purified by preparative HPLC (acetonitrile/water, 0.1% formic acid; 15→100% MeCN, 20 min) affording **GL-Rho-C** (2.1 mg, 3.1 µmol, 57%) as colourless solid.

**$^1$H-NMR** (800 MHz, DMSO-$d_6$): $\delta$ (ppm) = 8.21 (d, J = 8.1 Hz, 1H), 8.08 (d, J = 8.0 Hz, 1H), 7.64 (s, 1H), 7.16 (s, 1H), 6.87 – 6.82 (m, 2H), 6.78 (d, J = 2.5 Hz, 1H), 6.74 (dd, J = 9.0, 2.5 Hz, 1H), 6.58 (d, J = 8.9 Hz, 1H), 4.33 – 4.13 (m, 1H), 4.08 – 3.79 (m, 2H), 3.76 – 3.54 (m, 4H), 3.28 – 3.22 (m, 4H), 3.05 (d, J = 12.1 Hz, 1H), 2.05 (s, 3H), 1.61 – 1.53 (m, 6H).
**$^{13}$C-NMR** (201 MHz, DMSO-$d_6$): $\delta$ (ppm) = 170.1, 168.1, 166.1, 153.0, 152.5, 152.0, 151.7, 151.3, 131.0, 129.0, 128.5, 125.0, 124.2, 117.9, 115.9, 112.2, 110.1, 106.4, 100.7, 82.5, 58.4, 57.8, 48.3, 40.0, 24.8, 23.9, 21.9.
**HRMS** (ESI+): m/z calc. for $C_{35}H_{34}N_3O_8S_2^+$ [M+H]$^+$: 688.1782, found: 688.1765.

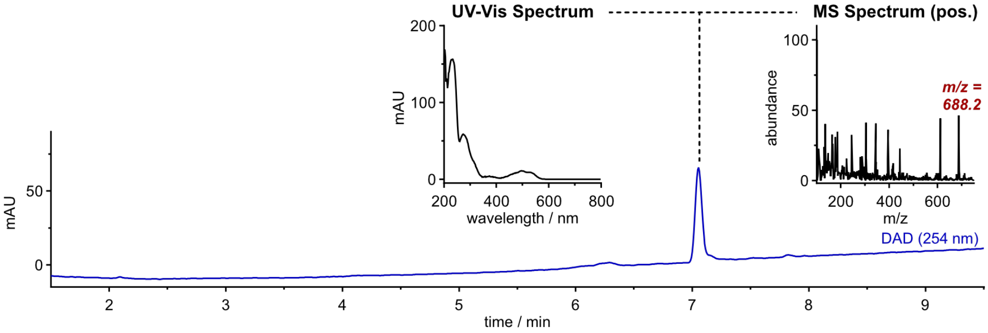

**GL-Rho-C$^{m}$**

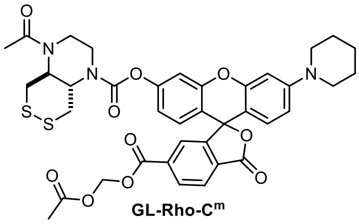

GL-Rho-C$^{m}$

Prepared according to General Procedure G from **GL-Rho-C** (3.7 mg, 5.4 µmol, 1.0 eq, 0.01 M) with 10 eq DIPEA and 5.0 eq acetoxymethyl bromide. The crude product was purified by preparative HPLC (acetonitrile/water, 0.1% formic acid; 20→100% MeCN, 20 min) affording **GL-Rho-C$^{m}$** (1.6 mg, 2.1 µmol, 70%) as colourless solid.

**$^{1}$H-NMR** (800 MHz, DMSO-$d_6$): $\delta$ (ppm) = 8.27 (d, J = 8.1 Hz, 1H), 8.19 (d, J = 8.1 Hz, 1H), 7.75 (s, 1H), 7.17 (s, 1H), 6.88 (d, J = 8.6 Hz, 1H), 6.85 (s, 1H), 6.78 (s, 1H), 6.74 (dd, J = 9.0, 1.9 Hz, 1H), 6.60 (d, J = 8.9 Hz, 1H), 5.87 (s, 2H), 4.31 – 4.14 (m, 1H), 3.95 (s, 2H), 3.78 – 3.53 (m, 4H), 3.53 – 3.37 (m, 2H), 3.27 (s, 4H), 3.04 (d, J = 11.9 Hz, 1H), 2.06 (s, 6H), 1.57 (s, 6H).
**$^{13}$C-NMR** (201 MHz, DMSO-$d_6$): $\delta$ (ppm) = 169.3, 167.6, 163.4, 153.0, 152.5, 152.2, 151.7, 151.4, 135.1, 131.3, 130.3, 129.2, 128.6, 125.8, 124.7, 117.9, 115.6, 112.2, 110.1, 105.9, 100.7, 82.9, 80.1, 58.5, 57.7, 48.3, 40.0, 24.9, 23.9, 21.9, 20.5.
**HRMS** (ESI+): m/z calc. for $C_{38}H_{38}N_3O_{10}S_2^+$ [M+H]$^+$: 760.1993, found: 760.1996.

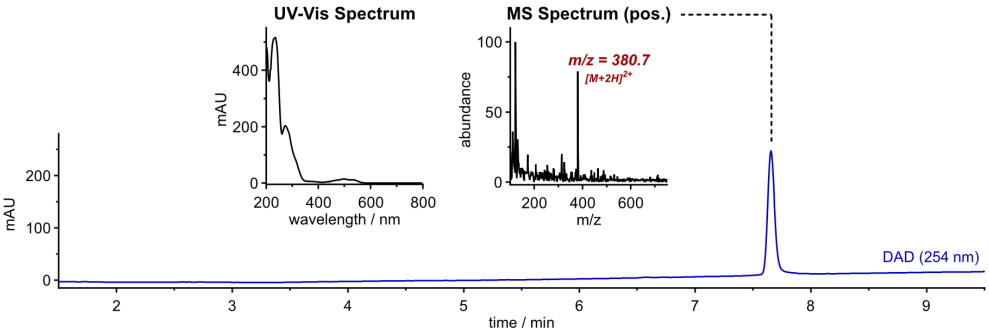

**H-Rho-C**

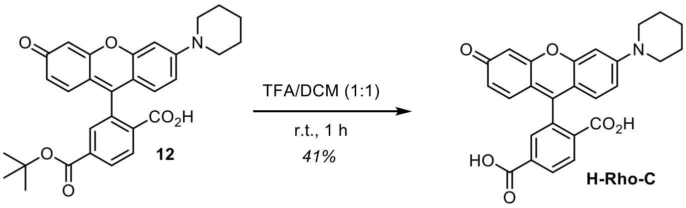

Prepared according to General Procedure F from **12** (6 mg, 12 µmol, 1.0 eq, 0.02 M). The crude product was purified by preparative HPLC (acetonitrile/water, 0.1% formic acid; 5→50% MeCN, 20 min) affording **H-Rho-C** (2.2 mg, 4.9 µmol, 41%) as red solid.

**$^{1}$H-NMR** (400 MHz, DMSO-$d_6$): $\delta$ (ppm) = 8.22 – 8.16 (m, 1H), 8.05 (d, J = 8.0 Hz, 1H), 7.60 (s, 1H), 6.76 (d, J = 2.4 Hz, 1H), 6.70 (dd, J = 9.0, 2.5 Hz, 1H), 6.68 (d, J = 2.3 Hz, 1H), 6.59 (d, J = 8.7 Hz, 1H), 6.57 – 6.49 (m, 2H), 3.25 (s, 4H), 1.56 (s, 6H).
**$^{13}$C-NMR** (101 MHz, DMSO-$d_6$): $\delta$ (ppm) = 168.2, 166.3, 159.6, 152.8, 152.1, 151.9, 140.3, 130.8, 129.2, 128.4, 124.9, 124.3, 112.6, 111.9, 109.3, 106.9, 102.2, 100.8, 83.7, 48.4, 24.8, 23.9.
**HRMS** (ESI+): m/z calc. for $C_{26}H_{22}NO_6^+$ [M+H]$^+$: 444.1442, found: 444.1435.

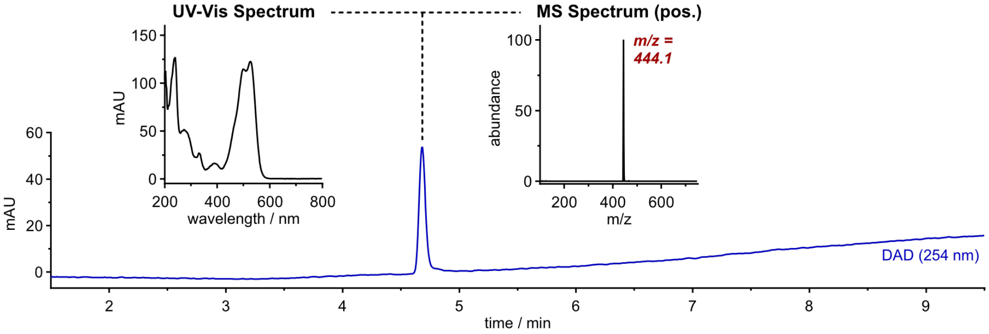

#### Synthetic Route for GL-Rho-AC and GL-Rho-AC^m (= GL-TraG)

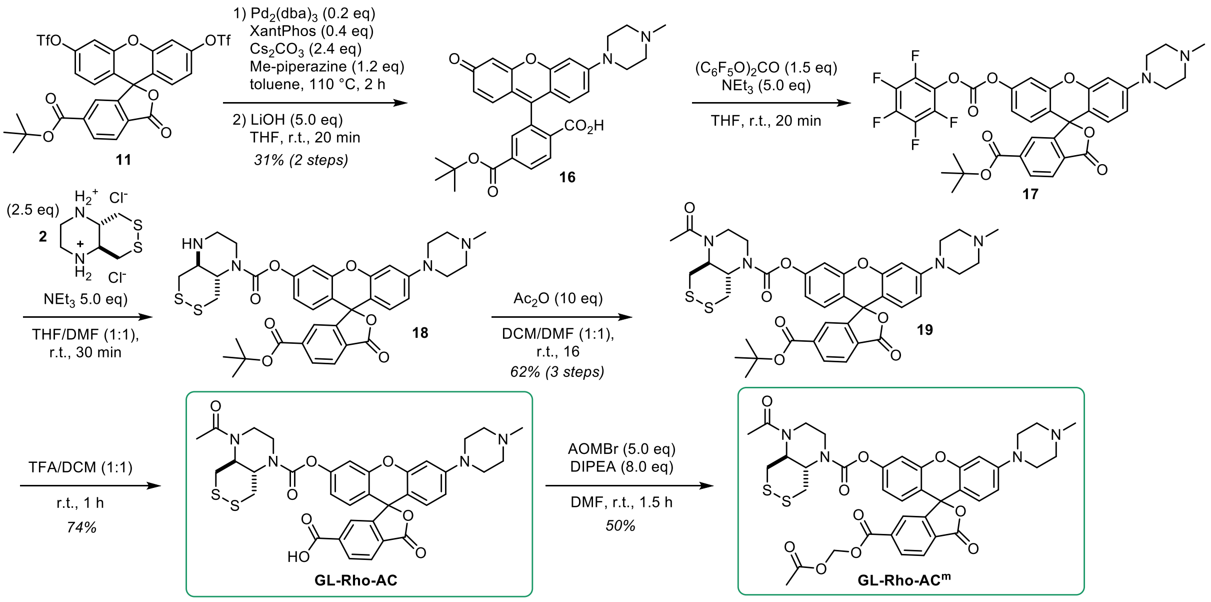

#### Compound 16

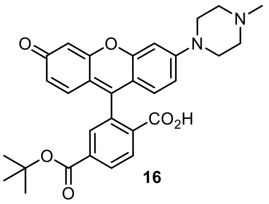

Buchwald-Hartwig coupling according to General Procedure D from triflate **11** (70 mg, 0.10 mmol, 1.0 eq, 0.05 M) with methyl piperazine (13 µL, 0.12 mmol, 1.2 eq) followed by triflate cleavage according to General Procedure E(i) (0.07 M). Purification by rp column chromatography: 2→35% MeCN.

The product **16** (16 mg, 31 µmol, 31% over 2 steps) was obtained as red solid.

**TLC** *Rf* = 0.43 (*np*, DCM:MeOH 4:1)
**$^1$H-NMR** (400 MHz, MeOD-*d*$_4$): *δ* (ppm) = 8.25 (dd, J = 8.0, 1.2 Hz, 1H), 8.08 (d, J = 8.0 Hz, 1H), 7.67 (s, 1H), 6.84 (d, J = 1.8 Hz, 1H), 6.75 (dd, J = 9.0, 1.9 Hz, 1H), 6.71 – 6.61 (m, 3H), 6.55 (dd, J = 8.8, 2.2 Hz, 1H), 3.47 (s, 4H), 3.28 (s, 4H), 2.86 (s, 3H), 1.52 (s, 9H).
**HRMS** (ESI+): m/z calc. for $C_{30}H_{31}N_2O_6^+$ [M+H]$^+$: 515.2177, found: 515.2169.

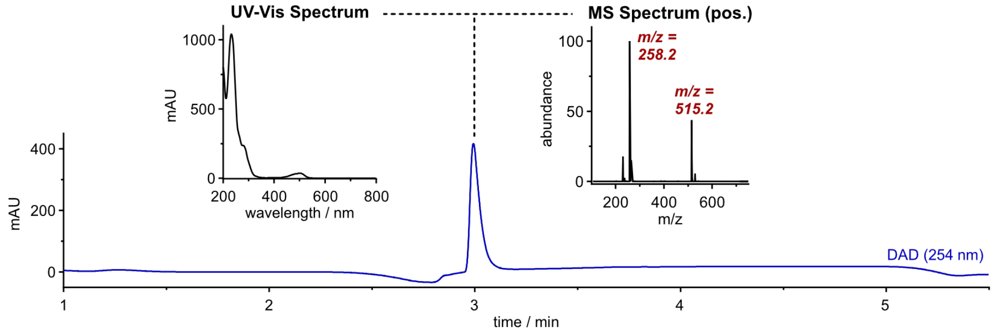

##### Compound 19

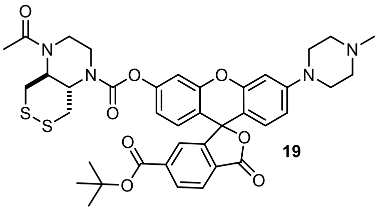

19

**16** (30 mg, 51 µmol, 1.0 eq, 0.03 M) was transformed into **19** following General Procedure C in THF with 1.5 eq bis(pentafluorophenyl)carbonate and 2.5 eq of amine **2** with 5 eq triethylamine (purification: 15→100% MeCN). Then, acetic anhydride (16 µL, 0.17mmol, 10 eq) was added to the reaction mixture of the intermediate product and stirred at room temperature for 16 h. The volatiles were removed *in vacuo* and the crude product was was purified by np flash column chromatography (DCM/MeOH, 0→8% MeOH) affording **19** (8 mg, 11 µmol, 62% over 3 steps) as colourless solid.

**TLC** *Rf* = 0.40 (*np*, DCM/MeOH 19:1)
**HRMS** (ESI+): m/z calc. for $C_{39}H_{43}N_4O_8S_2^+$ [M+H]$^+$: 759.2517, found: 759.2499

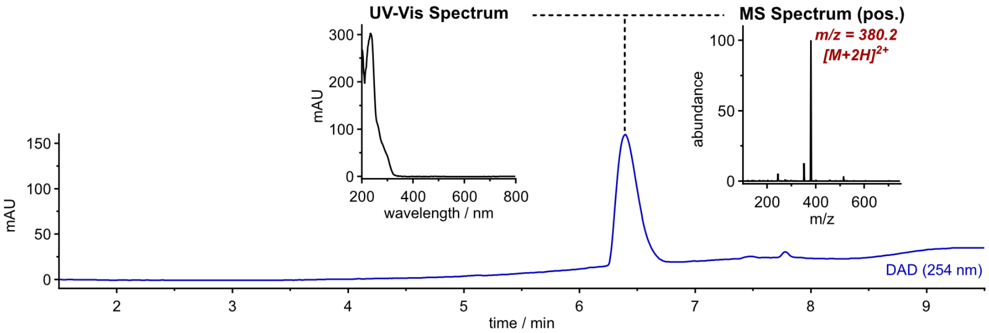

##### GL-Rho-AC

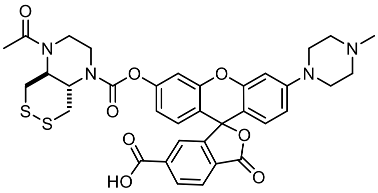

GL-Rho-AC

Prepared according to General Procedure F from **19** (8 mg, 11 µmol, 1.0 eq, 0.02 M). The crude product was purified by preparative HPLC (acetonitrile/water, 0.1% formic acid; 5→40% MeCN, 20 min) affording **GL-Rho-AC** (5.5 mg, 8.0 µmol, 74%) as colourless solid.

**$^1$H-NMR** (800 MHz, DMSO-$d_6$): $\delta$ (ppm) = 8.19 (d, J = 8.0 Hz, 1H), 8.05 (d, J = 7.4 Hz, 1H), 7.61 (s, 1H), 7.16 (s, 1H), 6.85 (s, 2H), 6.82 (s, 1H), 6.76 (d, J = 8.6 Hz, 1H), 6.61 (d, J = 8.8 Hz, 1H), 4.34 – 4.13 (m, 1H), 4.08 – 3.79 (m, 2H), 3.77 – 3.55 (m, 4H), 3.53 – 3.39 (m, 2H), 3.26 (s, 4H), 3.05 (d, J = 12.1 Hz, 1H), 2.59 – 2.53 (m, 4H), 2.30 (s, 3H), 2.05 (s, 3H).
**$^{13}$C-NMR** (201 MHz, DMSO-$d_6$): $\delta$ (ppm) = 170.1, 168.2, 166.5, 163.3, 152.6, 152.6, 152.0, 151.5, 151.2, 130.9, 129.1, 128.5, 127.7, 124.8, 124.1, 117.9, 116.0, 112.0, 110.1, 107.4, 101.0, 82.2, 58.5, 53.8, 46.6, 45.1, 36.9, 21.9.
**HRMS** (ESI+): m/z calc. for $C_{35}H_{35}N_4O_8S_2^+$ [M+H]$^+$: 703.1891, found: 703.1891.

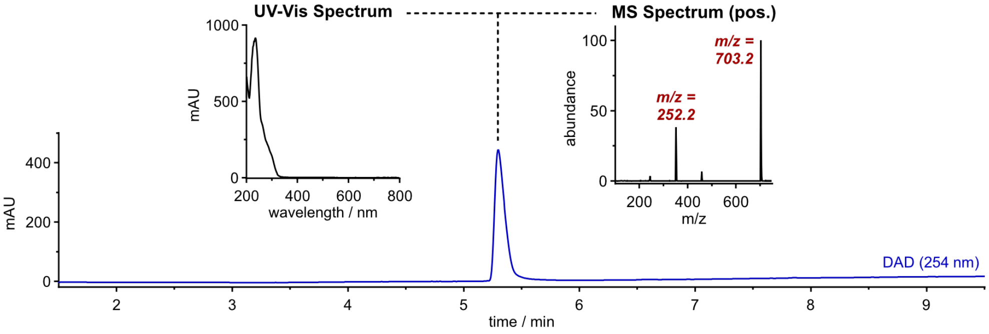

##### GL-Rho-AC$^m$ (= GL-TraG)

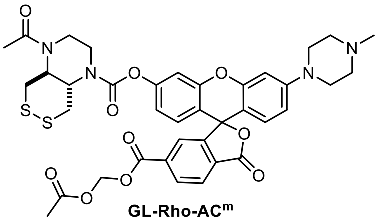

GL-Rho-AC$^m$

Prepared according to General Procedure G from **GL-Rho-AC** (5.5 mg, 7.8 µmol, 1.0 eq, 0.01 M) with 10 eq DIPEA and 5.0 eq acetoxymethyl bromide. The crude product was purified by preparative HPLC (acetonitrile/water, 0.1% formic acid; 10→50% MeCN, 20 min) affording **GL-Rho-AC$^m$** (2.2 mg, 2.7 µmol, 50%) as colourless solid.

**TLC** *Rf* = 0.24 (*np*, 3% MeOH / DCM)
**$^1$H-NMR** (400 MHz, DMSO-*d*$_6$): *δ* (ppm) = 8.27 (d, *J* = 8.1 Hz, 1H), 8.20 (d, *J* = 8.0 Hz, 1H), 7.75 (s, 1H), 7.17 (s, 1H), 6.89 (d, *J* = 8.6 Hz, 1H), 6.87 – 6.80 (m, 2H), 6.76 (d, *J* = 9.0 Hz, 1H), 6.63 (d, *J* = 8.9 Hz, 1H), 5.86 (s, 2H), 4.27 (t, *J* = 10.1 Hz, 1H), 4.09 – 3.83 (m, 2H), 3.79 – 3.52 (m, 4H), 3.24 (d, *J* = 4.7 Hz, 4H), 3.04 (d, *J* = 12.0 Hz, 1H), 2.44 – 2.39 (m, 4H), 2.21 (s, 3H), 2.06 (s, 6H).
**HRMS** (ESI+): m/z calc. for $C_{38}H_{39}N_4O_{10}S_2^+$ [M+H]$^+$: 775.2102, found: 775.2094

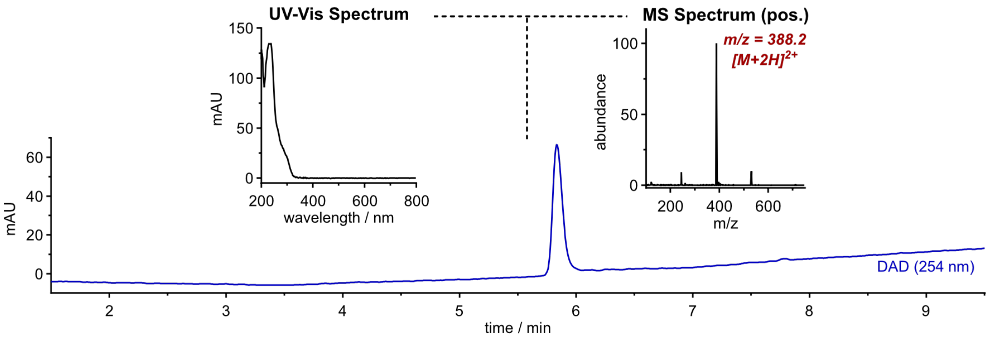

##### H-Rho-AC

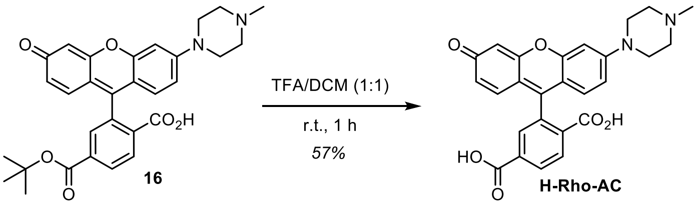

Prepared according to General Procedure F from **16** (4 mg, 7.8 µmol, 1.0 eq, 0.02 M). The crude product was purified by preparative HPLC (acetonitrile/water, 0.1% formic acid; 2→30% MeCN, 20 min) affording **H-Rho-AC** (2.0 mg, 4.4 µmol, 57%) as red solid.

**$^1$H-NMR** (400 MHz, DMSO-*d*$_6$): *δ* (ppm) = 8.18 (d, J = 9.1 Hz, 1H), 7.99 (d, J = 7.9 Hz, 1H), 7.55 (s, 1H), 6.80 (d, J = 2.3 Hz, 1H), 6.72 (dd, J = 9.0, 2.3 Hz, 1H), 6.67 (d, J = 2.2 Hz, 1H), 6.63 – 6.49 (m, 3H), 3.24 (s, 4H), 2.27 (s, 3H).
**HRMS** (ESI+): m/z calc. for $C_{26}H_{23}N_2O_6^+$ [M+H]$^+$: 459.1551, found: 459.1541.

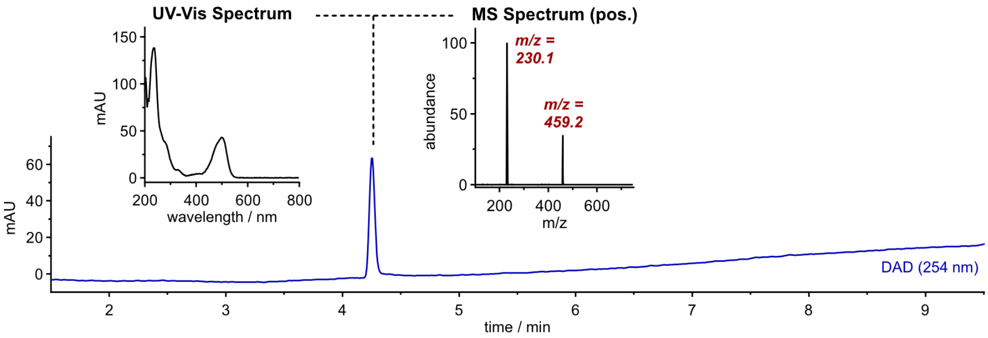

##### 7.2.5 Hydrogen Peroxide Sensor HP-TraG

**Synthetic Route for HP-TraG**

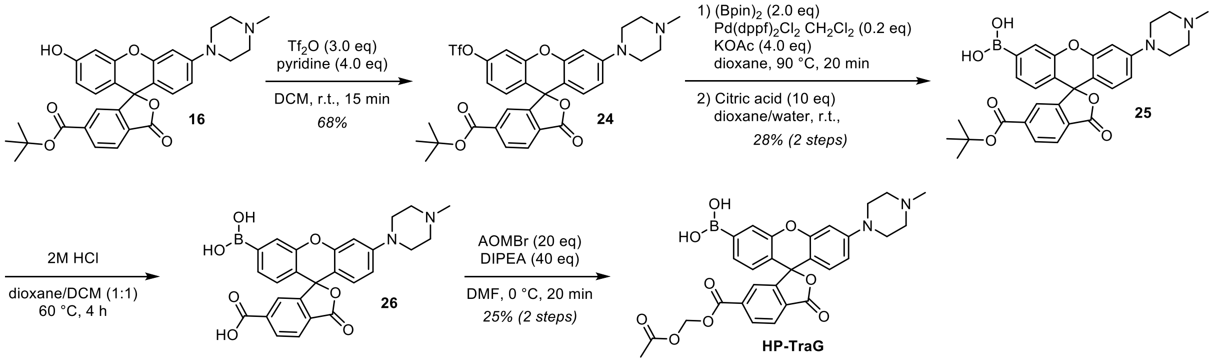

**Compound 24**

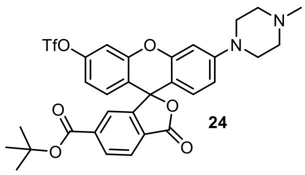

**16** (8 mg, 16 µmol, 1.0 eq) was dissolved in anhydrous DCM (1.5 mL), pyridine (5 µL, 62 µmol, 4.0 eq) and trifluoromethanesulfonic anhydride (7.8 µL, 46 µmol, 3.0 eq) were added and the reaction mixture was stirred at room temperature for 15 min. The volatiles were removed *in vacuo* and the crude product was purified by rp flash column chromatography (5→60% MeCN, acetonitrile/water, 0.1% FA) to afford **24** (6.8 mg, 11 µmol, 68%) as light-red solid.

**TLC** *Rf* = 0.18 (*np*, 4% MeOH / DCM)
**$^1$H-NMR** (600 MHz, $CD_2Cl_2$): *δ* (ppm) = 8.25 (dd, J = 8.0, 1.3 Hz, 1H), 8.06 (dd, J = 8.0, 0.7 Hz, 1H), 7.75 (dd, J = 1.2, 0.8 Hz, 1H), 7.28 (d, J = 2.5 Hz, 1H), 6.98 (dd, J = 8.8, 2.5 Hz, 1H), 6.89 (d, J = 8.8 Hz, 1H), 6.77 (d, J = 2.3 Hz, 1H), 6.68 (dd, J = 8.9, 2.3 Hz, 1H), 6.65 (d, J = 8.8 Hz, 1H), 3.40 – 3.35 (m, 4H), 2.73 (s, 4H), 2.45 (s, 3H), 1.53 (s, 9H).
**$^{13}$C-NMR** (151 MHz, $CD_2Cl_2$): *δ* (ppm) = 168.4, 164.3, 153.3, 152.7, 152.7, 152.3, 150.5, 139.2, 131.6, 130.6, 129.9, 129.1, 125.4, 125.3, 120.2, 119.9, 118.0, 117.0, 112.9, 110.9, 108.3, 102.3, 82.9, 82.6, 54.7, 47.6, 45.6, 28.1.
**HRMS** (ESI+): m/z calc. for $C_{31}H_{30}F_3N_2O_8S^+$ $[M+H]^+$: 647.1669, found: 647.1671.

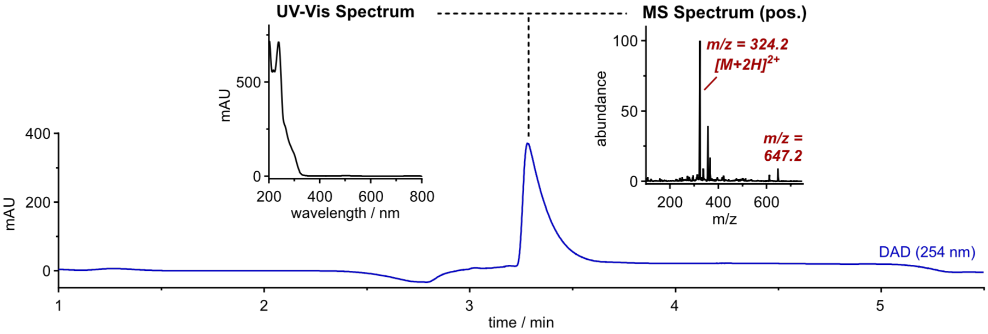

#### Compound 25

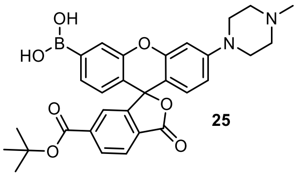

**25**

**24** (6.8 mg, 11 µmol, 1.0 eq) was dissolved in anhydrous 1,4-dioxane (1 mL) under nitrogen atmosphere and degassed by bubbling nitrogen through the solution for 2 min. Potassium acetate (4.1 mg, 42 µmol, 4.0 eq), bis(pinacolato)diboron ($(Bpin)_2$, 5.3 mg, 21 µmol, 2.0 eq) and bis(diphenylphosphino)ferrocene)palladium(II) dichloride ($Pd(dppf)Cl_2$, 1.5 mg, 2.1 µmol, 20 mol%) were added and the mixture was degassed for 5 min. Then the reaction mixture was heated to 90 °C for 20 min and allowed to cool to room temperature. Citric acid (20 mg, 0.11 mmol, 20 eq) was dissolved in water (0.2 mL), added to the reaction mixture and heated to 40 °C for 40 min. The crude product was semi-purified by preparative HPLC (acetonitrile/water, 0.1% formic acid; 5→60% MeCN, 20 min) affording **25** (1.6 mg, 3.0 µmol, 28% over 2 steps) as light-red solid.

**LRMS** (ESI+): m/z calc. for $C_{30}H_{32}BN_2O_7^+$ $[M+H]^+$: 543.2, found: 543.2.

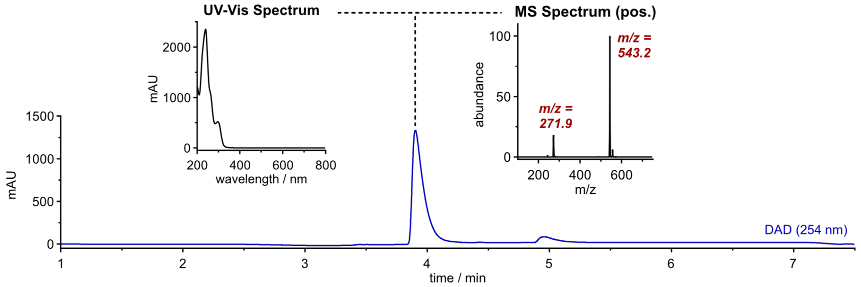

#### HP-TraG

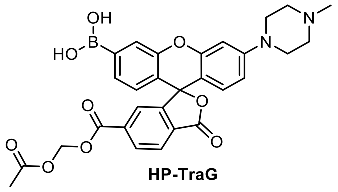

**HP-TraG**

**25** (1.6 mg, 3.0 µmol, 1.0 eq) was dissolved in anhydrous DCM (0.5 mL) and HCl (4 M in dioxane, 0.5 mL) was added. The reaction mixture was heated to 60 °C for 4 h and the volatiles were removed *in vacuo*. The intermediate product was acetoxymethylated without purification following General Procedure G (0.01 M) with 40 eq DIPEA and 20 eq acetoxymethyl bromide. The crude product was purified by preparative HPLC (acetonitrile/water, 0.1% formic acid; 5→40% MeCN, 20 min) affording **HP-TraG** (0.41 mg, 0.73 µmol, 25% over 2 steps) as light-red solid. The reaction can deliver a second species that contains an additional formaldehyde equivalent (m/z +30)) at the pendant carboxylate ($^1$H-NMR suggests an ortho-ester or acetoxymethoxymethyl ($AcOCH_2OCH_2O$-ester)), that cannot easily be separated from it by prep-HPLC and np-flash column chromatography, but which has identical performance with respect to hydrogen peroxide oxidation and esterase cleavage and therefore chromatographed **HP-TraG** can be used without any issues (see Figs S26–27).

**$^1$H-NMR** (HP-TraG): (600 MHz, DMSO-$d_6$): $\delta$ (ppm) = 8.47 (s, 2H), 8.31 – 8.25 (m, 1H), 8.23 – 8.17 (m, 1H), 7.70 (s, 2H), 7.45 (dd, J = 7.8, 1.0 Hz, 1H), 6.83 (d, J = 2.4 Hz, 1H), 6.80 – 6.71 (m, 2H), 6.63 (dd, J = 8.9, 2.4 Hz, 1H), 5.86 (s, 2H), 3.27 – 3.22 (m, 4H), 2.46 – 2.40 (m, 4H), 2.21 (s, 3H), 2.05 (s, 3H).

**$^1$H-NMR** (*side product*): (600 MHz, DMSO-$d_6$): $\delta$ (ppm) = 8.47 (s, 2H), 8.31 – 8.25 (m, 1H), 8.23 – 8.17 (m, 1H), 7.70 (s, 2H), 7.45 (dd, J = 7.8, 1.0 Hz, 1H), 6.83 (d, J = 2.4 Hz, 1H), 6.80 – 6.71 (m, 2H), 6.63 (dd, J = 8.9, 2.4 Hz, 1H), 5.54 (s, 2H), 5.32 (s, 2H), 3.27 – 3.22 (m, 4H), 2.46 – 2.40 (m, 4H), 2.21 (s, 3H), 1.71 (s, 3H).

**HRMS** (HP-TraG): (ESI+): m/z calc. for $C_{29}H_{28}BN_2O_9^+$ $[M+H]^+$: 559.1882, found: 559.1895.

**HRMS** (*side product*): (ESI+): m/z calc. for $C_{30}H_{30}BN_2O_{10}^+$ $[M+H]^+$: 589.1988, found: 589.2001.

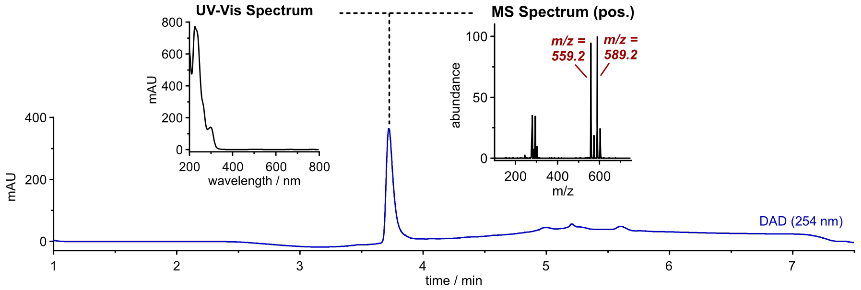

#### 7.2.6 Enzyme Activity Probe TR-TraG

##### Synthetic Route for TR-TraG

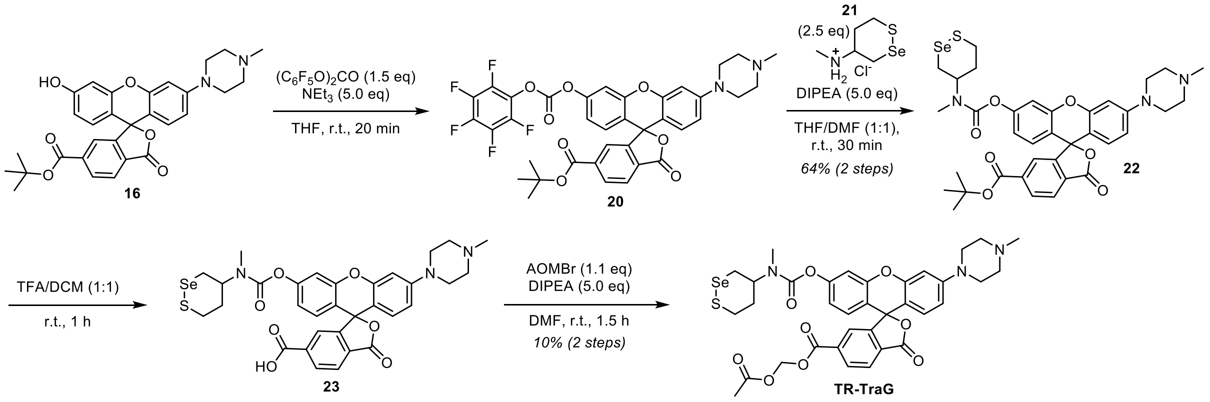

Compound **21** was prepared according to a previously described procedure (Zeisel *et al.*[29], compound S20 followed by Boc-deprotection).

##### Compound 22

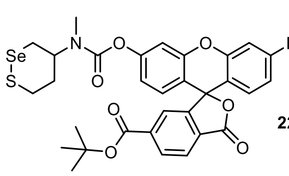

22

**16** (30 mg, 51 µmol, 1.0 eq) was dissolved in anhydrous THF (2 mL) under nitrogen atmosphere, DIPEA (86 µL, 0.51 mmol, 10 eq) and bis(pentafluorophenyl)carbonate (30 mg, 76 µmol, 1.5 eq) were added and the reaction mixture was stirred at room temperature for 20 min. **21** (9.8 mg, 42 µmol, 2.5 eq) was added as solid to the mixture, then anhydrous DMF was added. The reaction mixture was stirred at room temperature for 30 min. The volatiles were removed *in vacuo* and the crude product was semi-purified by np flash column chromatography (DCM/MeOH, 0→6% MeOH) to afford **22** (8 mg, 11 µmol, 64% over 2 steps) as light-red solid.

**TLC** *Rf* = 0.41 (*np*, 8% MeOH / DCM)
**HRMS** (SI): m/z calc. for $C_{36}H_{40}N3O_7SSe^+$ $[M+H]^+$: 738.1747, found: 738.1726.

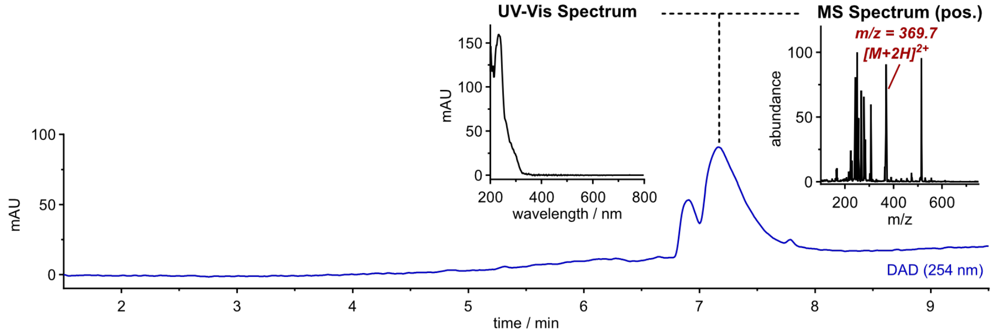

#### TR-TraG

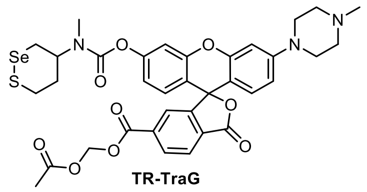

**22** (8 mg, 11 µmol, 1.0 eq, 0.02 M) was deprotected according to General Procedure F. The intermediate product was acetoxymethylated without purification following General Procedure G with 10 eq DIPEA and 5.0 eq acetoxymethyl bromide. The crude product was purified by preparative HPLC (acetonitrile/water, 0.1% formic acid; 15→50% MeCN, 20 min) affording **TR-TraG** (0.82 mg, 1.1 µmol, 10% over 2 steps) as colourless solid.

**$^{1}$H-NMR** (400 MHz, MeOD-$d_4$): $\delta$ (ppm) = 8.49 (s, 1H), 8.34 (dd, J = 8.0, 1.3 Hz, 1H), 8.13 (d, J = 8.0 Hz, 1H), 7.83 (s, 1H), 7.15 – 7.11 (m, 1H), 6.88 – 6.83 (m, 1H), 6.82 – 6.76 (m, 2H), 6.72 (dd, J = 8.9, 2.4 Hz, 1H), 6.64 (d, J = 8.9 Hz, 1H), 5.92 (s, 2H), 4.35 (s, 4H), 4.23 (d, J = 40.6 Hz, 1H), 3.51 – 3.40 (m, 1H), 3.25 (d, J = 12.6 Hz, 1H), 3.18 – 3.09 (m, 1H), 3.01 (s, 2H), 2.92 (s, 1H), 2.65 – 2.59 (m, 5H), 2.36 (s, 3H), 2.30 (d, J = 8.7 Hz, 1H), 2.07 (s, 3H).

**HRMS** (ESI+): m/z calc. for $C_{35}H_{36}N_3O_9SSe^+$ [M+H]$^+$: 754.1332, found: 754.1320.

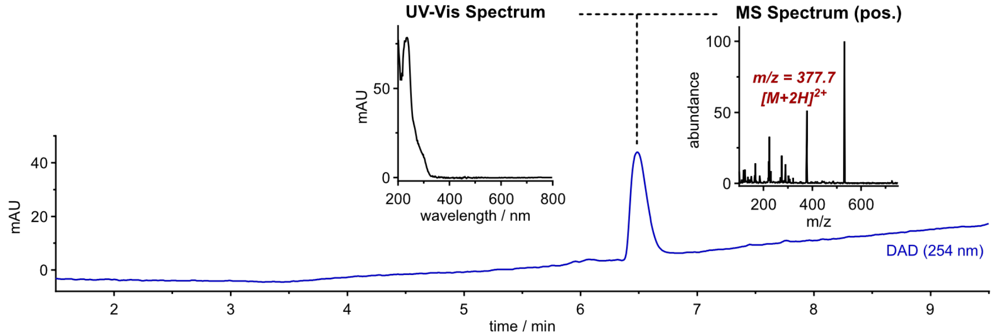

### 9 NMR spectra

| | |
|---|---|
| iC4-FS | 53 |
| iC7-FS | 54 |
| iC10-FS | 55 |
| iC4-FC | 56 |
| iC4-F | 57 |
| H-C4-FS | 58 |
| H-C7-FS | 59 |
| H-C10-FS | 60 |
| H-C4-FC | 61 |
| H-C4-F | 62 |
| GL-C4-FC | 63 |
| H-Rho | 65 |
| GL-Rho | 66 |
| H-Rho-A | 67 |
| GL-Rho-A | 68 |
| Compound 12 | 69 |
| GL-Rho-C | 70 |
| GL-Rho-Cm | 71 |
| H-Rho-C | 72 |
| Compound 16 | 73 |
| GL-Rho-AC | 74 |
| GL-Rho-ACm (= GL-TraG) | 75 |
| H-Rho-AC (= H-TraG) | 75 |
| Compound 24 | 76 |
| HP-TraG | 77 |
| TR-TraG | 77 |

**iC4-FS**

**$^{1}$H-NMR**

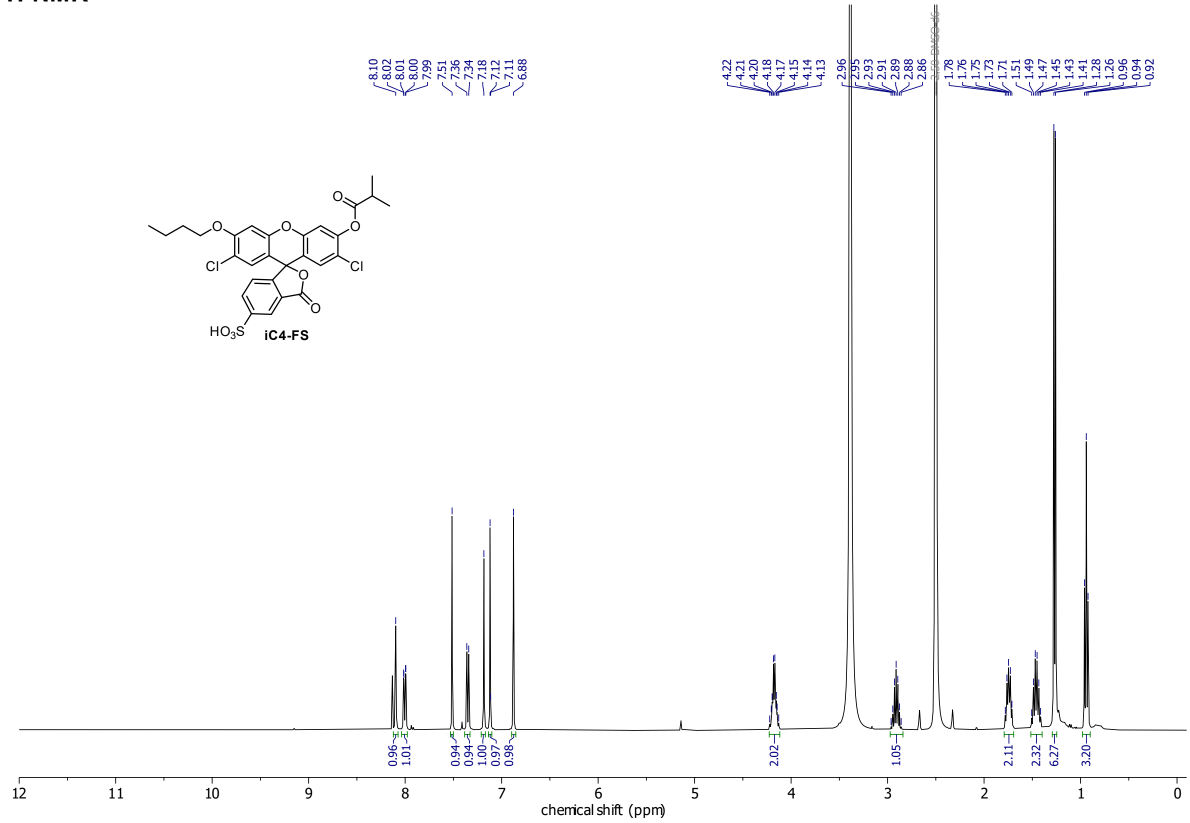

**$^{13}$C-NMR**

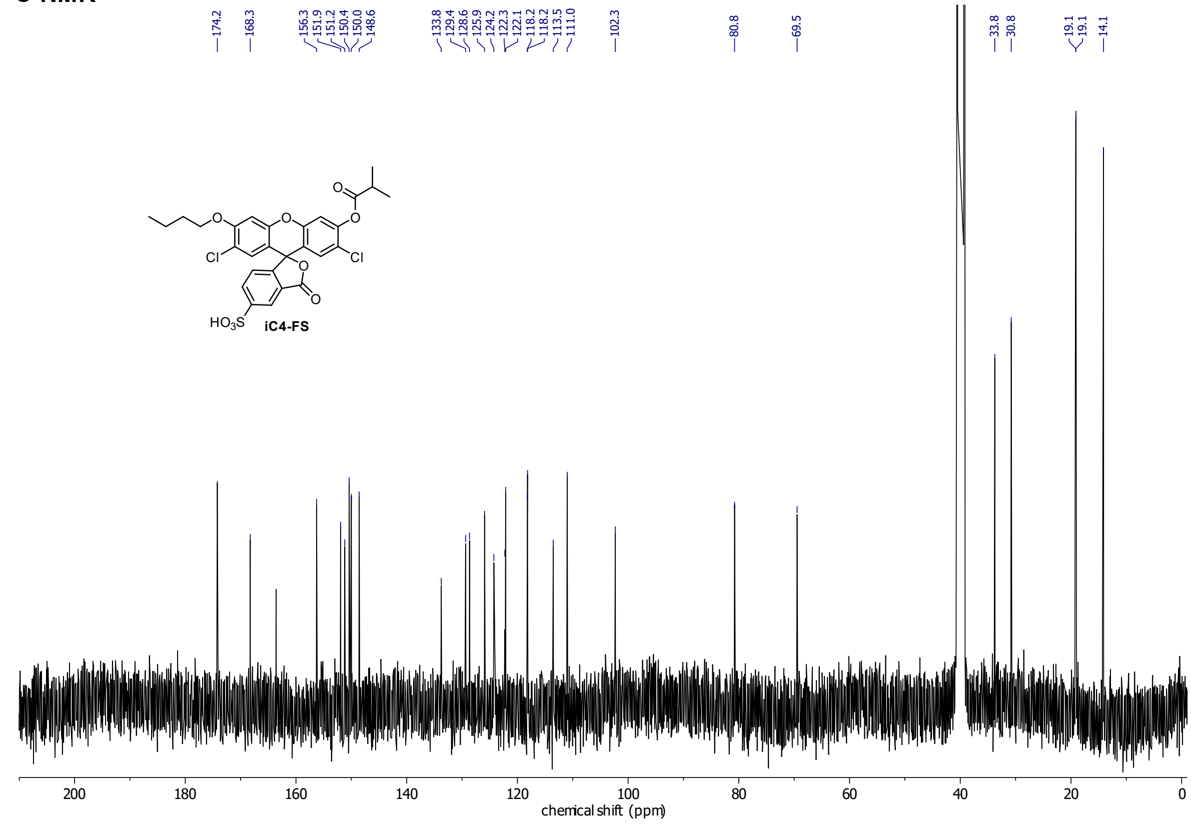

**iC7-FS**

**¹H-NMR**

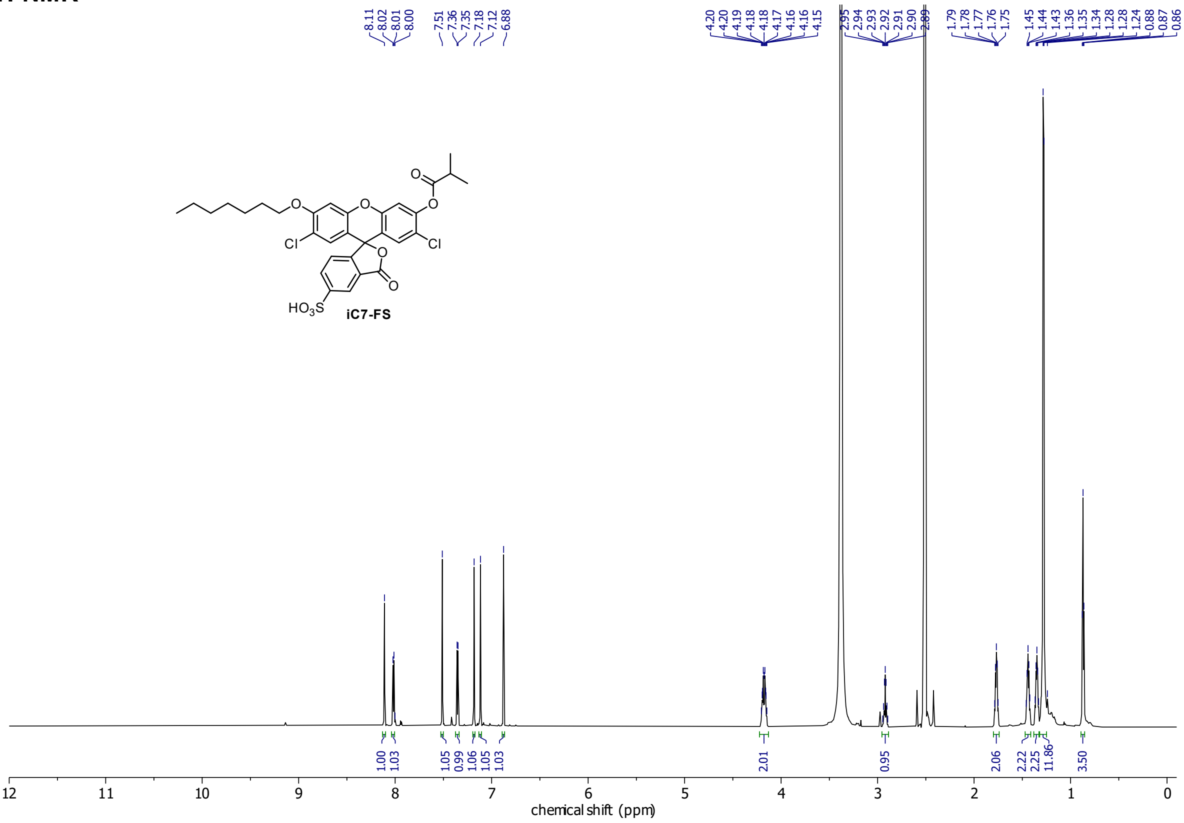

**¹³C-NMR**

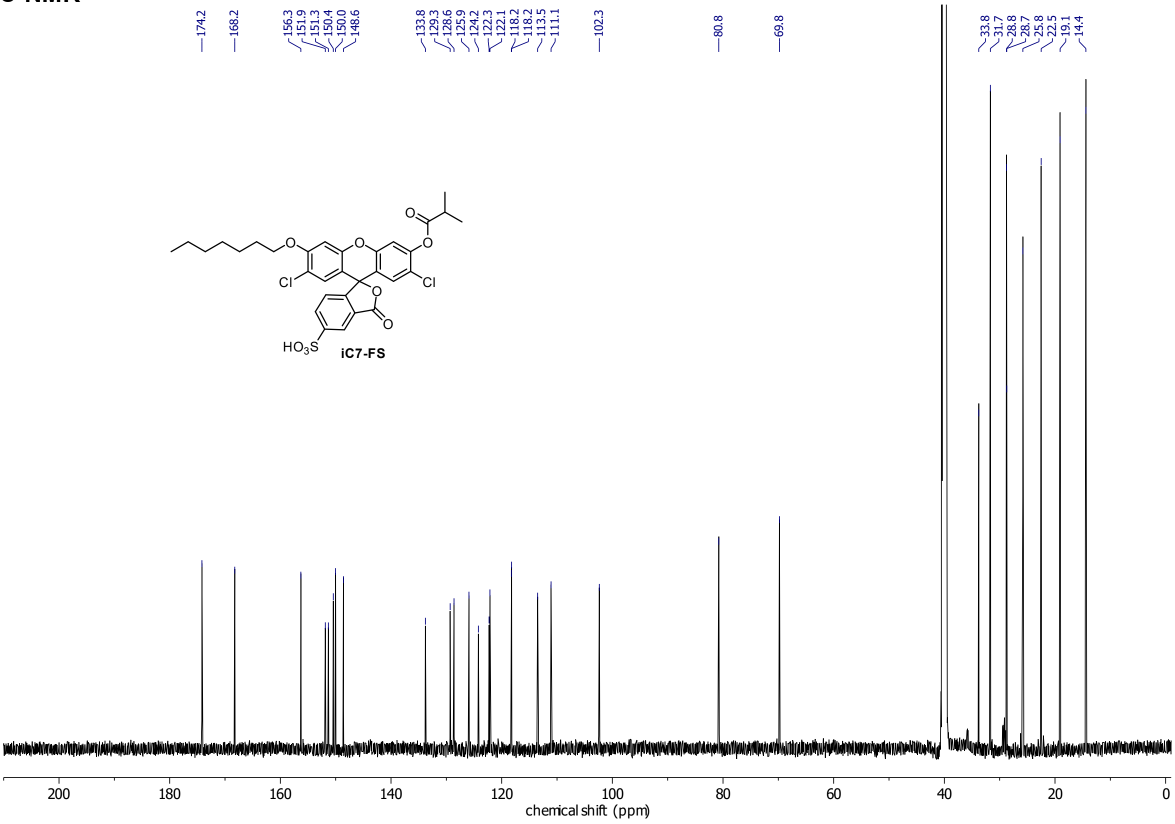

**iC10-FS**

**$^{1}$H-NMR**

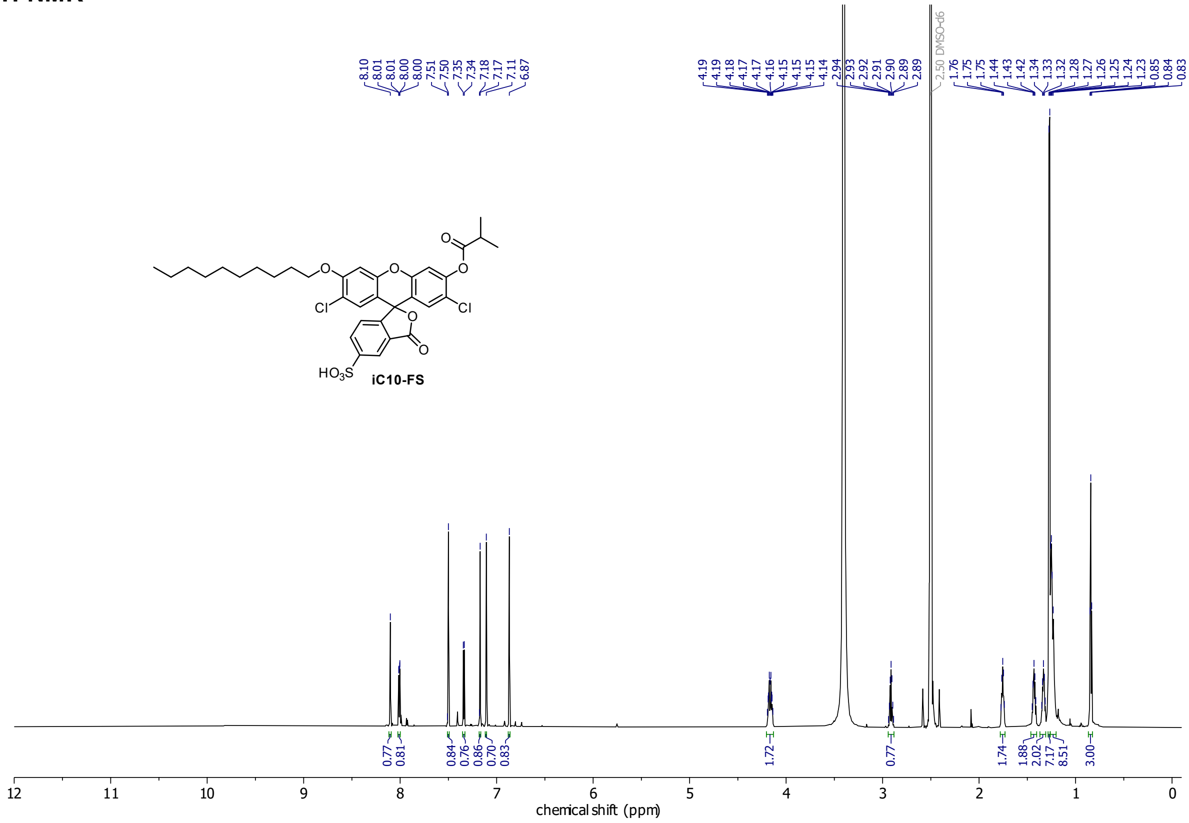

**$^{13}$C-NMR**

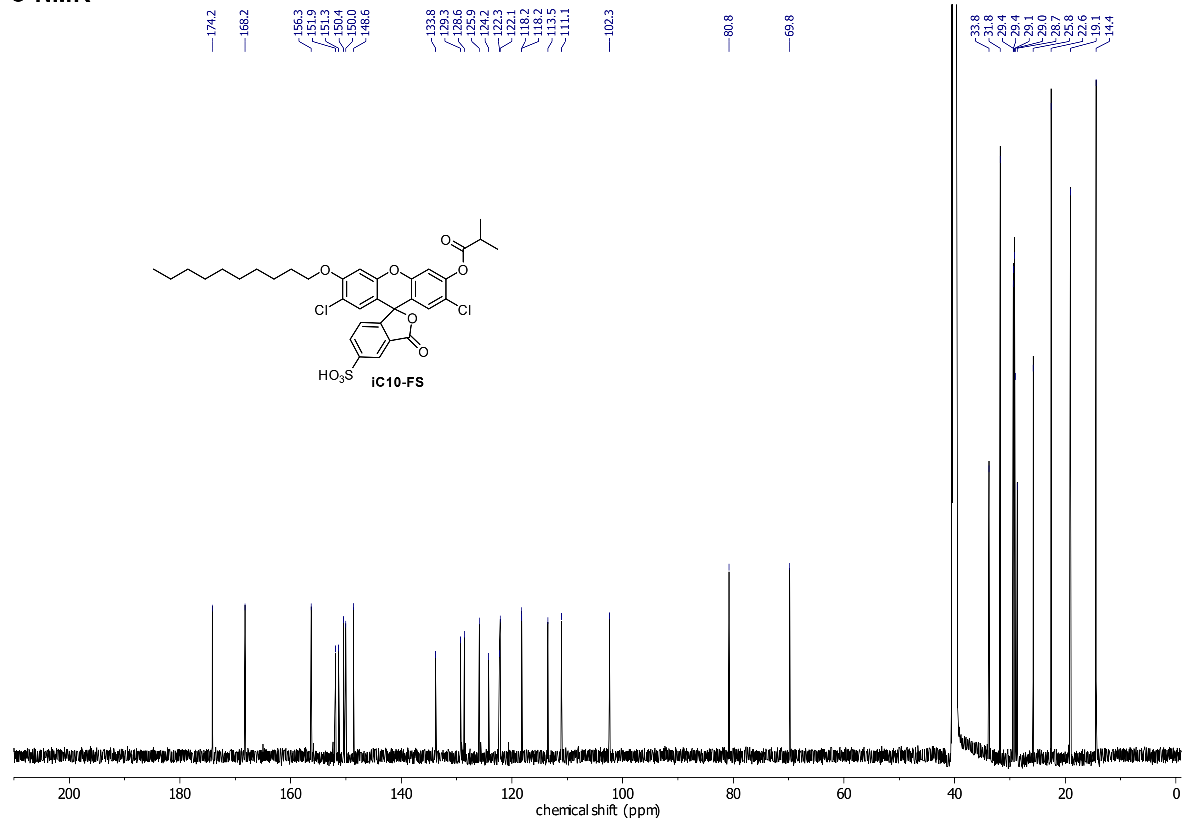

## iC4-FC

##### ¹H-NMR

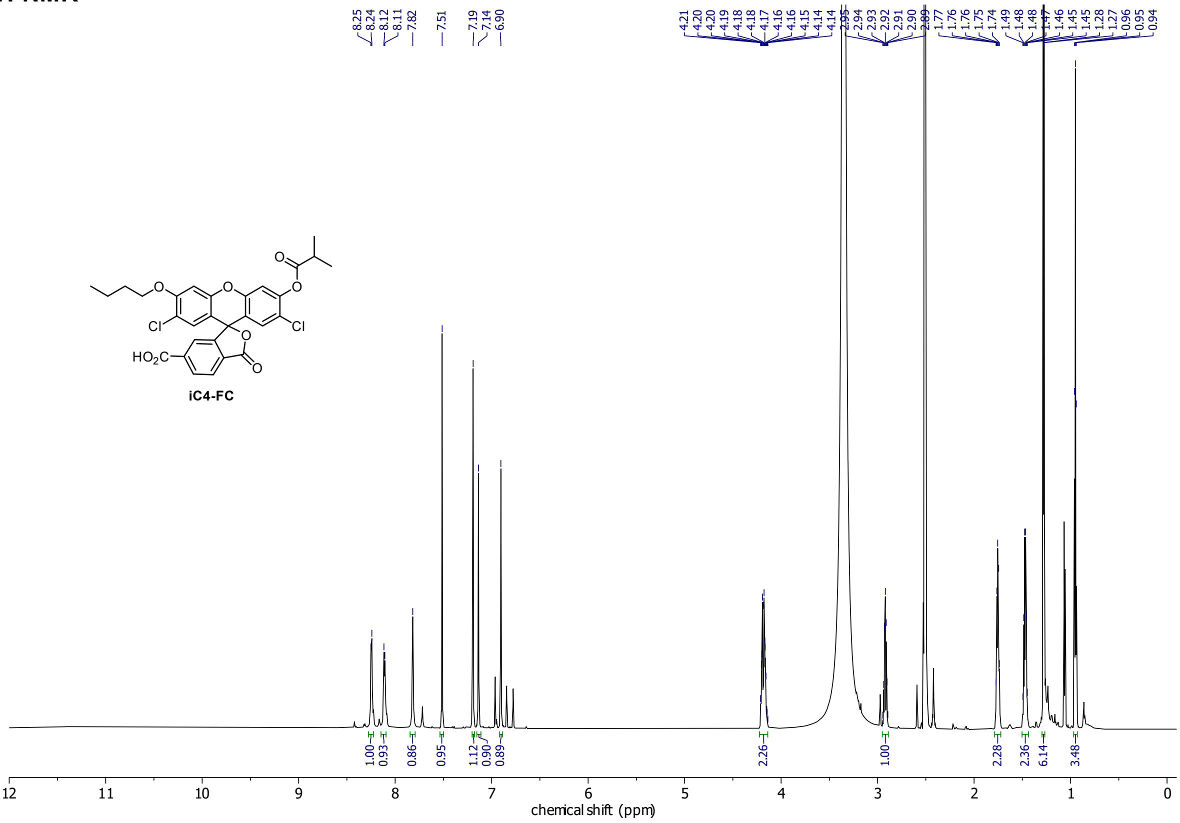

##### ¹³C-NMR

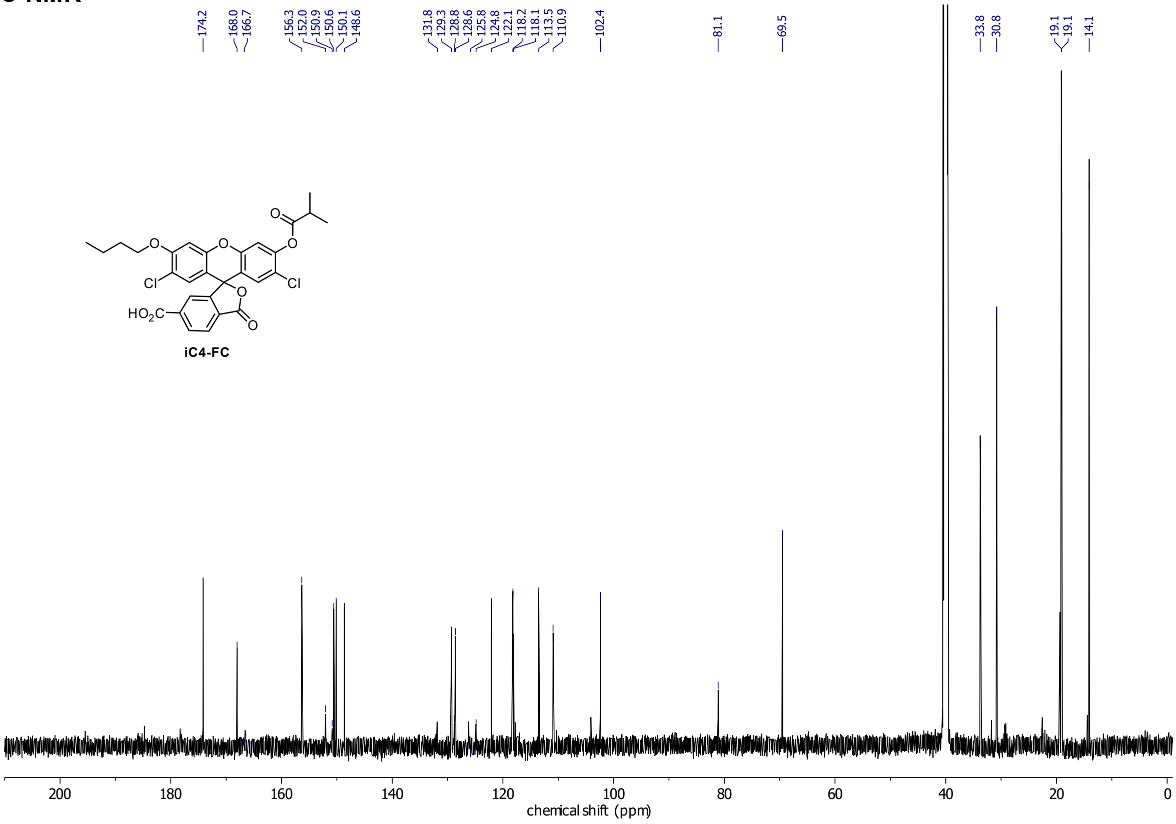

**iC4-F**

**$^{1}$H-NMR**

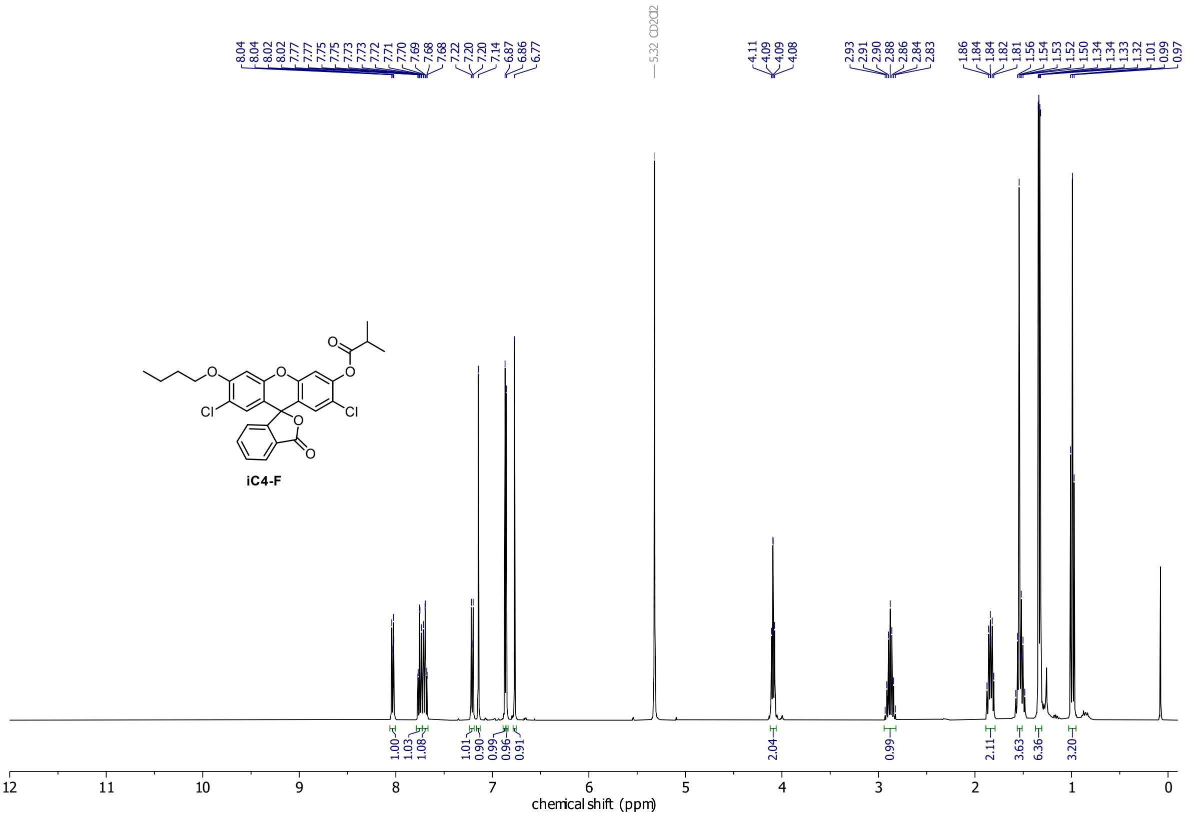

**$^{13}$C-NMR**

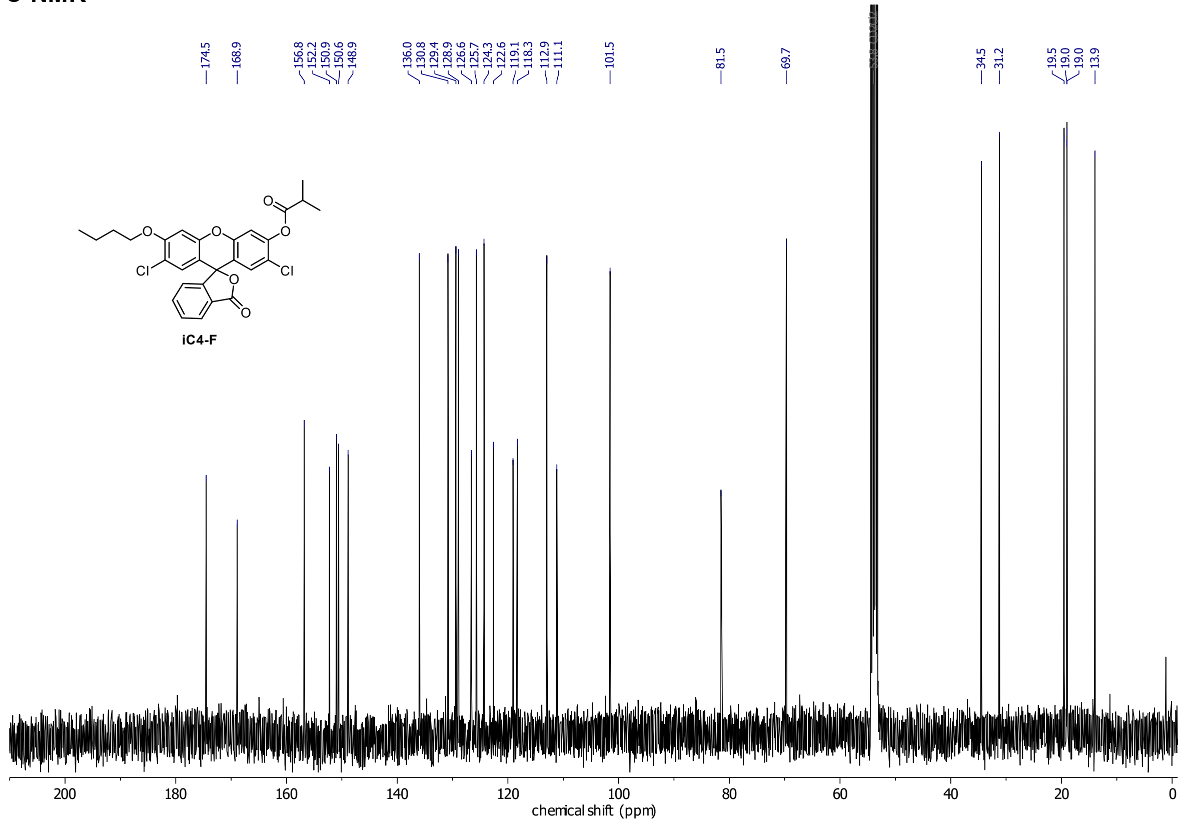

## H-C4-FS

##### $^1$H-NMR

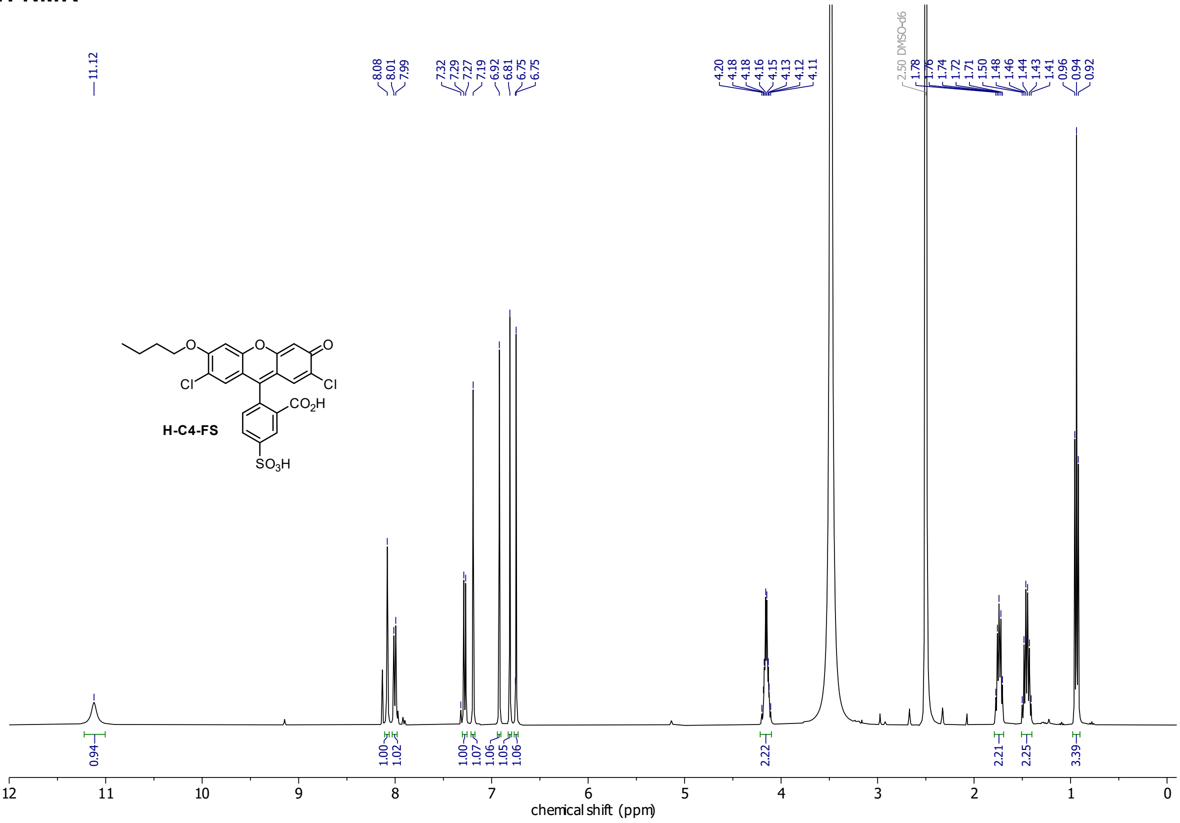

##### $^{13}$C-NMR

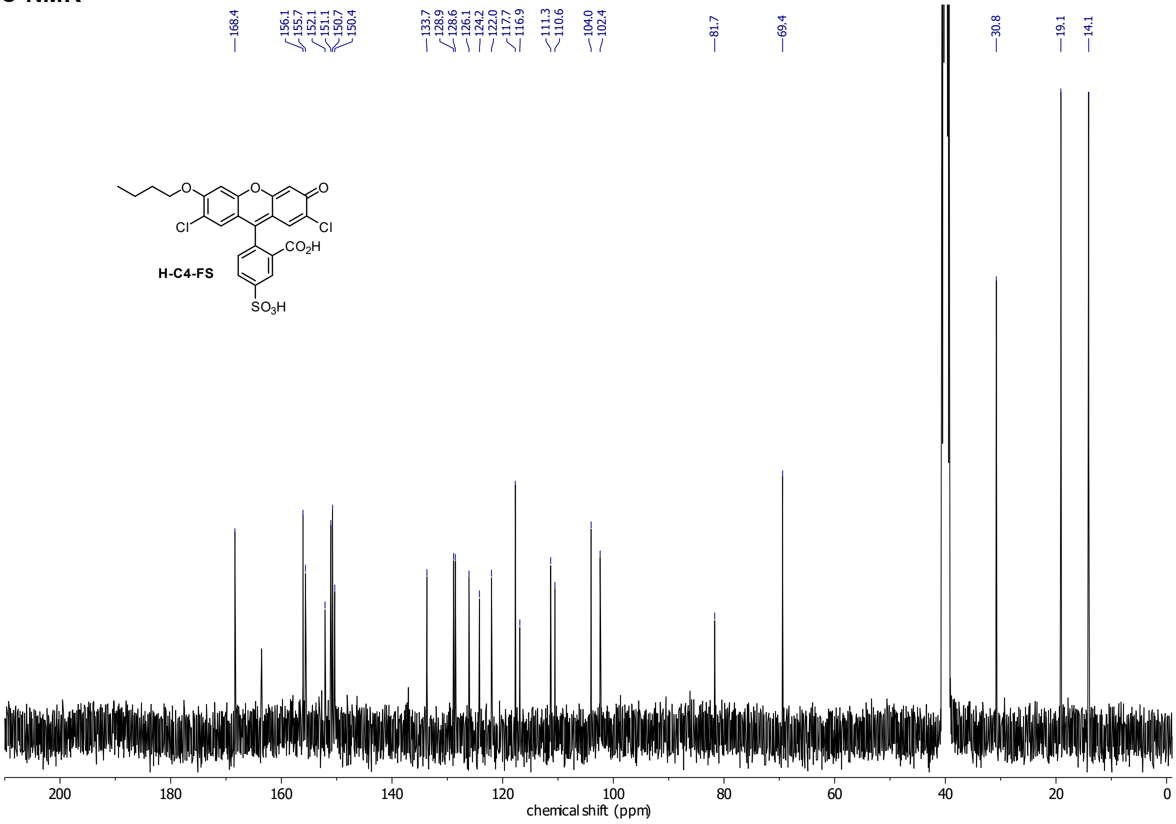

**H-C7-FS**

**$^{1}$H-NMR**

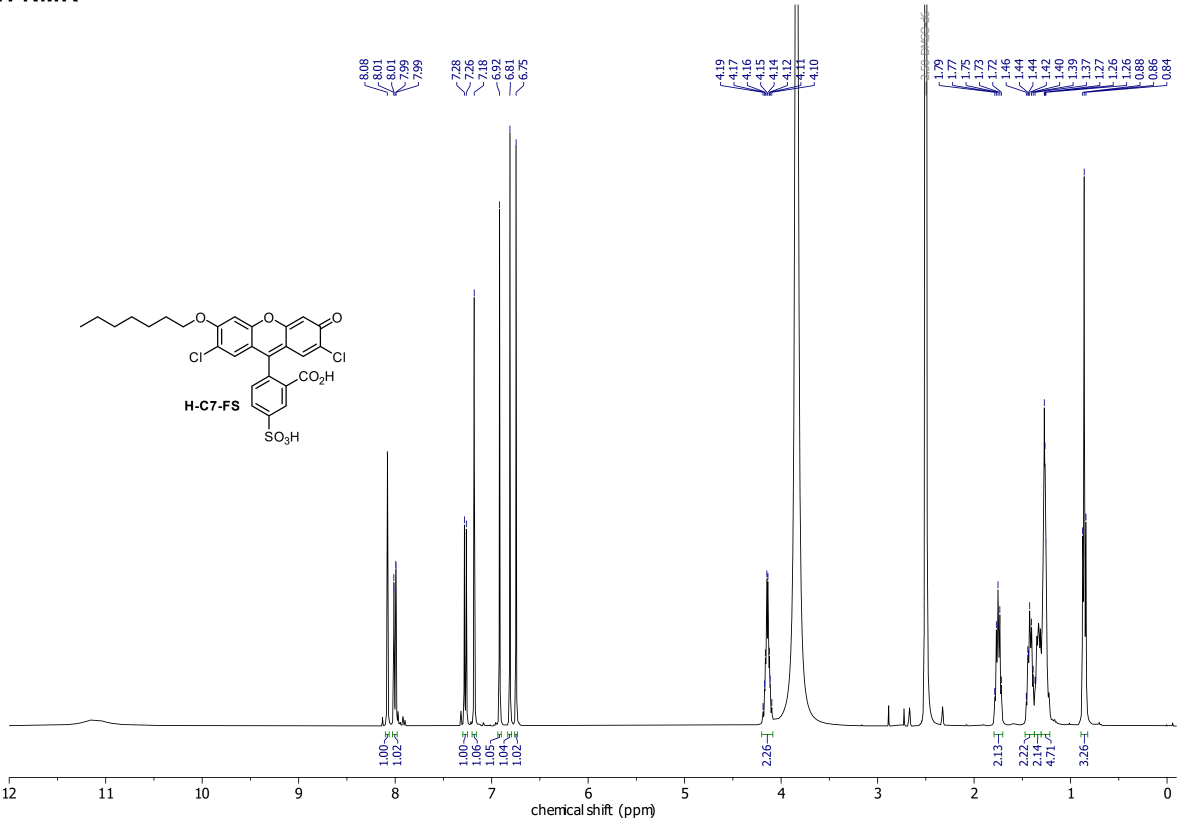

**$^{13}$C-NMR**

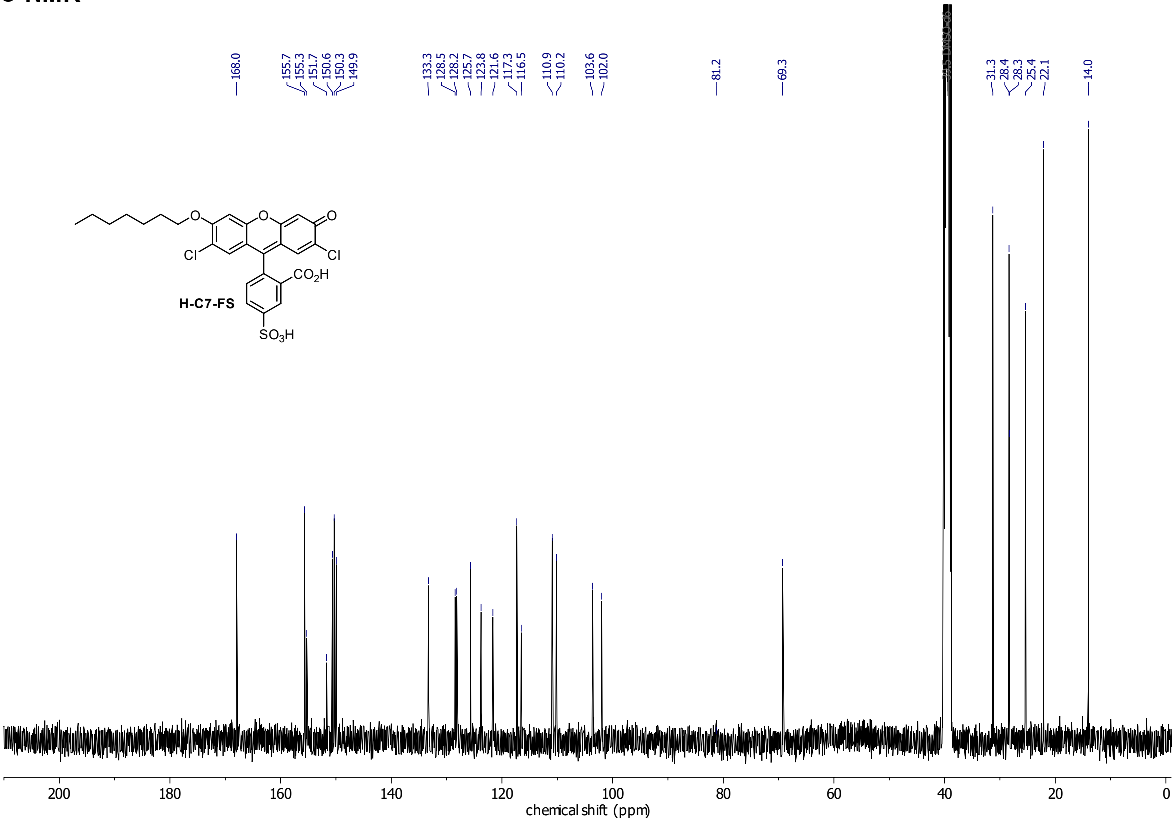

## H-C10-FS

#### $^{1}$H-NMR

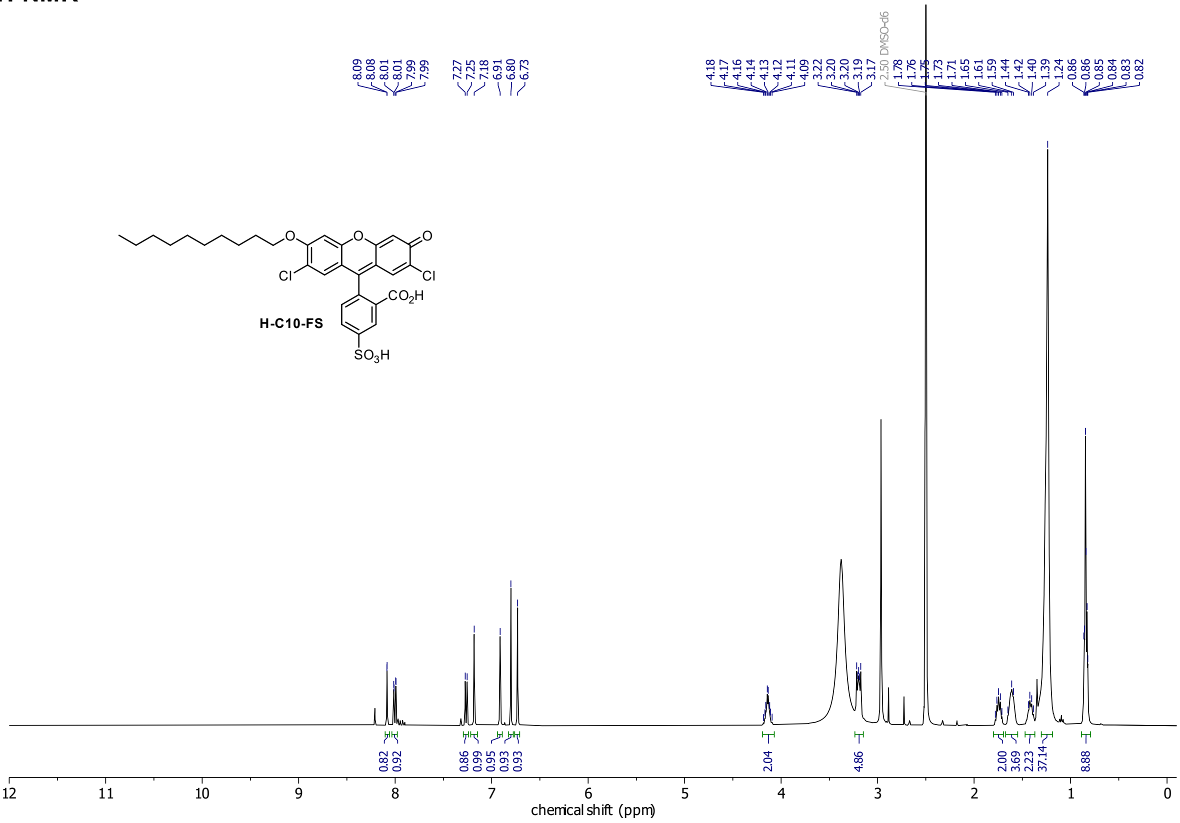

#### $^{13}$C-NMR

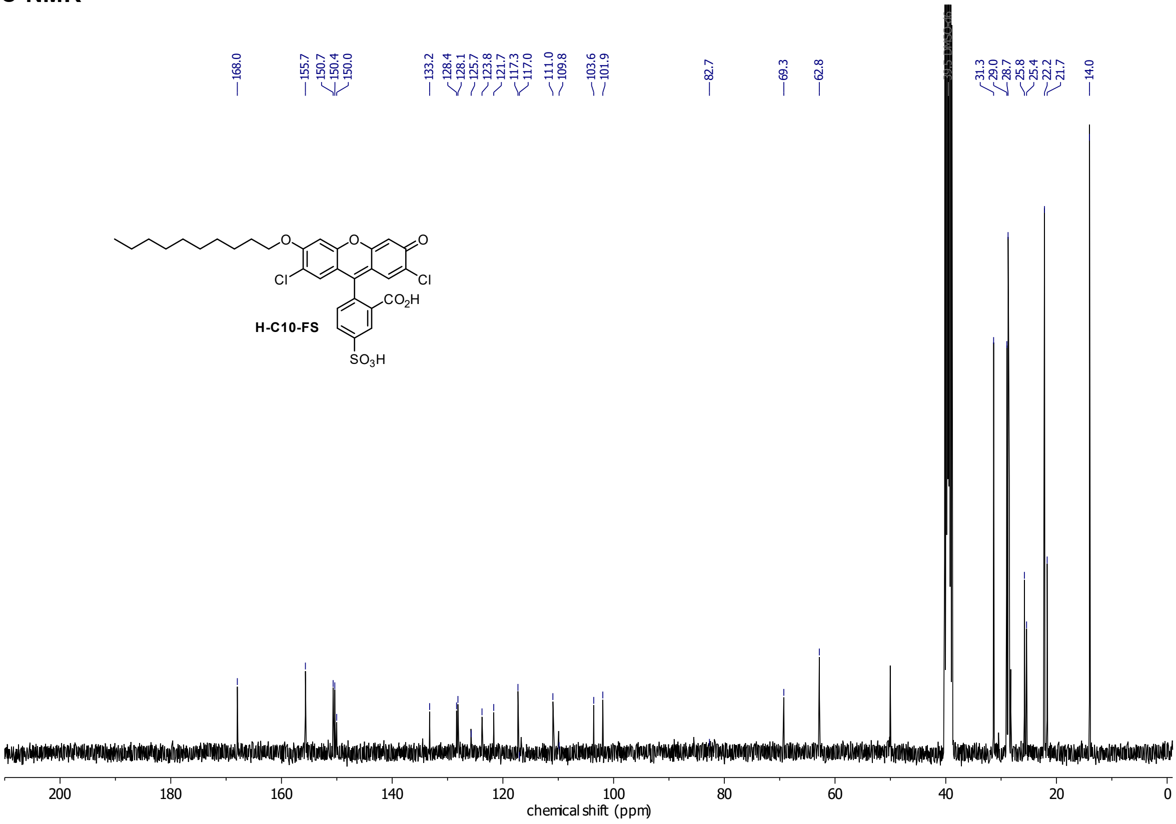

## H-C4-FC

##### $^{1}$H-NMR

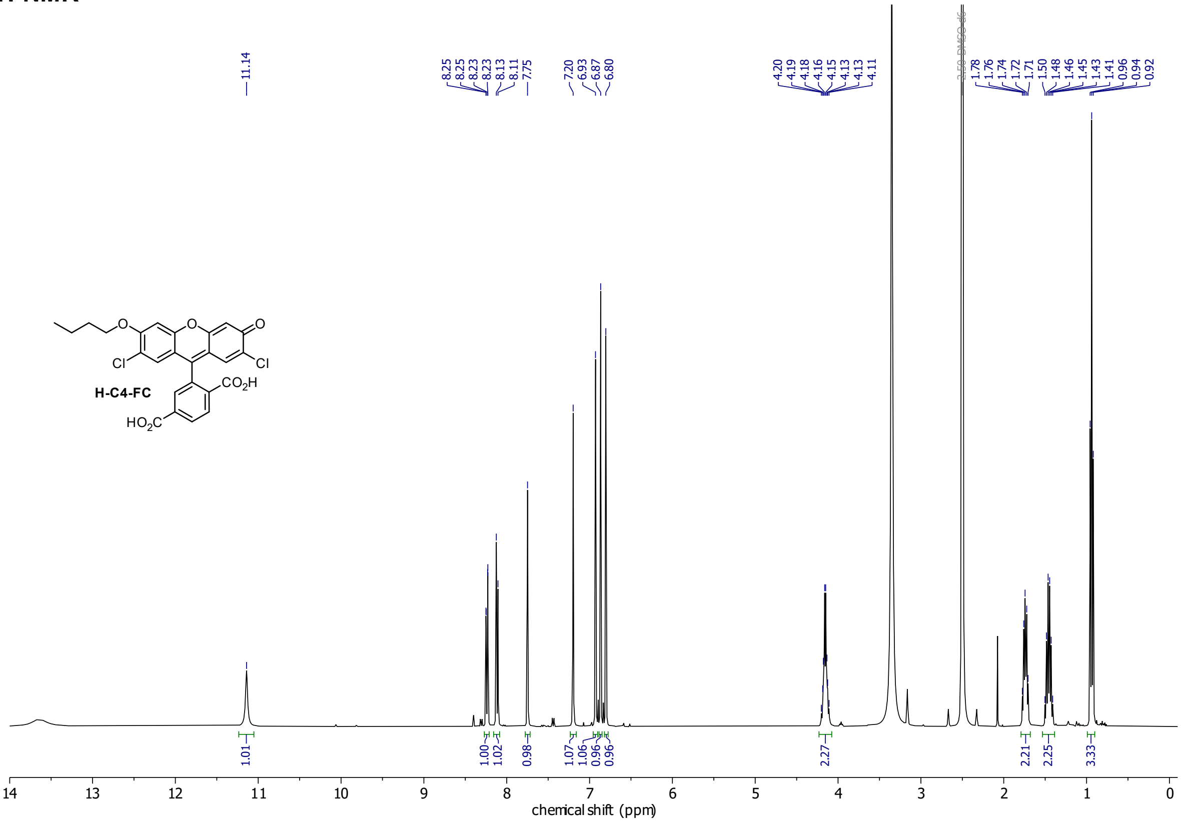

##### $^{13}$C-NMR

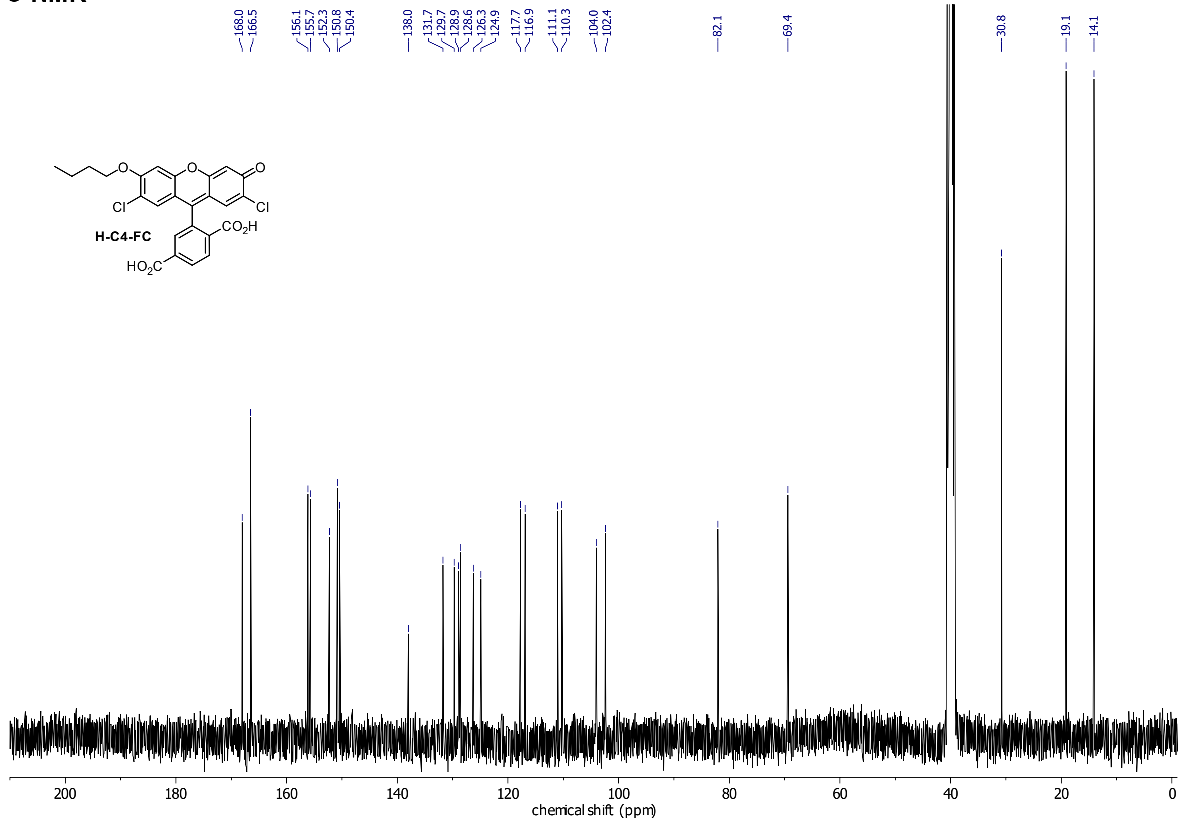

**H-C4-F**

**$^{1}$H-NMR**

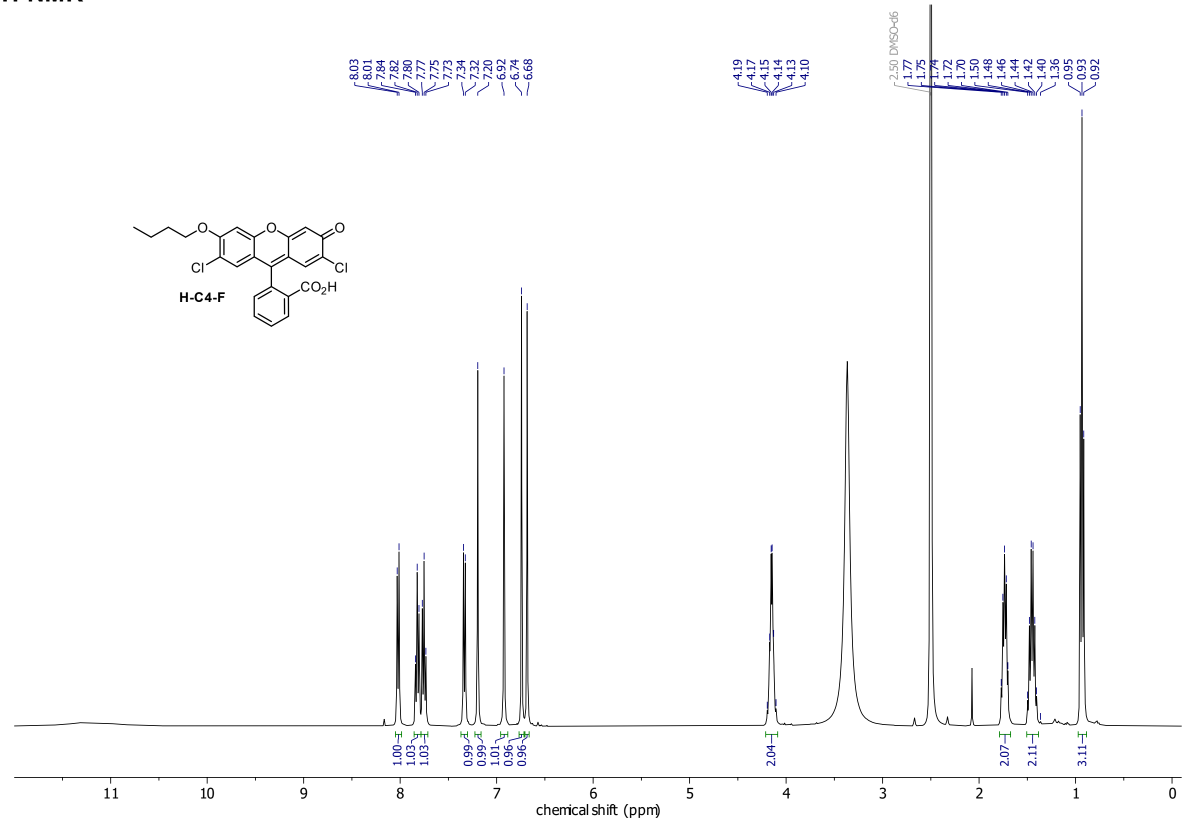

**$^{13}$C-NMR**

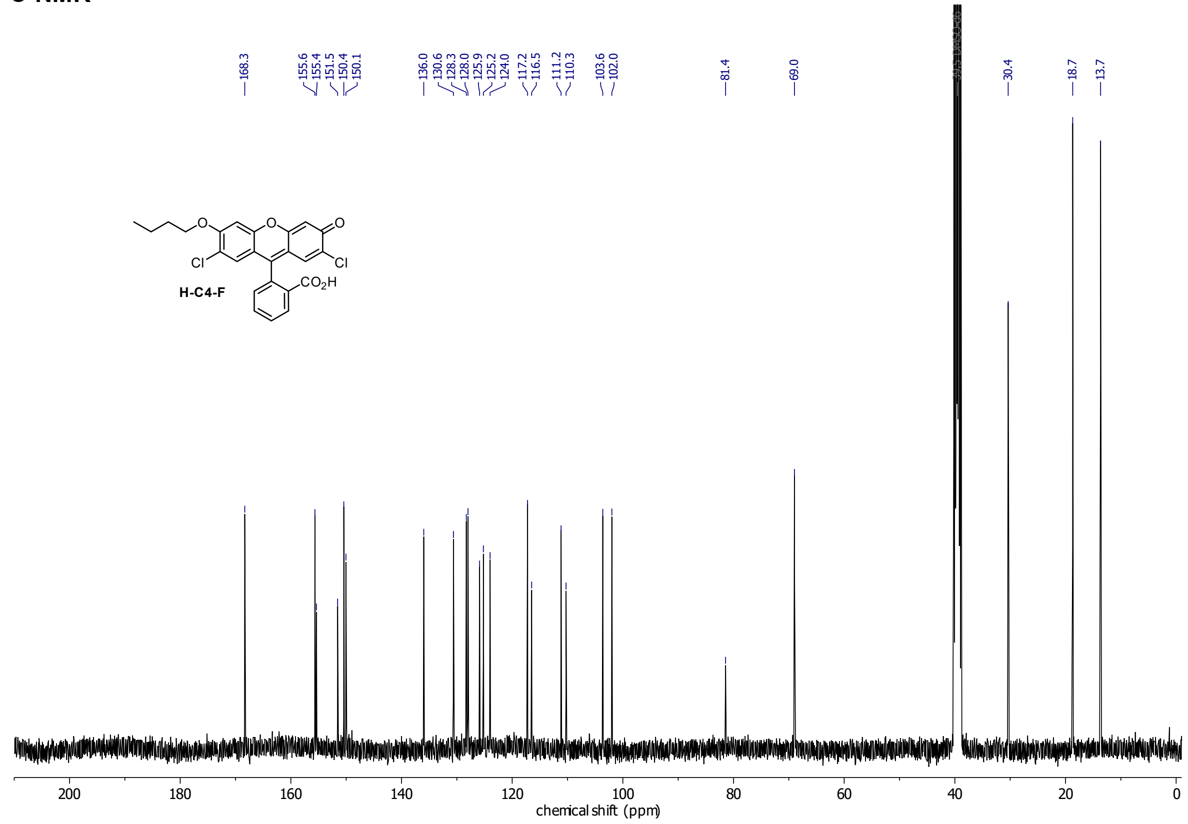

**GL-C4-FC**

**$^1$H-NMR**

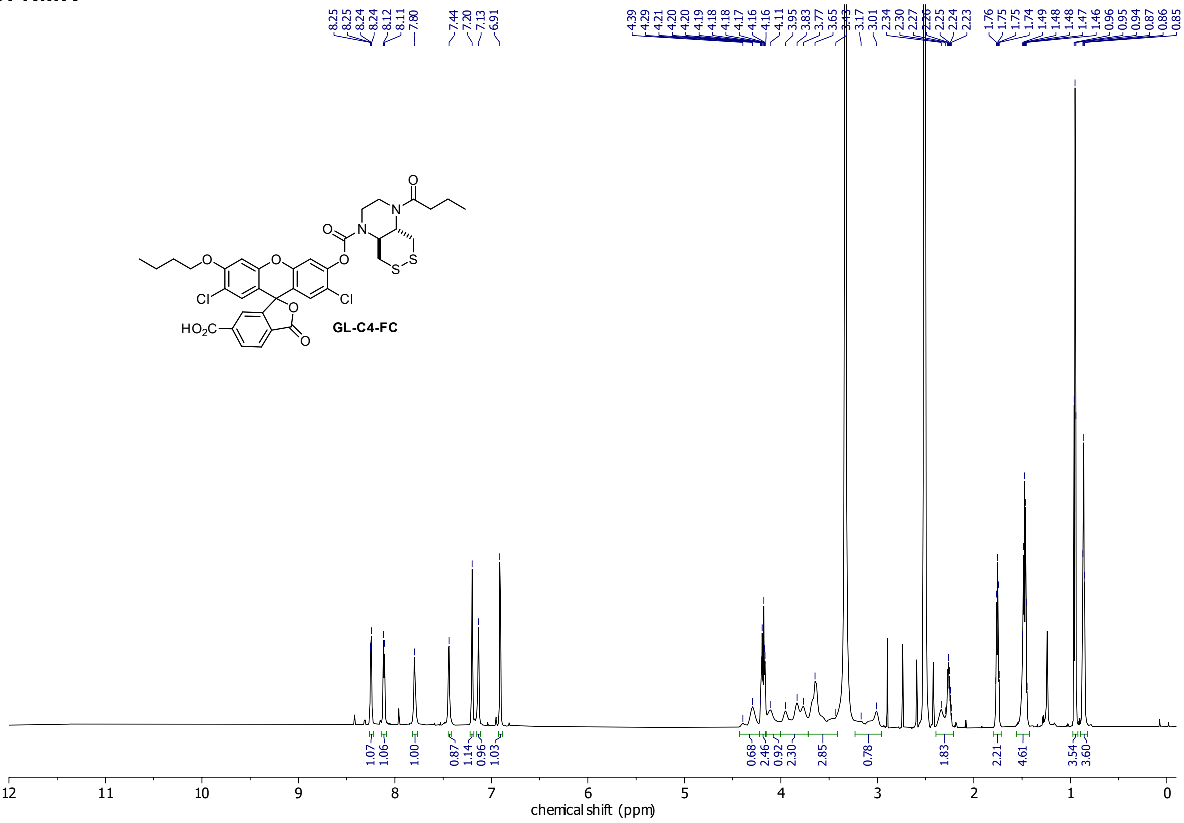

**$^{13}$C-NMR**

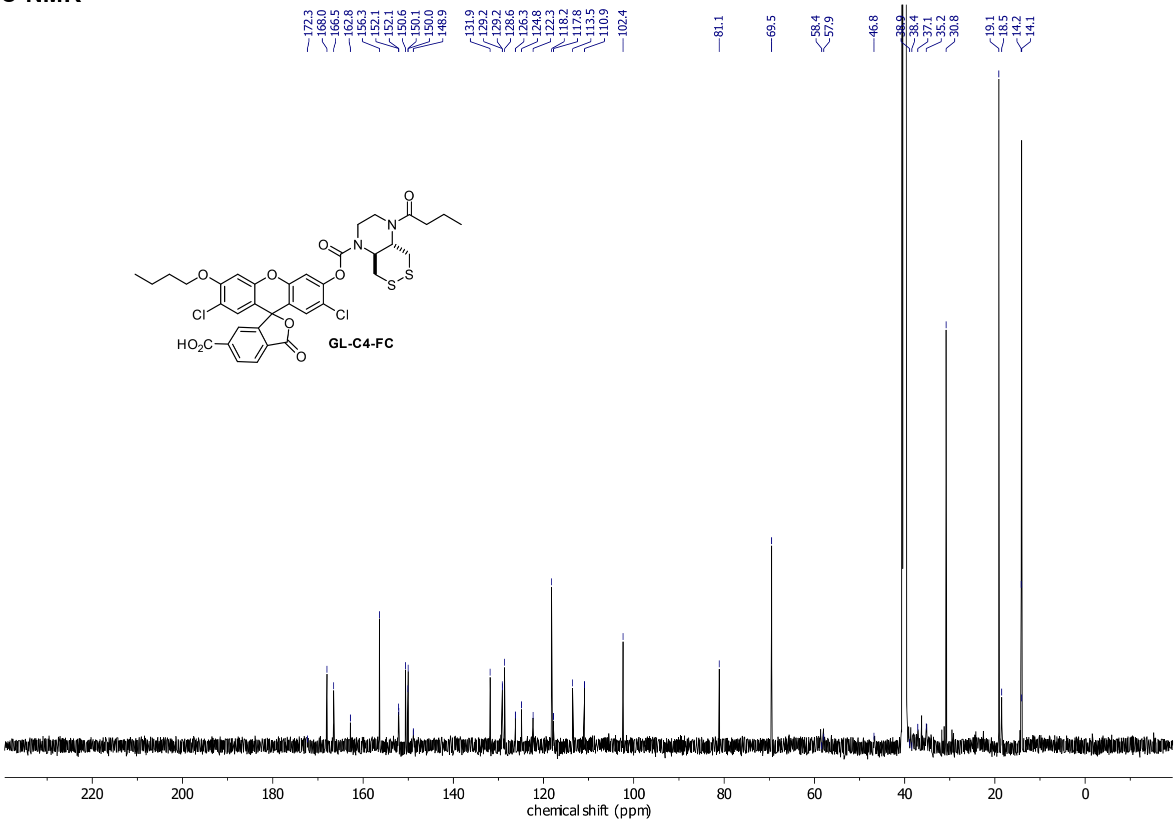

#### HSQC

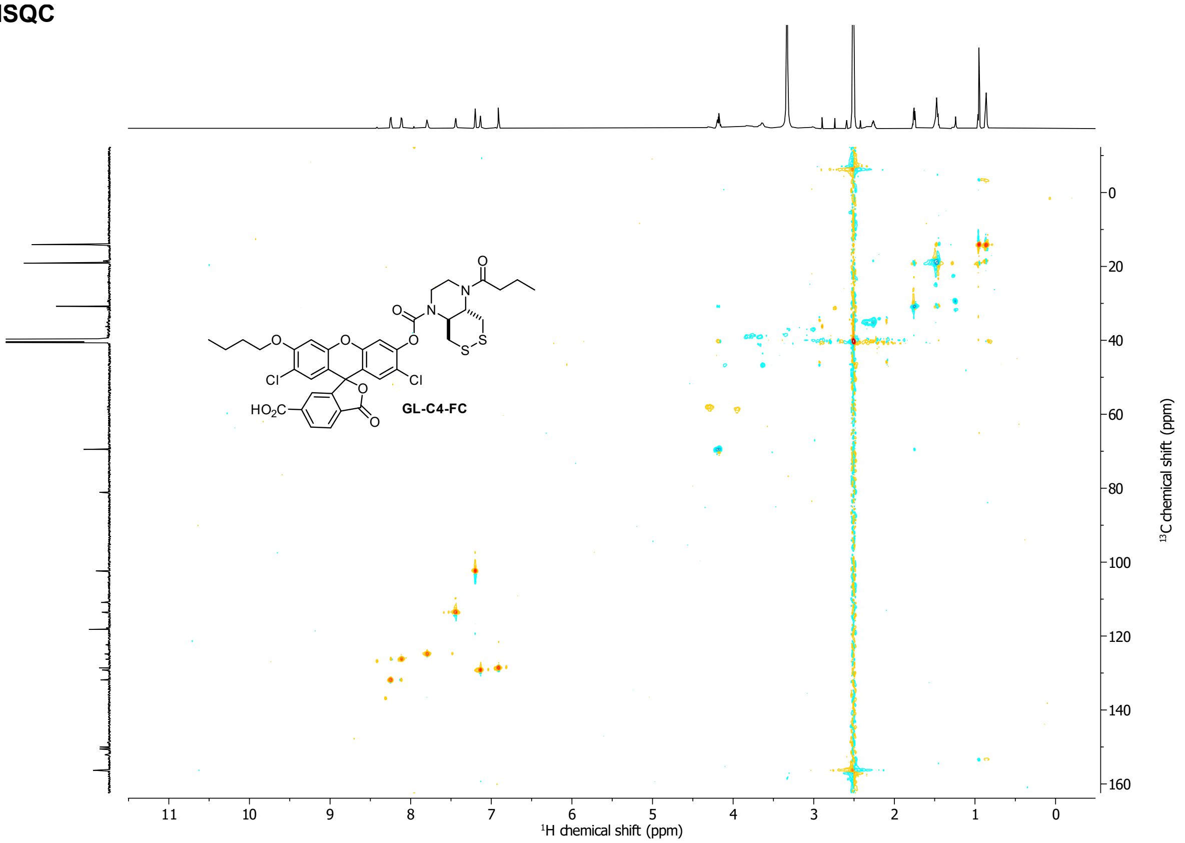
GL-C4-FC
HO2C
Cl
Cl
O
N
S
11 10 9 8 7 6 5 4 3 2 1 0
1H chemical shift (ppm)
0 20 40 60 80 100 120 140 160
13C chemical shift (ppm)

#### HMBC

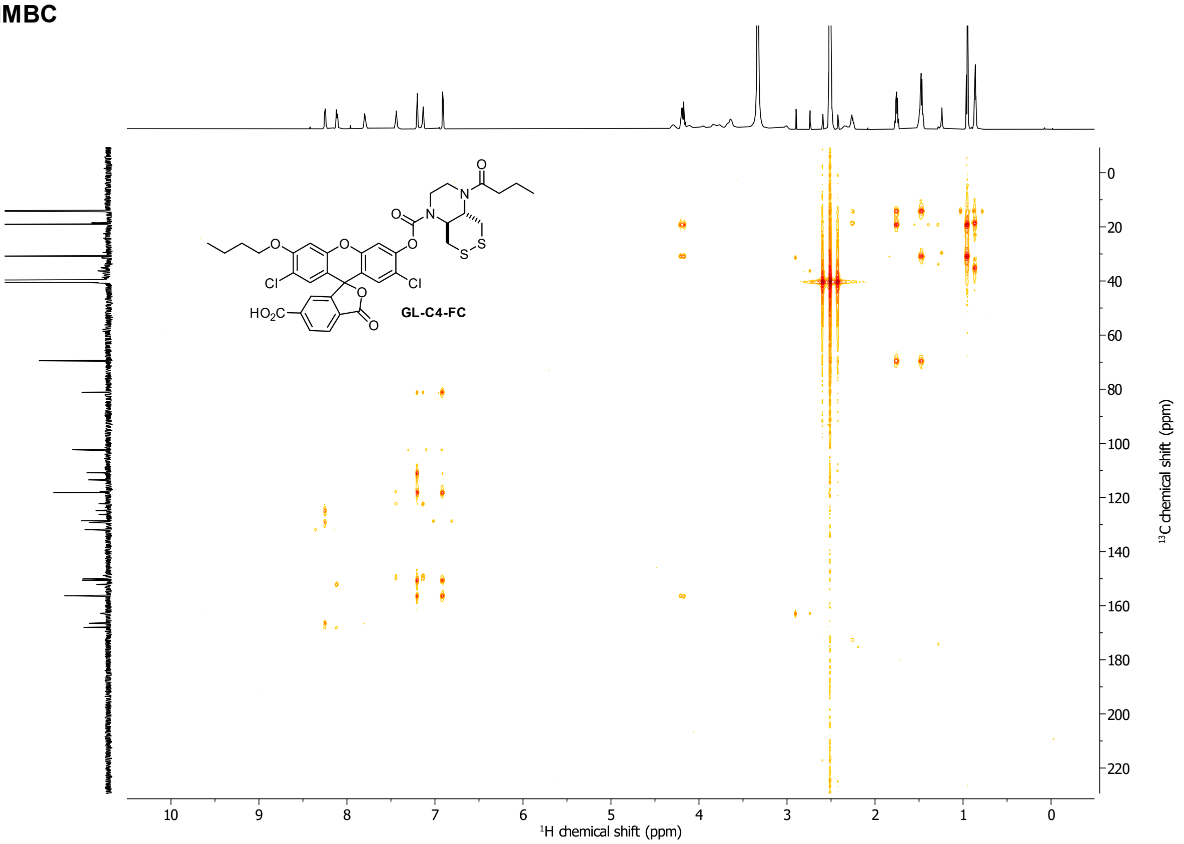
GL-C4-FC
HO2C
Cl
Cl
O
N
S
10 9 8 7 6 5 4 3 2 1 0
1H chemical shift (ppm)
0 20 40 60 80 100 120 140 160 180 200 220
13C chemical shift (ppm)

**H-Rho**

**$^{1}$H-NMR**

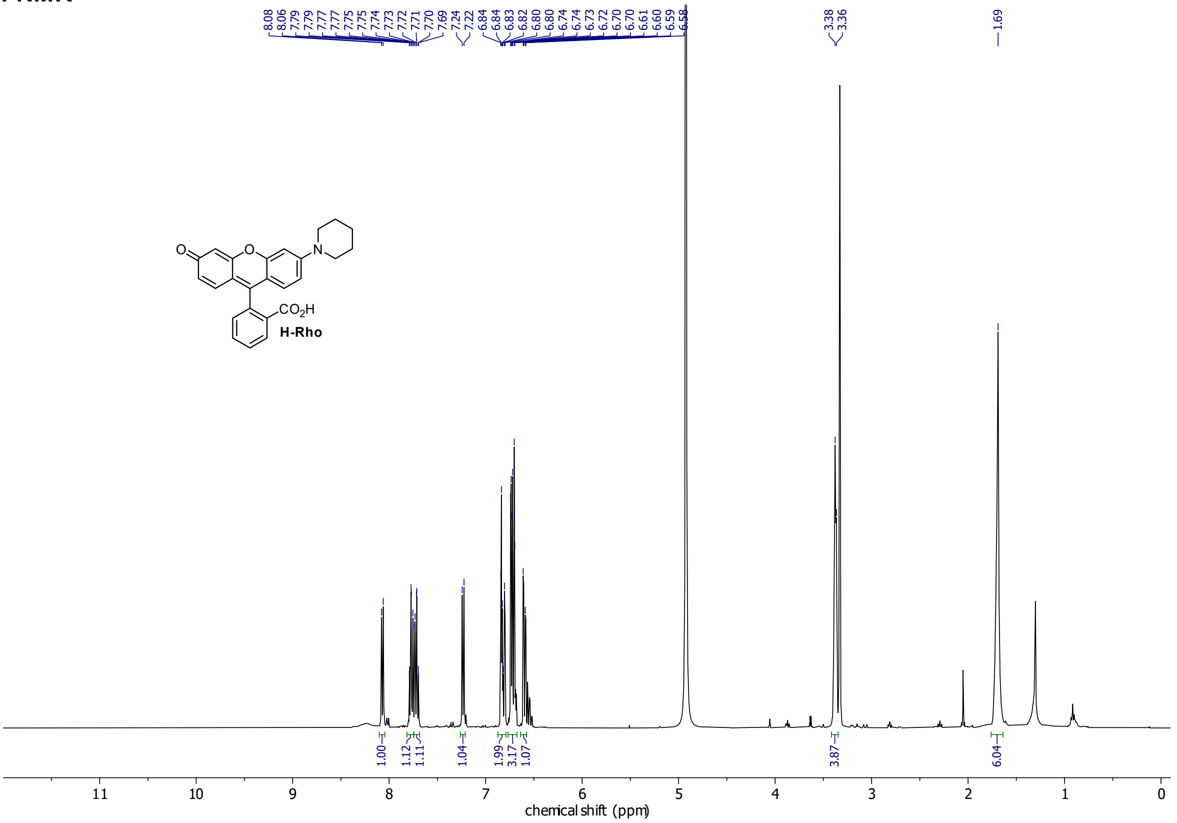
8.08
8.06
7.79
7.79
7.77
7.77
7.75
7.75
7.74
7.73
7.72
7.71
7.70
7.69
7.24
7.22
6.84
6.84
6.83
6.82
6.80
6.80
6.80
6.74
6.73
6.72
6.70
6.70
6.61
6.60
6.59
6.58
3.38
3.36
1.69
O
O
N
CO2H
H-Rho
1.00
1.12
1.11
1.04
1.99
3.17
1.07
3.87
6.04
11
10
9
8
7
6
5
4
3
2
1
0
chemical shift (ppm)

**GL-Rho**

**$^{1}$H-NMR**

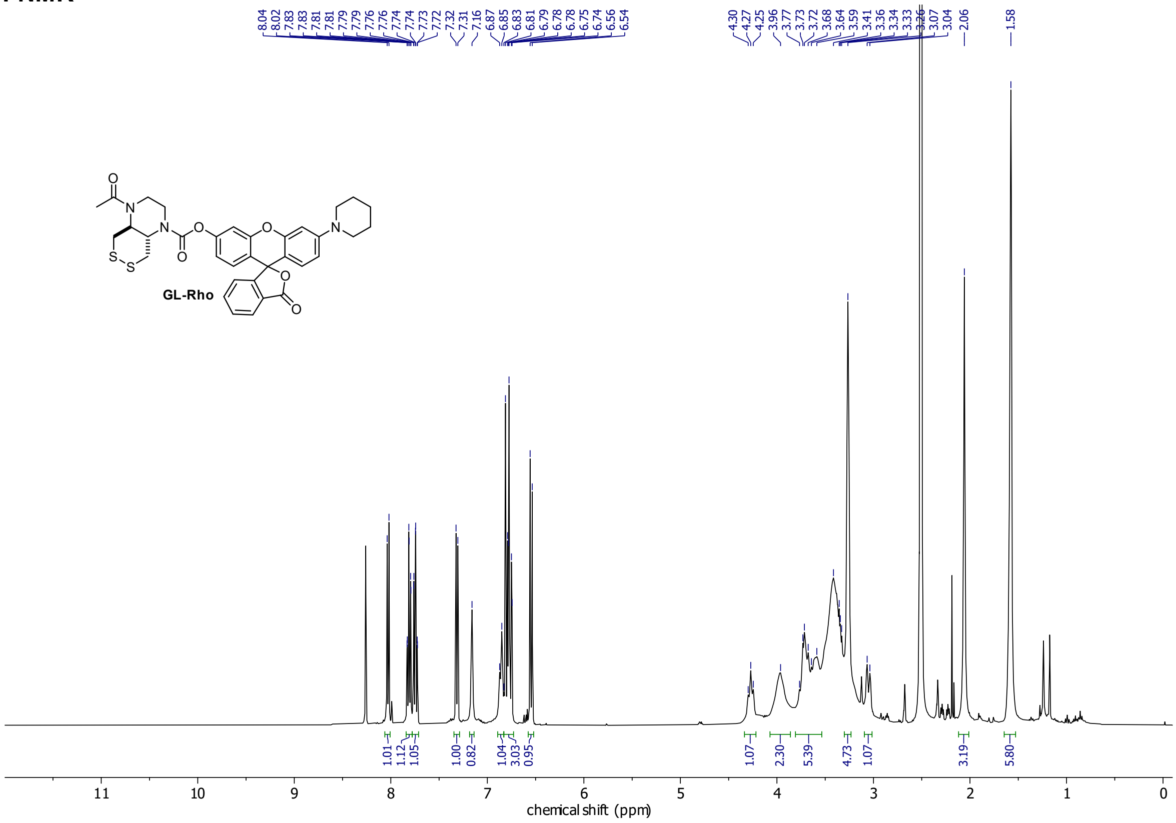

**$^{13}$C-NMR**

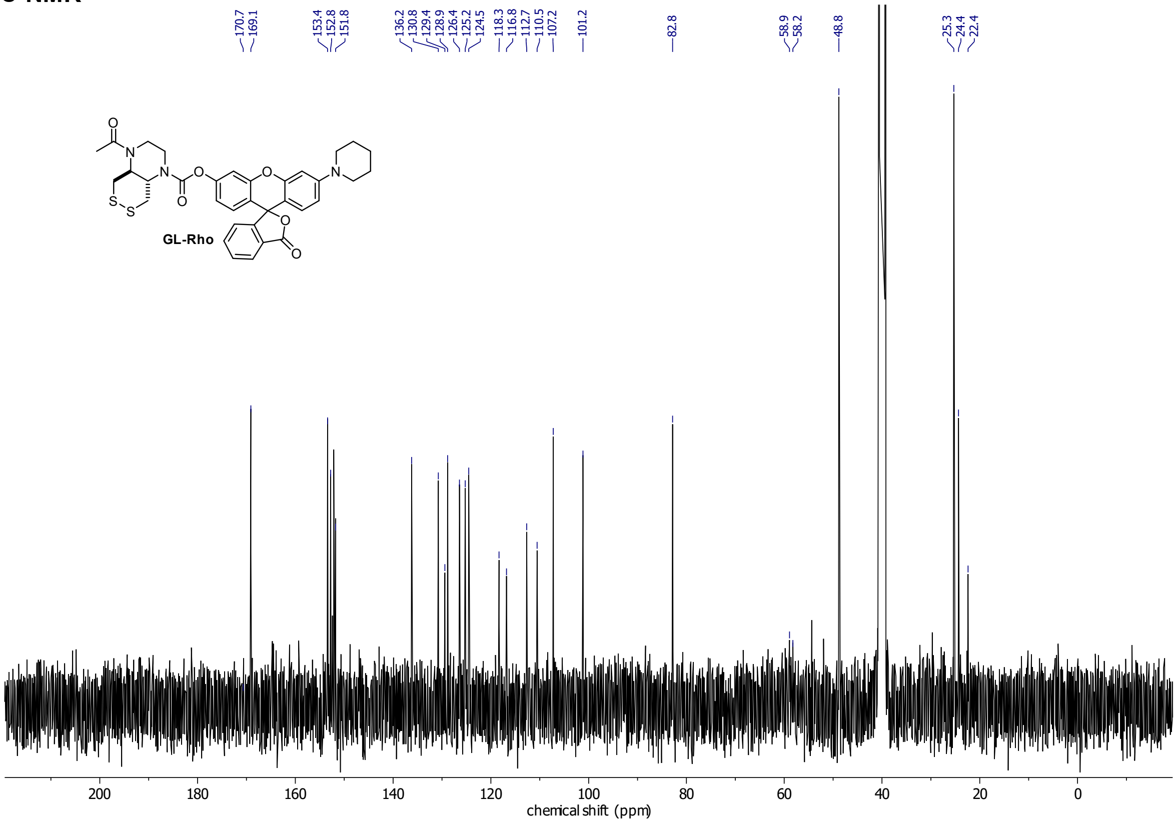

**H-Rho-A**
**$^{1}$H-NMR**

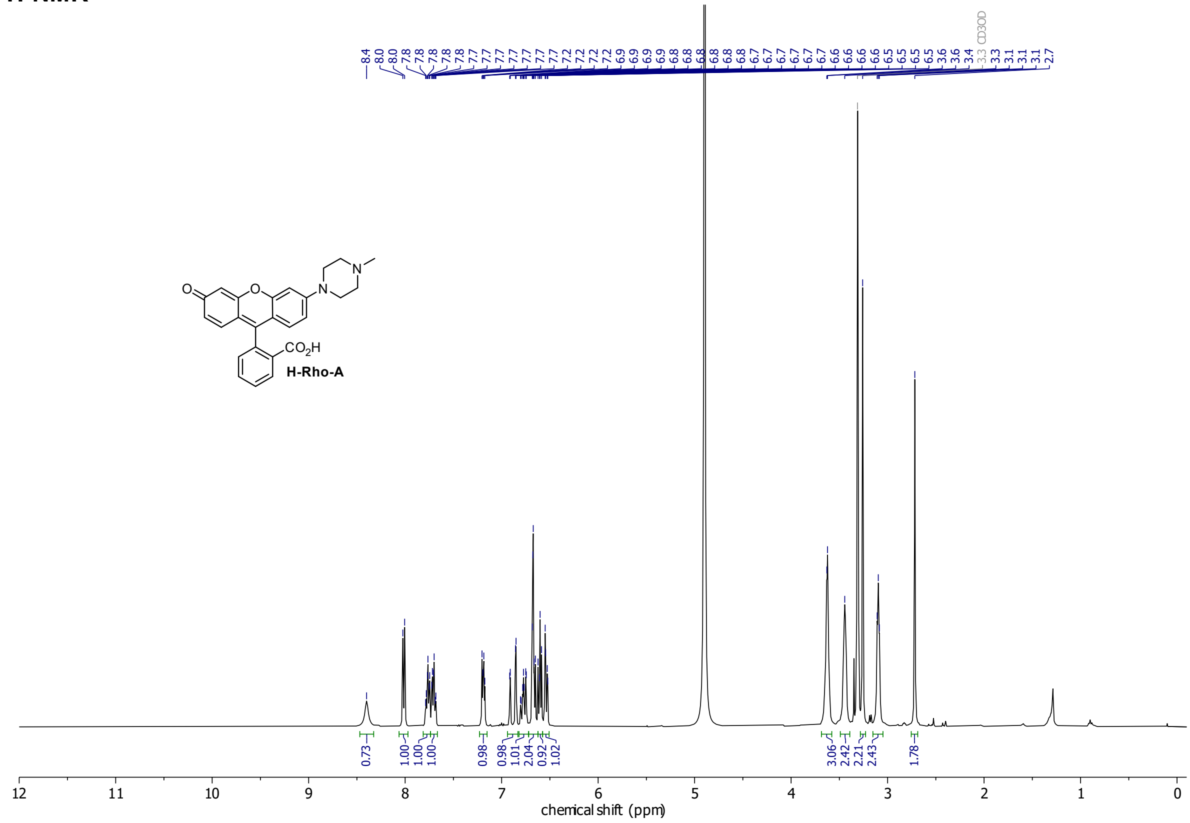

O
N
N
CO2H
H-Rho-A
0.73
1.00
1.00
1.00
0.98
0.98
1.01
2.04
0.92
1.02
3.06
2.42
2.21
2.43
1.78
12
11
10
9
8
7
6
5
4
3
2
1
0
chemical shift (ppm)

**GL-Rho-A**

**$^{1}$H-NMR**

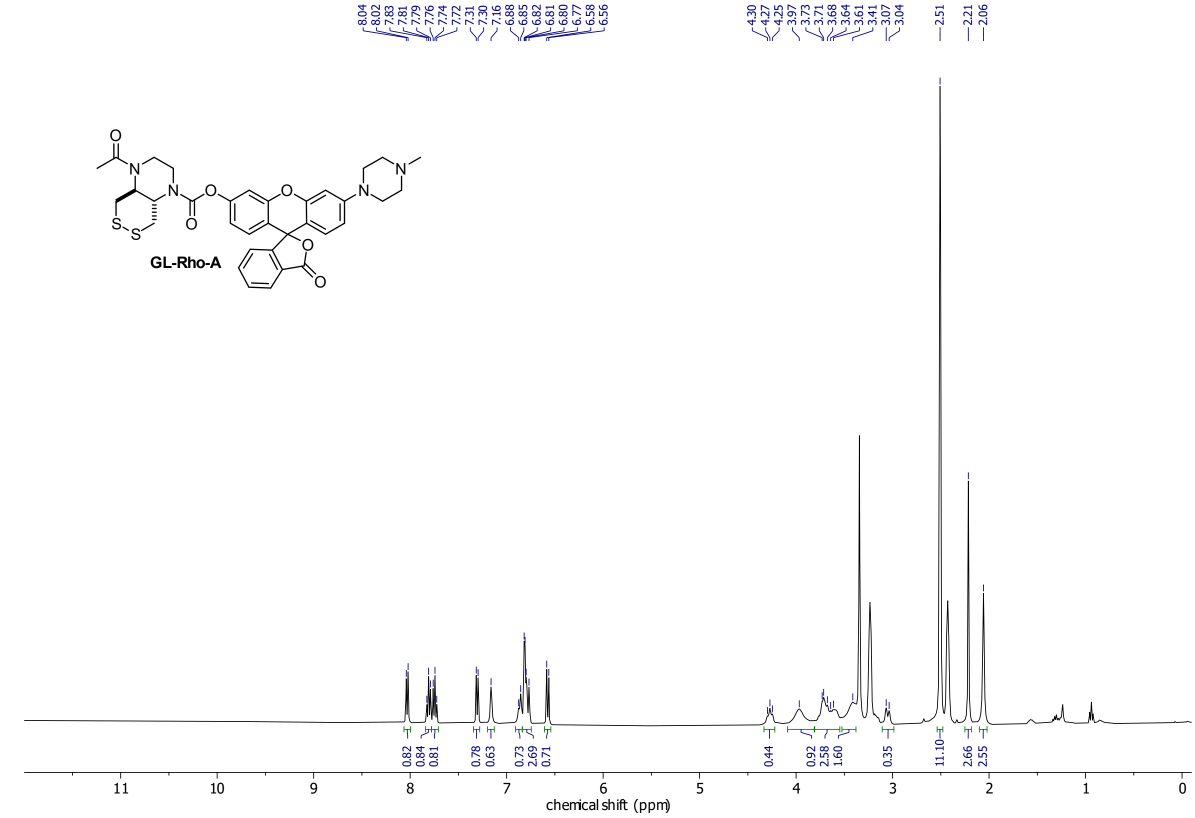

**$^{13}$C-NMR**

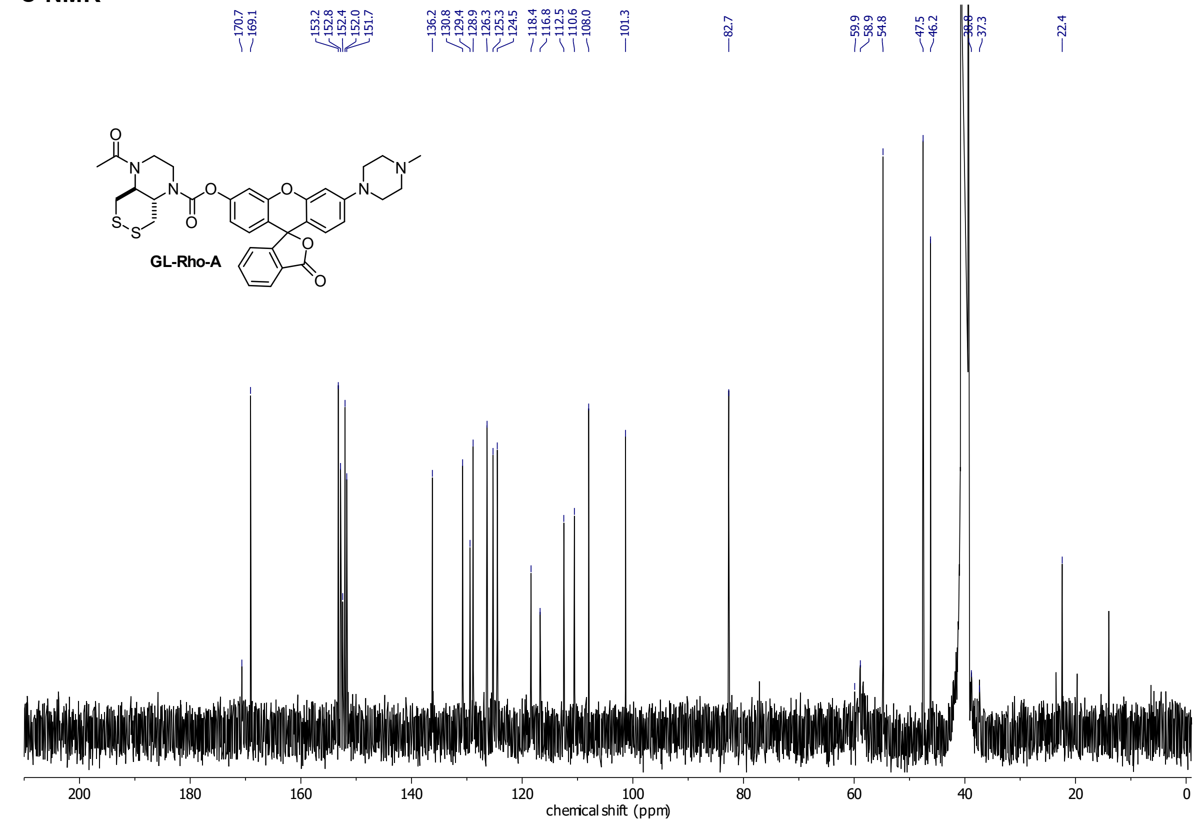

**Compound 12**
**$^1$H-NMR**

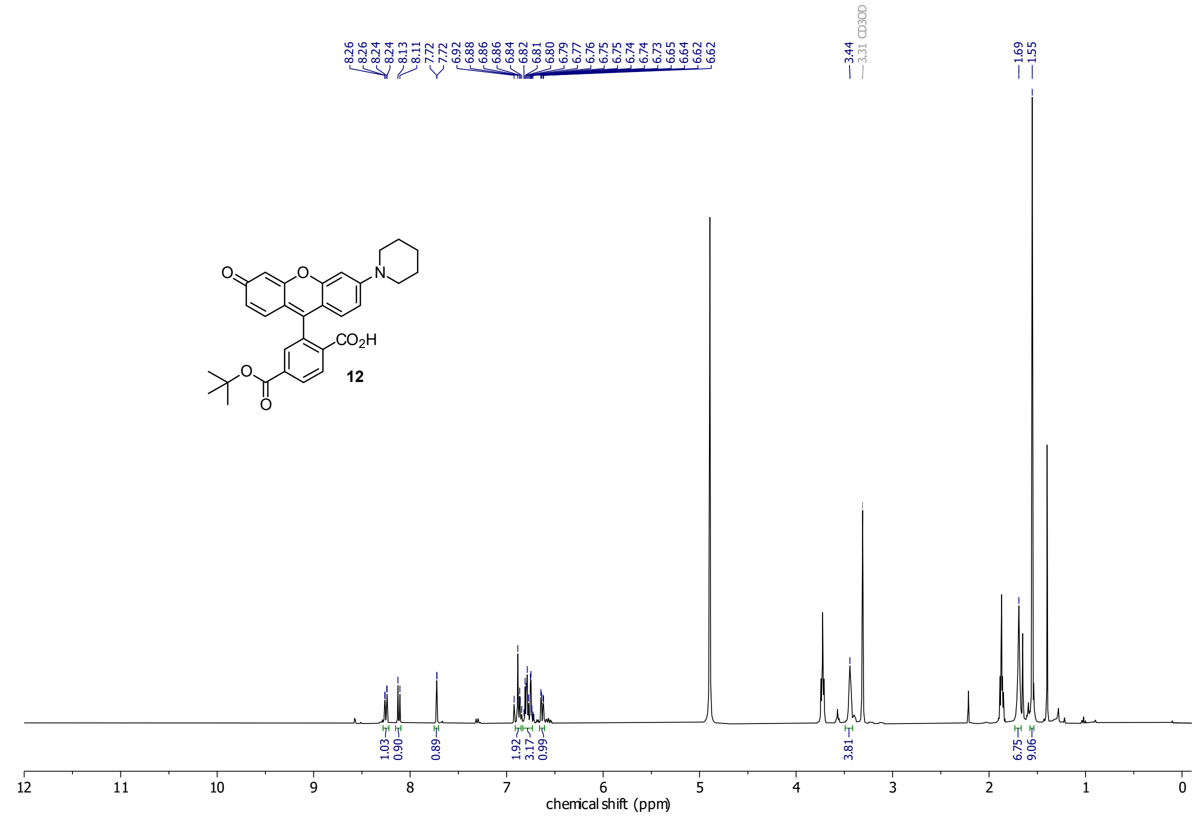

#### GL-Rho-C

##### $^{1}$H-NMR

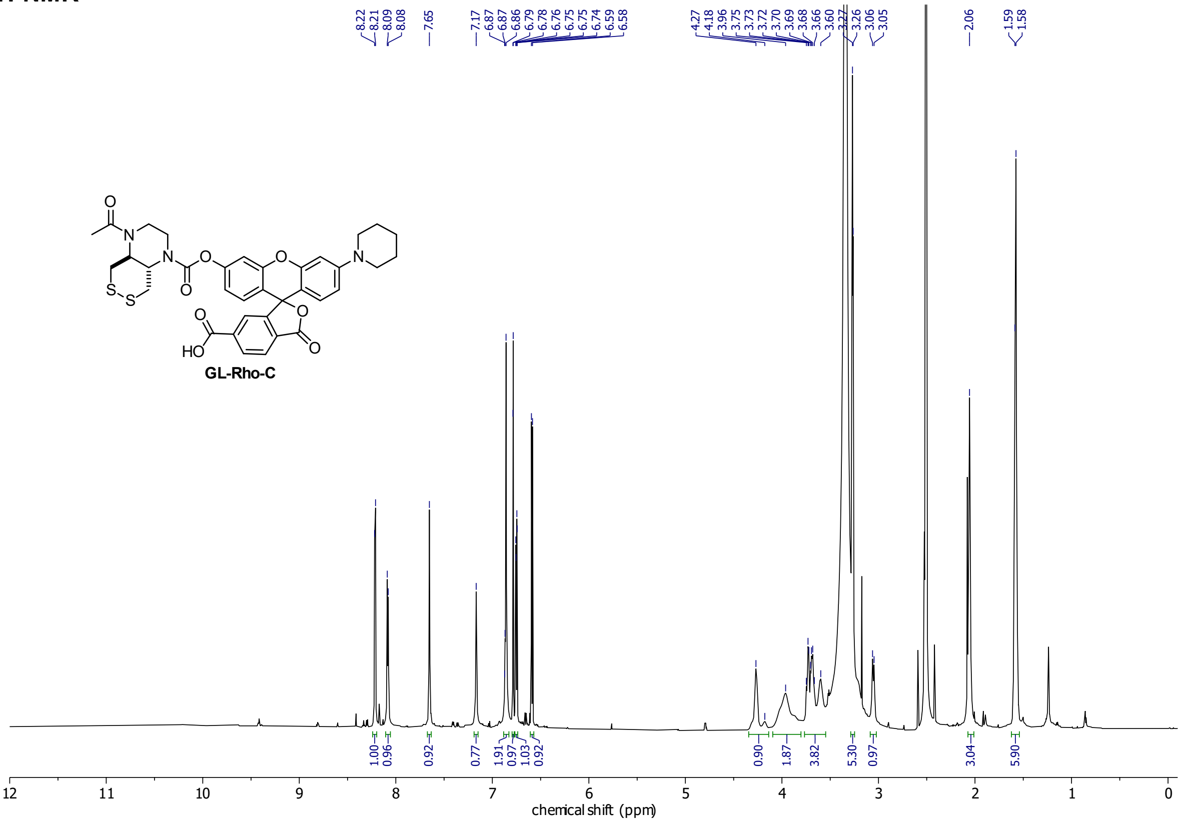

##### $^{13}$C-NMR

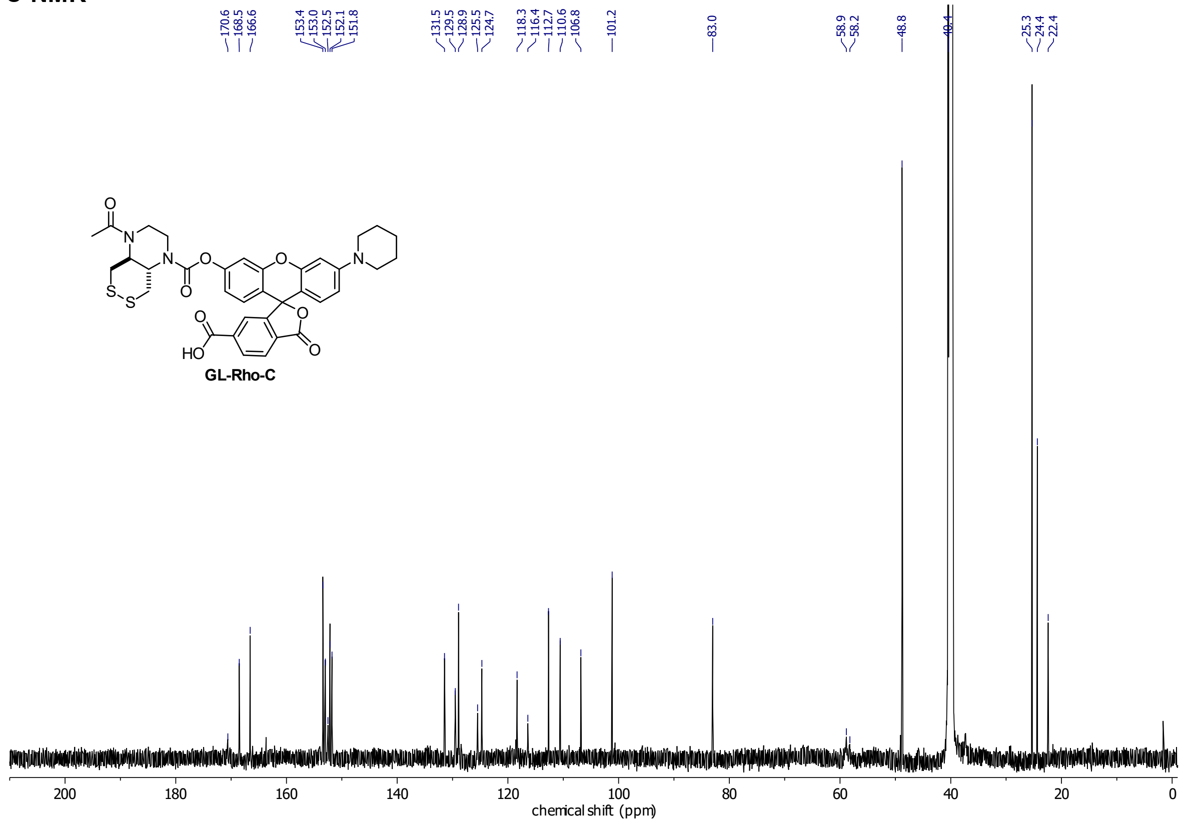

**GL-Rho-C^m**
**$^{1}$H-NMR**

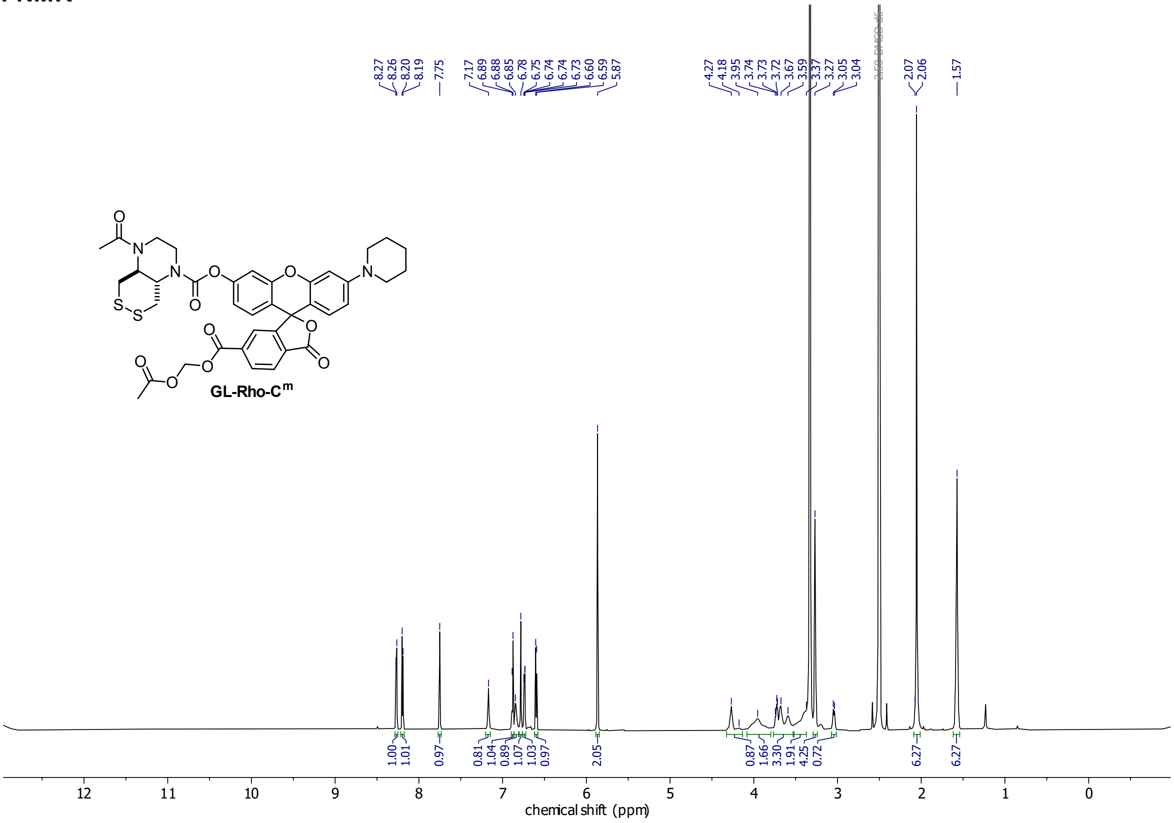

**$^{13}$C-NMR**

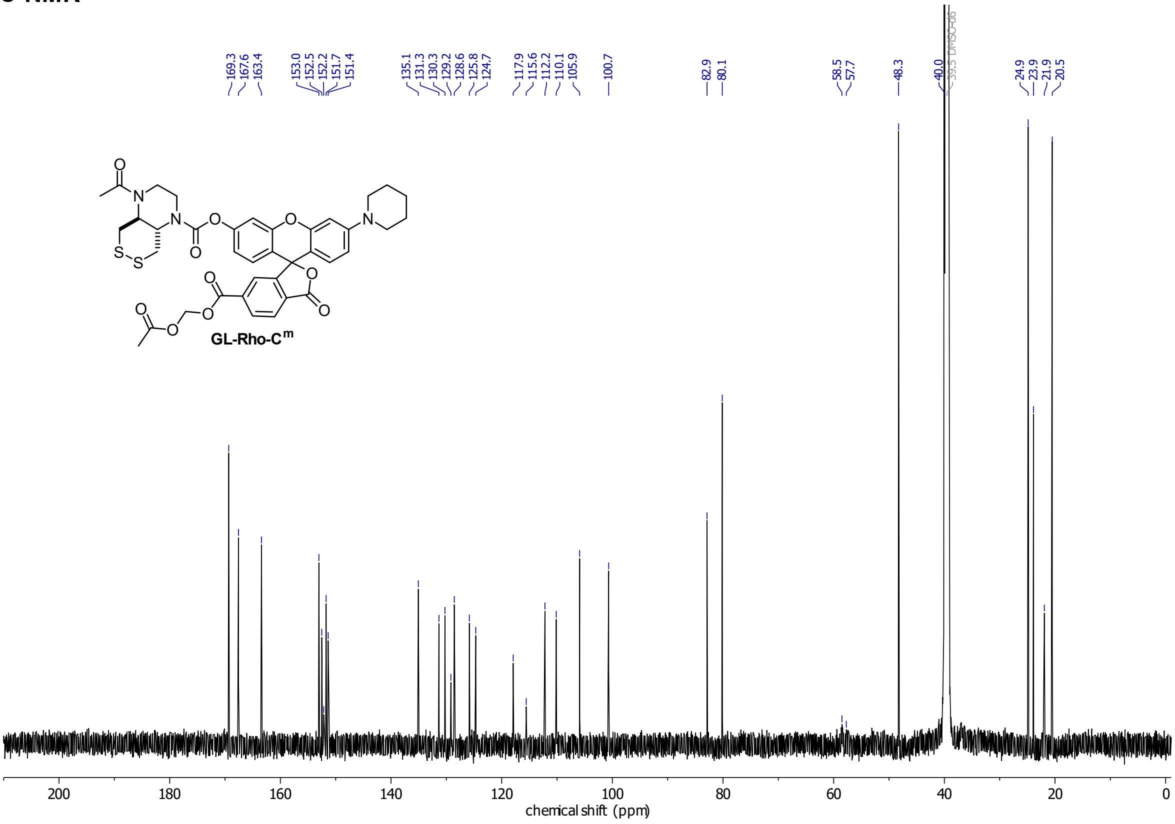

**H-Rho-C**

**$^{1}$H-NMR**

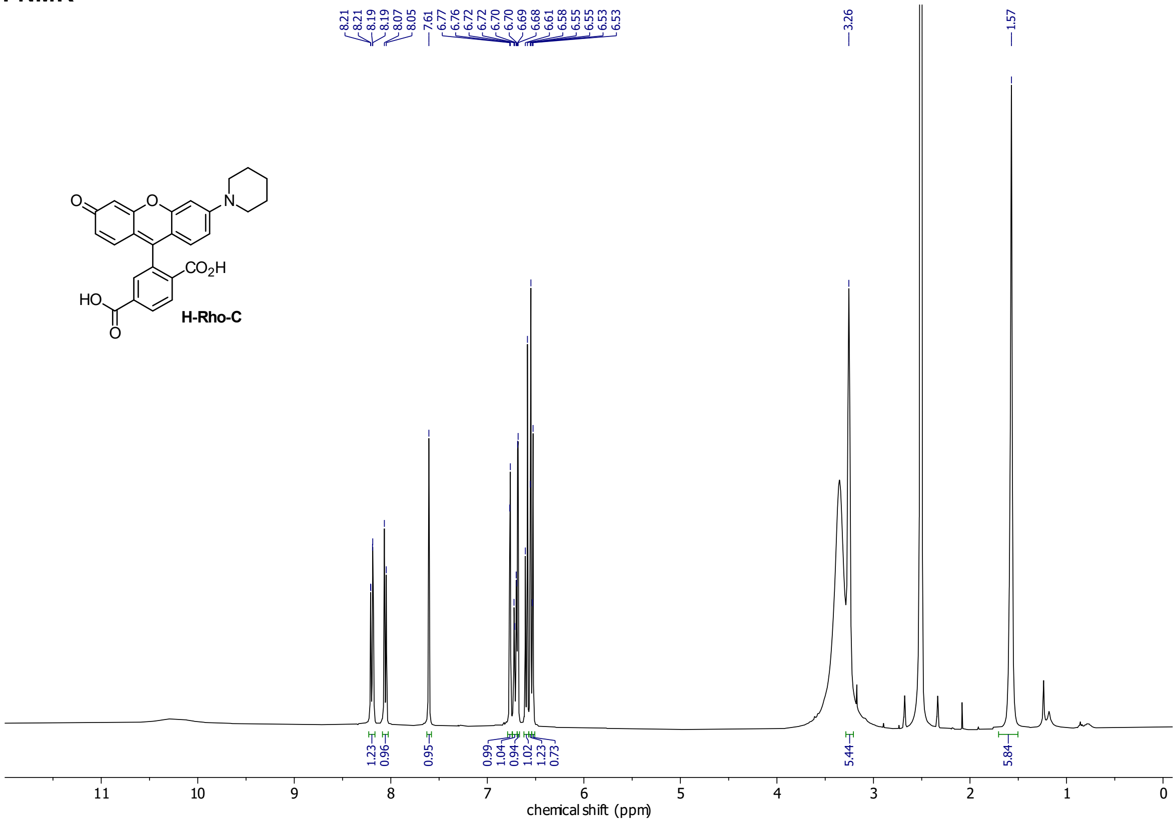

**$^{13}$C-NMR**

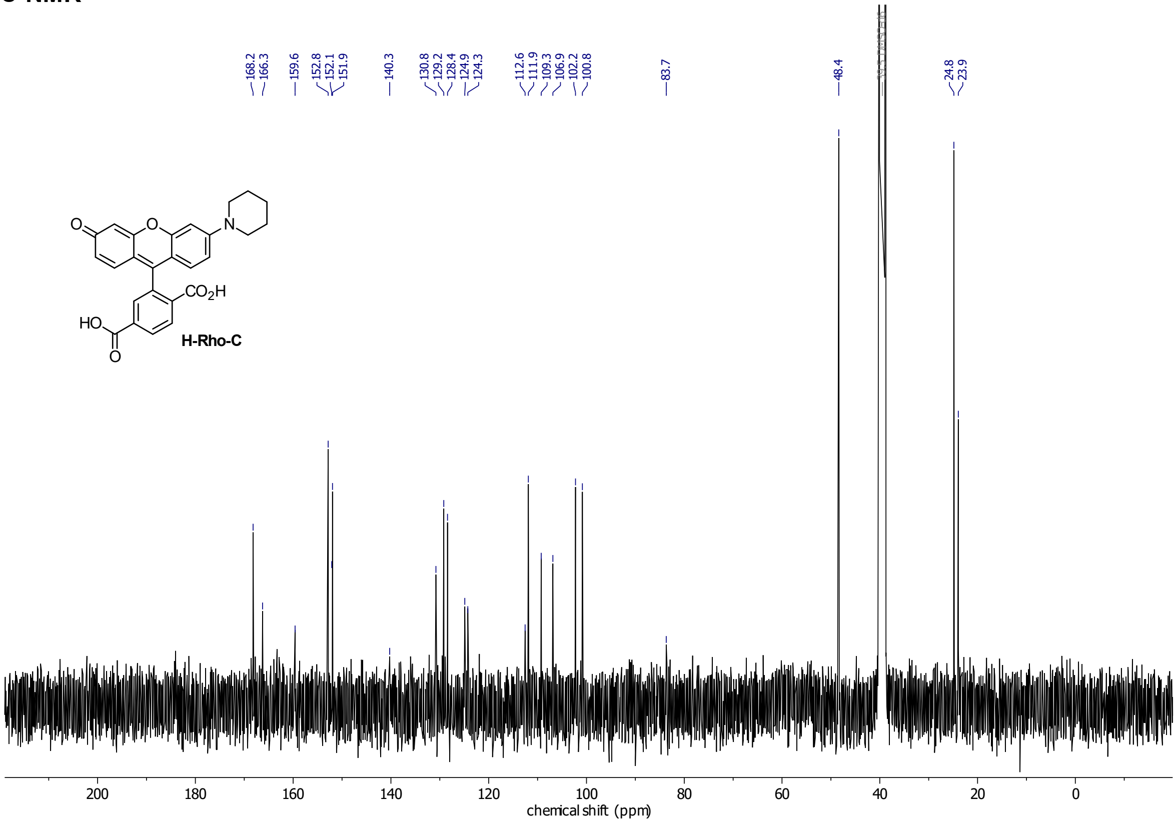

**Compound 16**
**$^{1}$H-NMR**

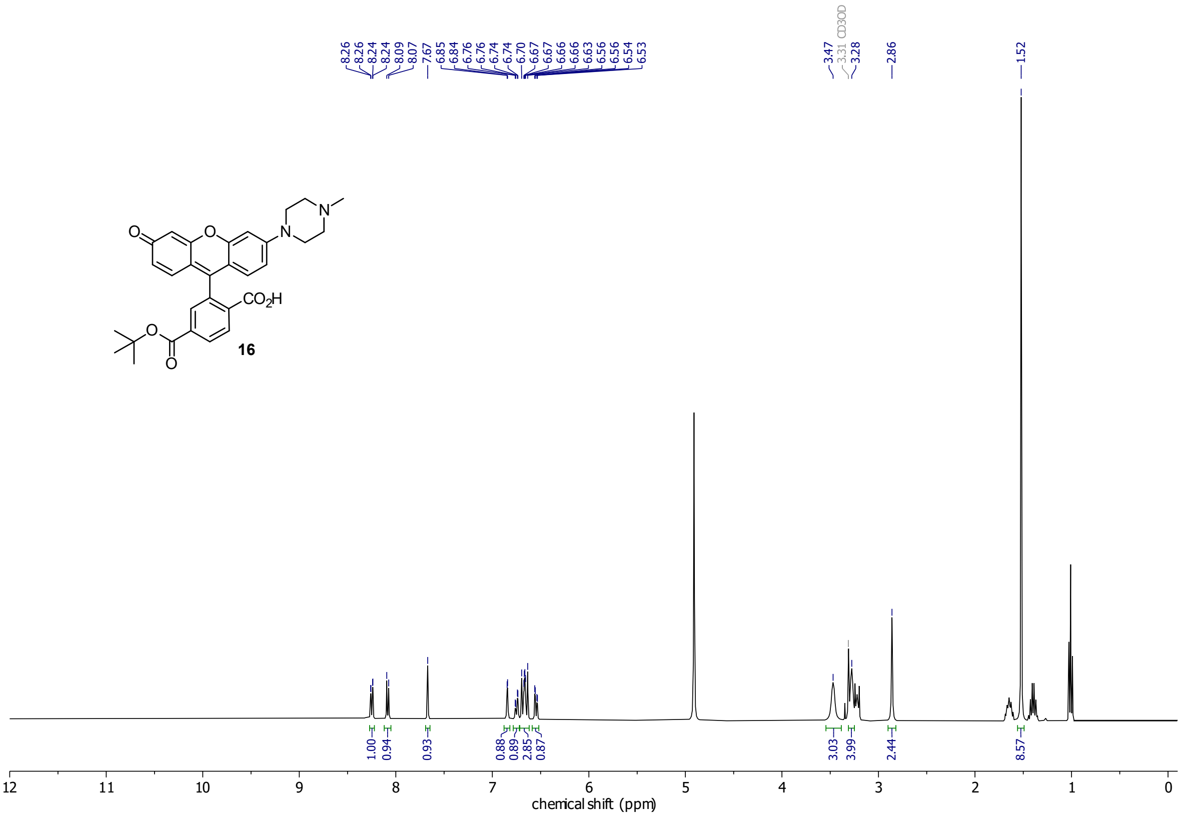

**GL-Rho-AC**

**1H-NMR**

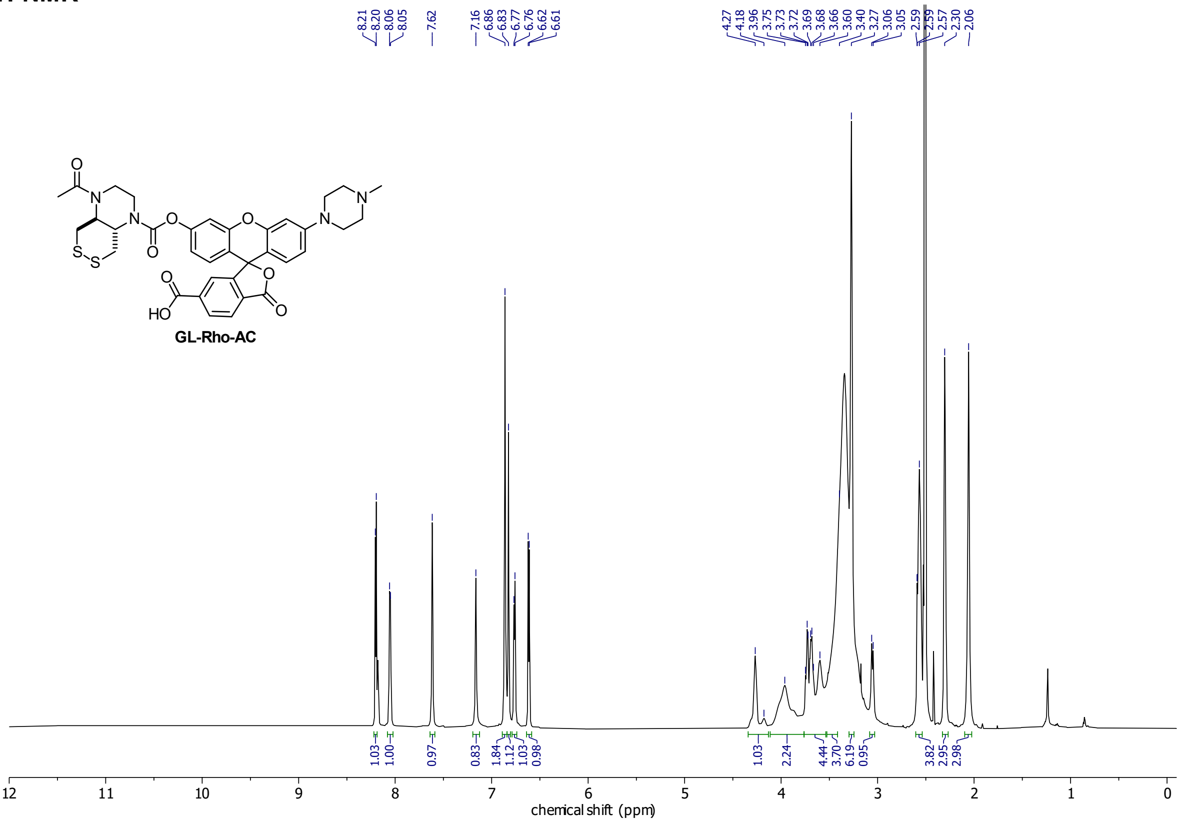

**13C-NMR**

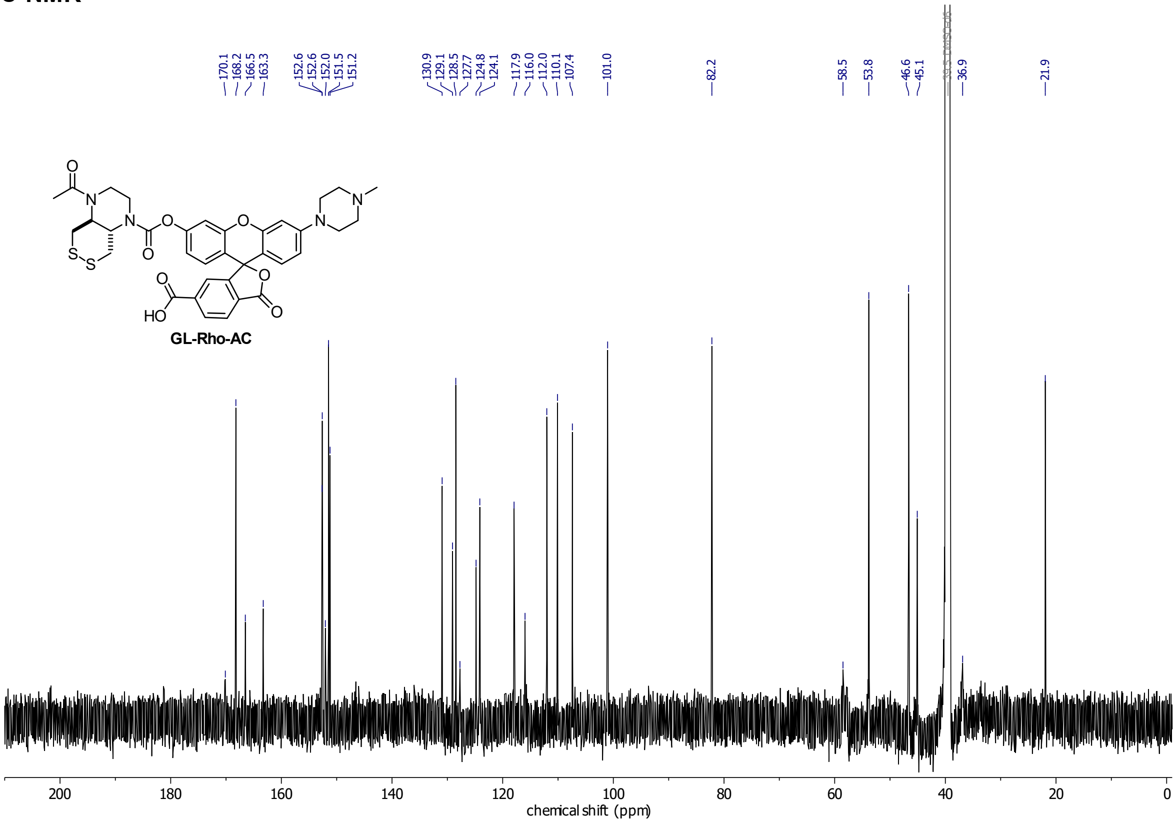

**GL-Rho-AC$^{m}$ (= GL-TraG)**
**$^{1}$H-NMR**

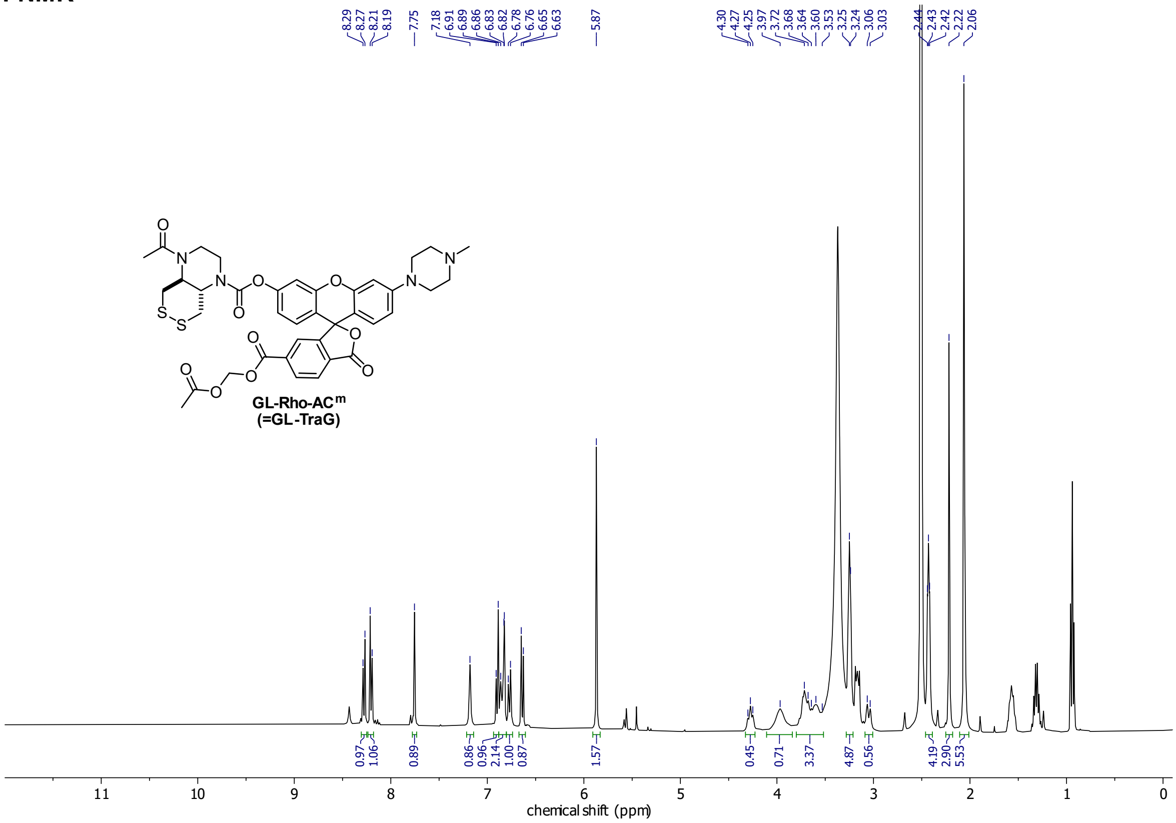

**H-Rho-AC (= H-TraG)**
**$^{1}$H-NMR**

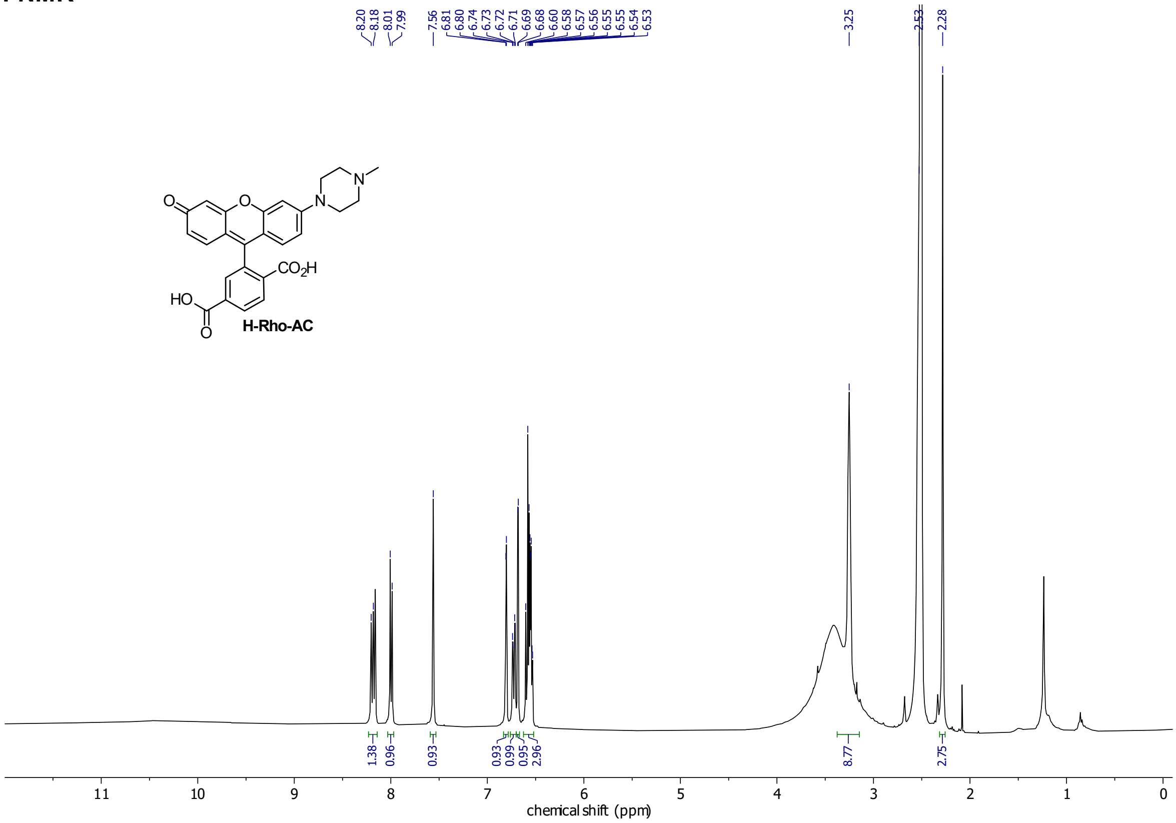

#### Compound 24

##### $^{1}$H-NMR

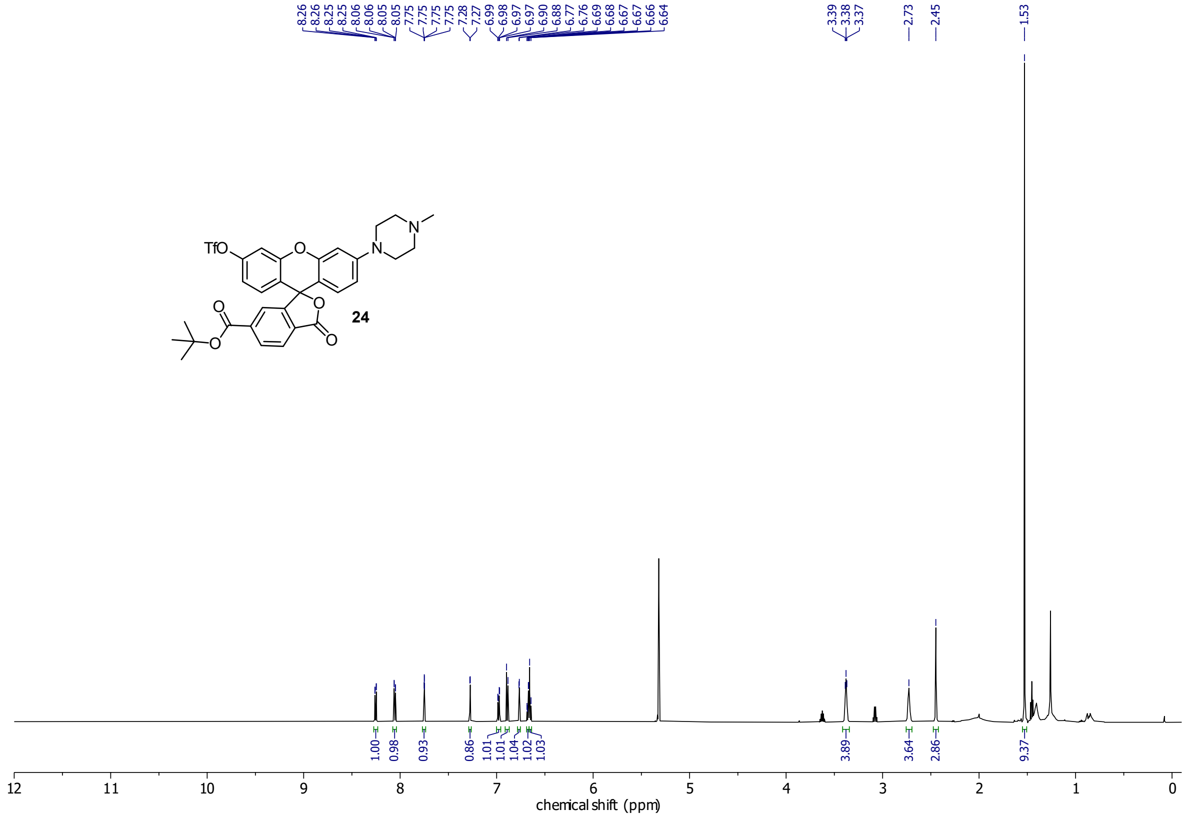

##### $^{13}$C-NMR

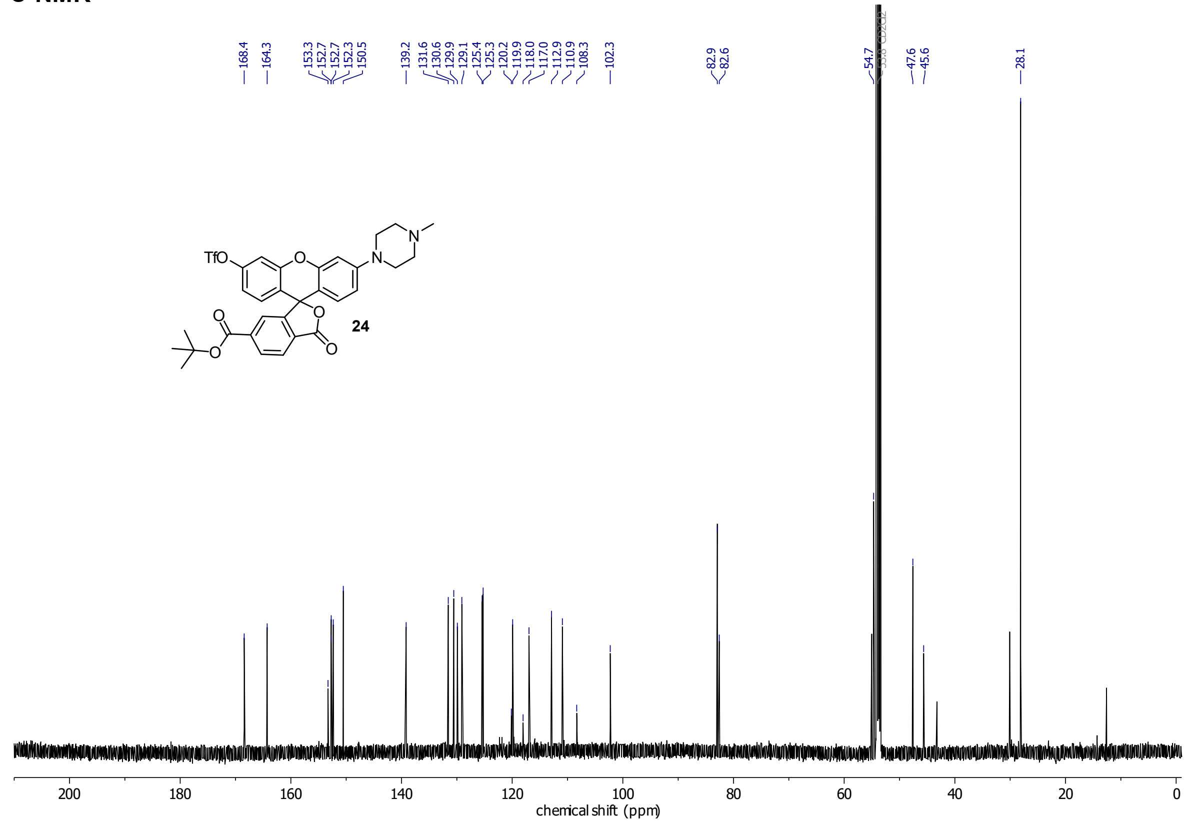

**HP-TraG**
**$^{1}$H-NMR**

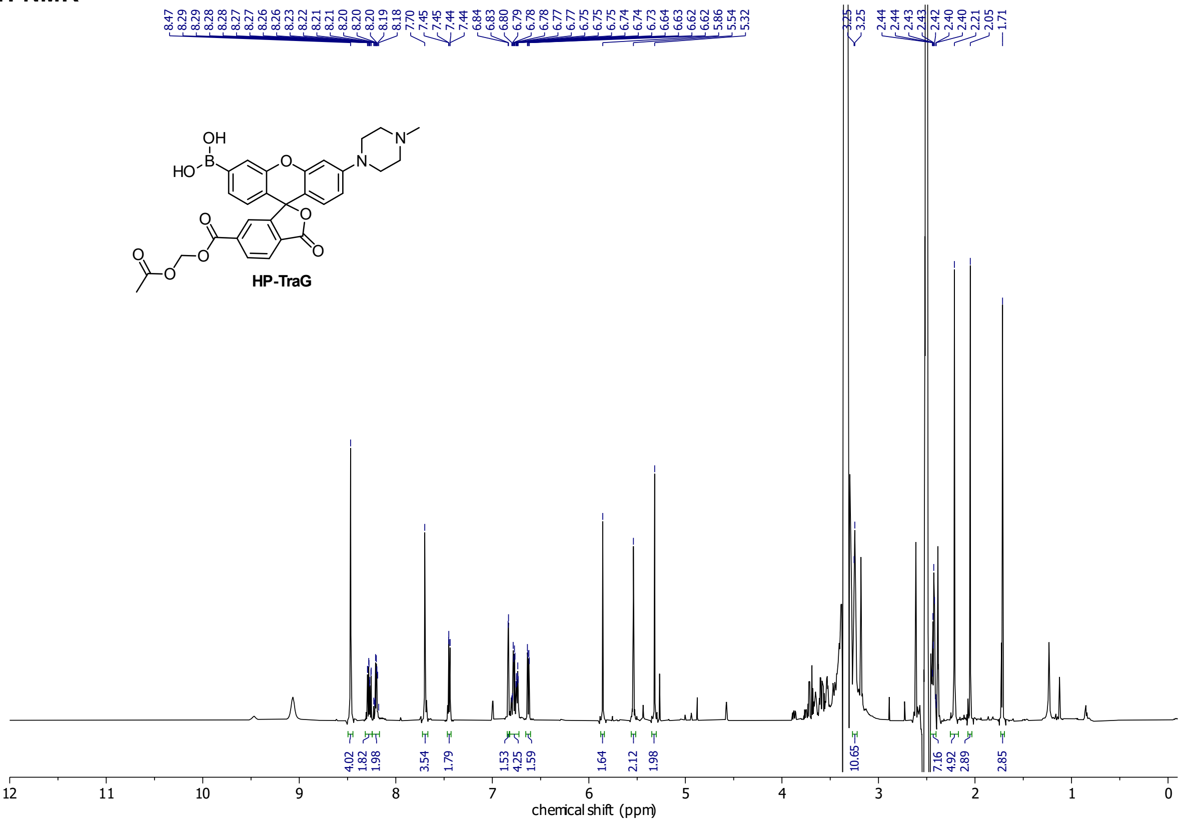

*contains other species (~50 mol%) with equal properties regarding esterase cleavage and hydrogen peroxide oxidation (see Figs26-27).

**TR-TraG**
**$^{1}$H-NMR**

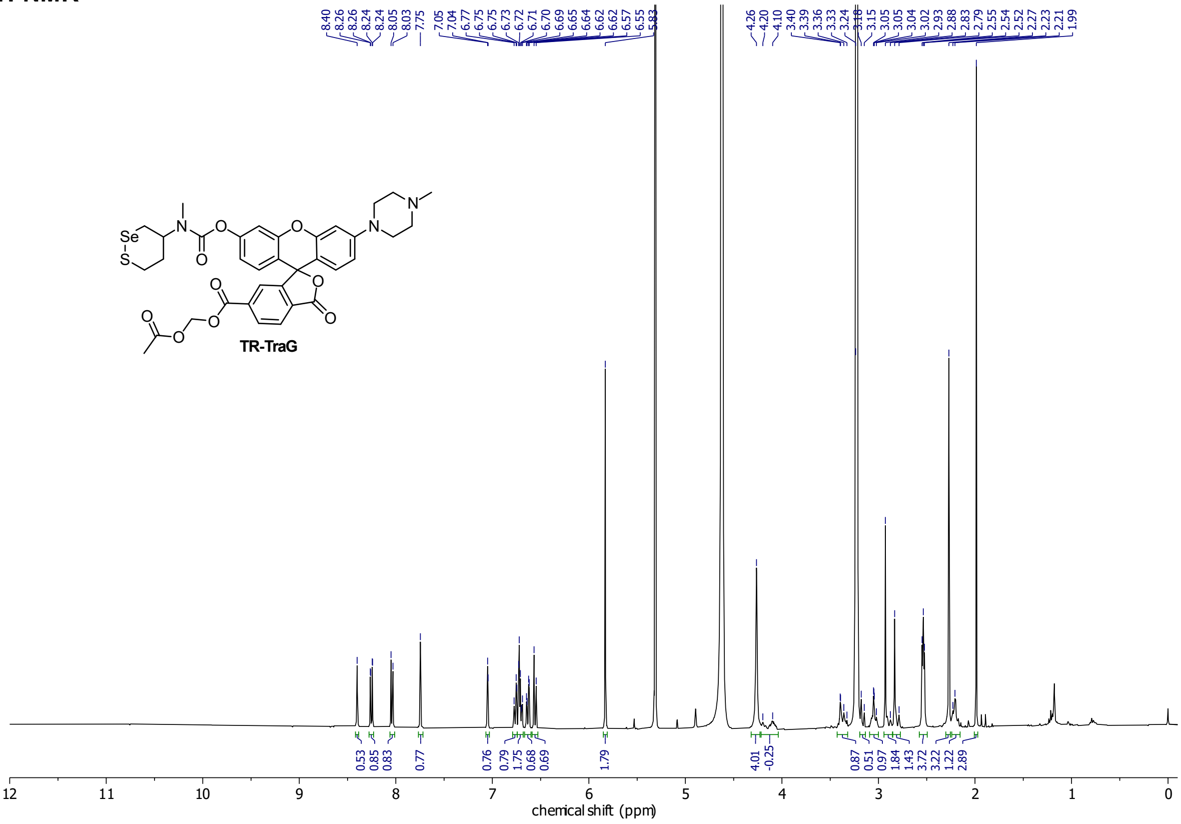
